# Accurate *de novo* design of high-affinity protein binding macrocycles using deep learning

**DOI:** 10.1101/2024.11.18.622547

**Authors:** Stephen A. Rettie, David Juergens, Victor Adebomi, Yensi Flores Bueso, Qinqin Zhao, Alexandria N. Leveille, Andi Liu, Asim K. Bera, Joana A. Wilms, Alina Üffing, Alex Kang, Evans Brackenbrough, Mila Lamb, Stacey R. Gerben, Analisa Murray, Paul M. Levine, Maika Schneider, Vibha Vasireddy, Sergey Ovchinnikov, Oliver H. Weiergräber, Dieter Willbold, Joshua A. Kritzer, Joseph D. Mougous, David Baker, Frank DiMaio, Gaurav Bhardwaj

## Abstract

The development of macrocyclic binders to therapeutic proteins typically relies on large-scale screening methods that are resource-intensive and provide little control over binding mode. Despite considerable progress in physics-based methods for peptide design and deep-learning methods for protein design, there are currently no robust approaches for *de novo* design of protein-binding macrocycles. Here, we introduce RFpeptides, a denoising diffusion-based pipeline for designing macrocyclic peptide binders against protein targets of interest. We test 20 or fewer designed macrocycles against each of four diverse proteins and obtain medium to high-affinity binders against all selected targets. Designs against MCL1 and MDM2 demonstrate K_D_ between 1-10 μM, and the best anti-GABARAP macrocycle binds with a K_D_ of 6 nM and a sub-nanomolar IC_50_ *in vitro*. For one of the targets, RbtA, we obtain a high-affinity binder with K_D_ < 10 nM despite starting from the target sequence alone due to the lack of an experimentally determined target structure. X-ray structures determined for macrocycle-bound MCL1, GABARAP, and RbtA complexes match very closely with the computational design models, with three out of the four structures demonstrating Ca RMSD of less than 1.5 Å to the design models. In contrast to library screening approaches for which determining binding mode can be a major bottleneck, the binding modes of RFpeptides-generated macrocycles are known by design, which should greatly facilitate downstream optimization. RFpeptides thus provides a powerful framework for rapid and custom design of macrocyclic peptides for diagnostic and therapeutic applications.

## MAIN TEXT

Macrocyclic peptides present a promising avenue for developing new therapeutics that bridge the gap between small-molecule drugs and large biologics^1,2^. Biologics, while capable of binding diverse therapeutic targets with high affinity and selectivity, are usually unable to cross cell membranes due to their large size and high polarity, limiting them to extracellular targets. Conversely, small molecules can access intracellular targets but are not ideal for targeting proteins lacking deep hydrophobic pockets. In principle, macrocyclic peptides with sizes between small molecules and proteins can be developed to modulate molecular targets inaccessible to traditional therapeutic modalities^3^. The ability to develop custom protein-binding macrocycles for diverse protein targets would have many diagnostic and therapeutic applications. Traditionally, the development of peptide therapeutics has relied on natural product discovery or high-throughput screening of trillions of random peptides for target binding using display-based techniques^1,2^. However, natural product discovery has several challenges, particularly synthetic difficulties, marginal stability, and low mutational tolerance of identified hits^4^. While powerful, the high-throughput screening methods are time-, cost-, and labor-intensive, and only span a small fraction of the rich chemical and structural diversity accessible to macrocycles. Moreover, such approaches frequently fail to simultaneously optimize for multiple biophysical properties, such as target binding, selectivity, and membrane permeability, due to the precise structural control required to achieve such functional properties^5^.

Structure-guided design methods offer a complementary approach to the library screening approaches, enabling rapid *in silico* exploration of a large chemical and structural diversity to design macrocycle binders for therapeutic targets. We previously developed physics-based methods for designing hyperstable constrained peptides, structured macrocycles, and binders to protein targets by borrowing the motifs or interactions from previously described binding partners as anchors^6–9^. However, despite the high accuracy observed in the design of monomeric macrocycles with these methods^7^, the design of protein-binding macrocycles has had limited success, achieving only modest binding affinities and, in many cases, with the experimentally determined structures not agreeing with the design models^7,8,10^. The reliance on previously described binding partners for starting motifs also restricts such approaches to well-studied protein targets. In recent work, we described a pipeline for hallucinating and predicting the structures of macrocyclic peptide monomers by modifying AlphaFold2 (AF2) to include cyclic relative positional encoding (named “AfCycDesign”)^11^. Other promising deep learning (DL) methods have been described recently to predict the structures of macrocycles and macrocycle-target complexes^12,13^ and to design peptide binders to protein targets^14–16^. However, these methods have not been extensively structurally validated to date or shown to robustly perform atomically accurate *de novo* design of macrocyclic peptide structures in complexes with diverse protein targets. Computational methods that can accurately design high-affinity macrocycle binders *de novo*, using just the information of target structure or sequence, are required for wider therapeutic applications.

We reasoned that recent breakthroughs in generative DL methods could be leveraged to develop a robust pipeline for the accurate and efficient design of macrocycle binders. Diffusion models for protein design, such as RFdiffusion^17^, are trained to generate diverse protein structures from randomly initialized residues as starting points and have demonstrated remarkable success in designing protein monomers, binders, and symmetric oligomers of medium to large-sized proteins. However, despite considerable recent progress in DL-based protein design methods, these methods do not readily apply to designing macrocyclic peptides. Developing analogous methods for peptide design from scratch has been challenging due to limited availability of experimental data for training such models. To address these challenges, we set out to extend the RoseTTAFold2^18^ (RF2) structure prediction network and the RFdiffusion^17^ protein backbone generation framework to incorporate cyclic relative positional encoding and enable the generation of the macrocyclic peptide backbones.

### Extending RoseTTAFold2 and RFdiffusion for macrocyclic peptides

We began by examining the ability of the RF2^18^ structure prediction network to model known macrocyclic peptide structures. We implemented a modified (see Methods) cyclic relative position encoding for RF2 (Figure 1A) and observed robust prediction of natural cyclic peptide structures (Figure S1). Given this success, we reasoned that the same relative position encoding should enable RFdiffusion^17^ to generate macrocyclic peptide structures due to its similar network architecture. We added the cyclic positional encoding scheme to RFdiffusion and observed robust generation of diverse macrocyclic peptides (Figure 1B-C, S2). Encouraged by the transferability of the cyclic positional encoding, we set out to employ RFdiffusion for the *de novo* design of protein-binding macrocycles. We chose RFdiffusion for several reasons: First, we expected the high experimental success rate of RFdiffusion^17,19^ for protein binder design to carry over to macrocycle binder design. Second, *de novo* binder design with AfCycDesign as-is would be far more computationally expensive and has not been successfully implemented or experimentally validated. Third, the method can still take advantage of the current built-in conditional generation functionalities of RFdiffusion, such as epitope-specific targeting and “motif” scaffolding. Finally, the method should be directly transferable to other current and future RoseTTAFold-based design networks, such as RFdiffusion All-Atom^20^, for incorporating non-peptidic molecules (nucleic acids, ions, etc.) during design calculations.

**Figure 1:**
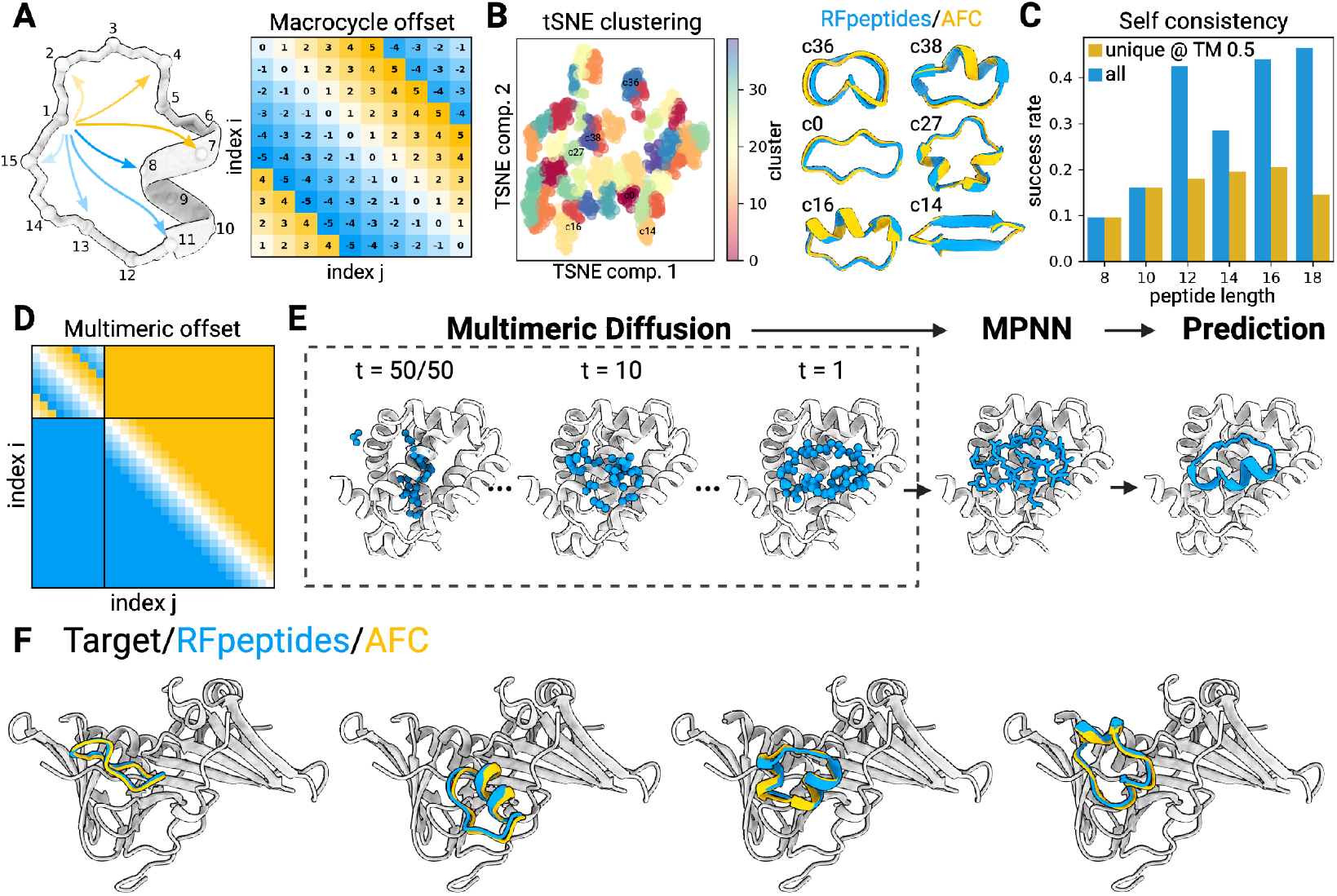
RFpeptides is a diffusion-based pipeline for the *de novo* design of protein-binding macrocycles. (A) Introduction of a cyclically symmetric relative position encoding to RFdiffusion enables the generation of macrocyclic peptide backbones with N- and C-termini covalently linked via a peptide bond. The relative position encodings are cyclized by switching from positive relative position encodings (i.e., “to the right”) to negative encodings (i.e., “to the left”) when index j is more than halfway around the peptide relative to index i. (B) RFpeptides produces diverse and designable cyclic peptides traversing α-helical, β-strand, and loop/coil secondary structures. Left, Structural clusters calculated using t-SNE^32^ to reduce the dimensionality of an all-by-all TMscore matrix computed with TMalign^33^ on an unfiltered set of 1200 macrocycles generated using RFpeptides. Clustering of the 1200 (now 2D) points was then performed using a 40-component Gaussian mixture model in Scikit-learn^34^. Right, Six RFpeptides outputs from differing structural clusters, all with < 1 Å backbone RMSD between the design model (blue) and the structure predicted by AfCycDesign (gold). (C) Self-consistency benchmark results for RFpeptides cyclic peptide design at various lengths. For each peptide length, the fraction (with N=200 per length) of backbones with at least one out of the of eight LigandMPNN^35^ sequences predicted by AfCycDesign to refold with pLDDT > 0.8 and within 2.0 Å backbone RMSD of the designed structure. Success rates for all sampled backbones are in blue, and success rates only counting unique structural clusters (as calculated using MaxCluster^36,37^ at a TMscore threshold of 0.5) are in orange. (D) For multi-chain diffusion trajectories (e.g., macrocycle binder design), the relative position encodings for the macrocycle chain are cyclized, whereas inter-chain and target chain relative positional encoding are kept as standard. (E) Pipeline for the design of protein-binding macrocycles using RFpeptides. Macrocycle backbones are generated from randomly initialized atoms by a stepwise RFdiffusion-based denoising process, followed by fixed-backbone amino acid sequence design using ProteinMPNN. Design models are downselected based on the computational metrics from structure prediction using AfCycDesign and physics-based interface quality metrics using Rosetta. (F) RFpeptides pipeline generates diverse macrocycles against selected targets. Four diverse cyclic peptide binders against the same target were generated using RFpeptides, with AfCycDesign iPAE < 0.3 and < 1.5 Å Cα RMSD between the design model (blue) and AfCycDesign prediction (gold).

We modified the RFdiffusion protein binder design pipeline to use cyclic relative position encodings for the generated chain, and standard positional encodings for the target and inter binder-target indices (Figure 1D). We then completed our design pipeline by using ProteinMPNN to design amino acid sequences compatible with the backbones generated by RFdiffusion (Figure 1E). This pipeline readily generated macrocycles with diverse secondary structure content against target proteins (Figure 1F), and the inclusion of standard RFdiffusion hotspot features clearly shifted the distribution of generated binders toward desired residues (Figure S3). We refer to this integrated pipeline as “RFpeptides” throughout the remainder of the text.

### *De novo* design and characterization of macrocyclic binders to MCL1 and MDM2

We selected myeloid cell leukemia-1 (MCL1) as our first target protein, given the availability of multiple high-resolution X-ray crystal structures available to initiate the design calculations. MCL1 is also a promising target for anti-cancer therapeutics due to its roles in autophagy, cell survival, DNA repair, and cellular proliferation^21^. For targeting MCL1, we used RFpeptides to generate 9,965 diverse cyclic peptide backbones, followed by four iterative rounds of ProteinMPNN and Rosetta Relax to design four amino acid sequences for each generated backbone. We expected the local changes to the generated backbone during the Rosetta Relax steps to allow for improved amino acid sequence diversity from the MPNN steps. While there are other ways to achieve increased sequence diversity, including generating multiple sequences per backbone from ProteinMPNN or adding noise during MPNN sequence generation, we did not explicitly try or compare them in this study. For downselecting the design candidates for experimental testing, we used AfCycDesign to re-predict the designed macrocycle-target complexes from the macrocycle sequence and the target structure as a template. We selected the designs based on the confidence metric (interface predicted aligned error (iPAE)) and the similarity between the original design model and the protein-macrocycle complex predicted by the AfCycDesign (Figure S4). For further stringency in the design selection process, we also used RF2 to re-predict the complex structures, reasoning that the design models predicted identically by two orthogonal structure prediction networks (AfCycDesign and RF2) should have a higher likelihood of binding to the target as designed. However, the 1,984 selected designs at this stage were still more than the number of designs we could reasonably synthesize and test experimentally. Therefore, we next used Rosetta^22^ to calculate the ‘physics-based’ metrics of interface and macrocycle quality, such as calculated binding affinity (ddG), spatial aggregation propensity (SAP) of the designed macrocycle, and the molecular surface area of the interface contacts (CMS) (Figure S4).

After strictly filtering the designed candidates on DL-based and physics-based metrics, we selected 27 designs for synthesis, biochemical, and biophysical characterization. Despite specifying no hotspots to guide the generation process to a specific patch on the MCL1 structure, all selected designs bound to the functionally-relevant MCL1-BIM interaction site (Figure S5). While all selected designs include an α-helical segment, they feature different sequences, macrocycle placement, and target interactions (Figure S5 and Table S1). In addition to the common helical motifs, the loop regions of the selected macrocycles also contribute significant side chain and backbone-mediated interactions to the binding interface. During the chemical synthesis using the Fmoc-based solid phase synthesis (see Methods), the yields for the correctly cyclized product for 13 designs were low and insufficient for further characterization. We tested the remaining 14 macrocycles for binding to biotinylated MCL1 using surface plasmon resonance (SPR) single-cycle kinetics experiments (Figure S6). Three macrocycles showed binding to the MCL1, with the best binder, MCB_D2 (refers to *<u>MC</u>L1 <u>B</u>inding <u>D</u>esign 2*), (Figure 2A), demonstrating a binding affinity of 2 μM (Figure 2B). To confirm if the designed macrocycle adopts the designed structure and engages MCL1 in the designed binding mode, we determined the X-ray crystal structure of MCB_D2 bound to MCL1 at 2.1 Å resolution. The crystal structure is nearly identical to the design model, with an RMSD of 0.7 Å over all the Cα atoms of the macrocycle with target chains aligned (Figure 2C) and Cα RMSD of 0.4 Å within the macrocycles when aligned (Figure 2D). The side chain rotamers of the interacting residues in the crystal structure also closely match the design model (Figure 2D). The crystal structure also confirms that the binding interactions are not restricted to the helix region of the designed macrocycle but are also contributed by the loop regions (Figure 2E-F).

**Figure 2:**
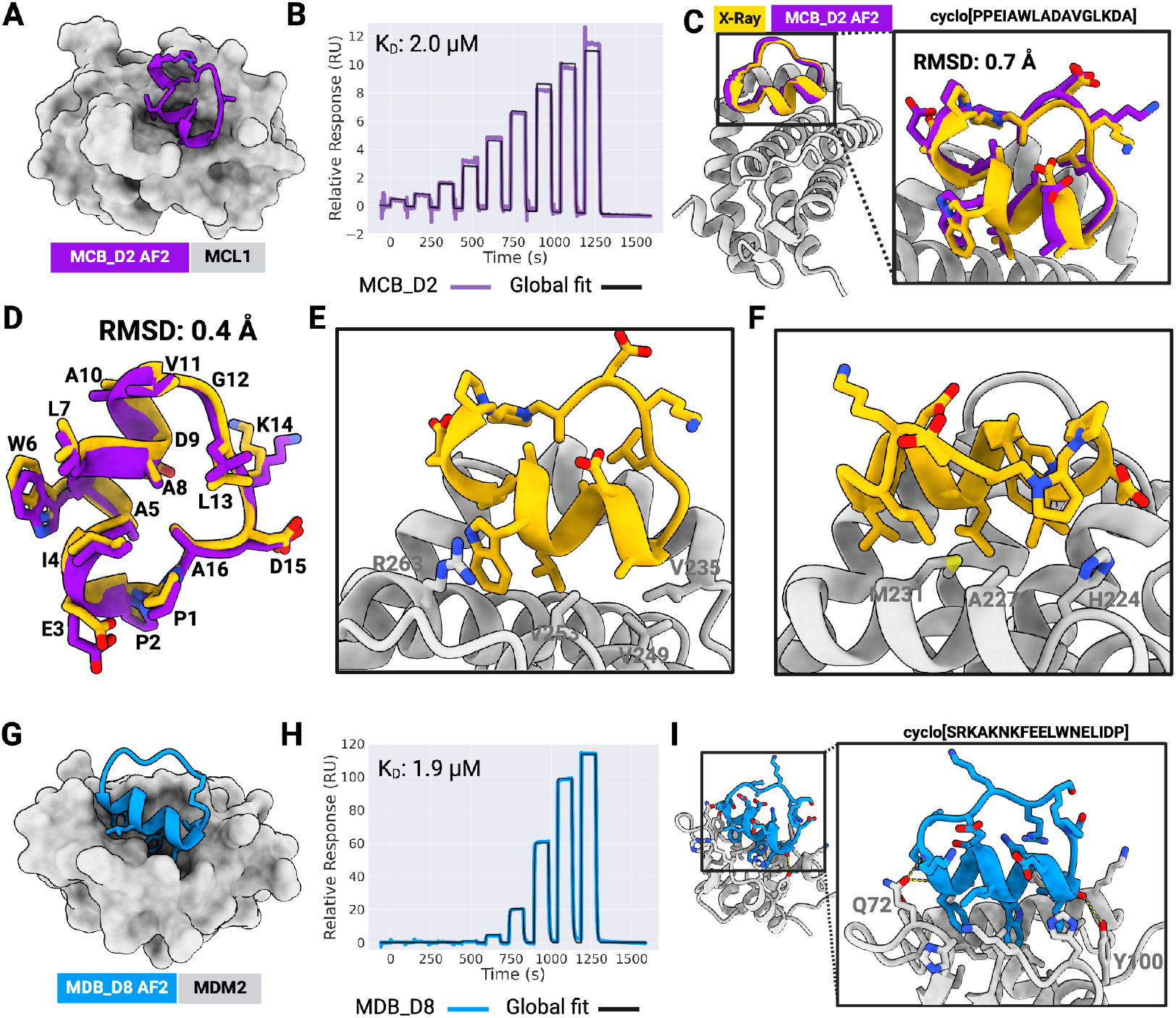
*De novo* design and characterization of macrocyclic binders to MCL1 and MDM2. (A) AfCycDesign prediction of MCB_D2 (purple) bound to MCL1 (gray surface). MCB_D2 side chains are shown as sticks. (B) Affinity determination of MCB_D2 using SPR. SPR sensorgram from a 9-point single-cycle kinetics experiment (2-fold dilution, highest concentration: 20 μM). Experimental data are shown in purple and global fits are shown with black lines. The dissociation constant (K_D_) is also shown on the plot. (C) Experimentally determined complex structures closely match the design model. Overlap of the X-ray crystal structure (gold and gray) with the design model for MCB_D2 (purple); Cα RMSD for macrocycle is 0.7 Å when experimental structure and design models are aligned by MCL1 residues. Close-up views demonstrate strong agreement between the side chain rotamers of the design model and the X-ray structure. (D) Overlay of the macrocycle model to the crystal structure shows a Cα RMSD of 0.4 Å with nearly identical backbones and side chain rotamers. (E) Close-up view of the macrocycle-bound MCL1 structure showing the cation-π interaction at the interface. (F) Close-up view of the macrocycle-bound MCL1 structure showing the hydrophobic contacts at the interface. (G) AfCycDesign prediction of MDB_D8 design (blue) in complex with MDM2 (gray) shown as cartoons with interacting side chains shown as sticks. (H) Affinity determination of MDB_D8 using SPR. SPR sensorgram from a 9-point single cycle kinetics experiment (5-fold dilution, highest concentration: 50 μM). Experimental data are shown in blue and global fits are shown with black lines. The dissociation constant (K_D_) is also shown on the plots. (I) Overall and close-up views of the AfCycDesign prediction of the MDB_D8 design model highlighting key interactions with the MDM2.

While several hydrophobic interactions from the MCB_D2 helical segment are similar to those seen in natural MCL1 binders (e.g., BH3 peptide), (Figure S7), the N-to-C orientation of the helix is flipped in the case of MCB_D2. The loop region of MCB_D2 makes additional hydrophobic contacts and a cation-π interaction with MCL1 (Figure 2E, S7D) that we did not observe in previously reported natural MCL1 binders and their analogs. All three hits with an observable binding signal at 100 μM feature this cation-π interaction.

Encouraged by the experimental validation of the MCL1 binding macrocycles, we next sought to design binders to MDM2, an E3 ligase that interacts with tumor suppressor protein p53 and has multiple critical roles in tumor growth and survival^23^. We generated 10,000 macrocycle backbones spanning diverse lengths amenable to chemical synthesis (16-18 residues) and designed four amino acid sequences for each generated backbone using iterative rounds of ProteinMPNN and Rosetta Relax protocols (see Methods). Design models were filtered based on the confidence metrics and similarity of the AfCycDesign predictions to the designed complexes and the interface quality metrics calculated using Rosetta (Figure S4). AfCycDesign predicted 7,495 of the 40,000 design models to bind MDM2 with high confidence (normalized iPAE less than 0.3), (Figure S4). In contrast to our approach for MCL1, we chose not to do any additional filtering with RF2 as the results between AfCycDesign and RF2 were fairly consistent. After filtering on interface metrics (see Methods), we identified 17 designs with iPAE < 0.3, ddG < -50 kcal/mol, CMS > 300, and SAP score < 35. We selected 11 top-ranked designs by ddG for biochemical and biophysical characterization. The 11 selected designs had diverse sizes, shapes, and sequences (Figure S8 and Table S2); however, they were all predicted to bind the same site as the p53 transactivation domain (Figure S8). Three of the selected designs had poor yields during the cyclization step of the chemical synthesis, preventing further experimental characterization with them. We tested the remaining eight peptides for binding to the biotinylated MDM2 by SPR and identified three binders with observable binding signals at 100 μM (Figure S9). The best design, MDB_D8 (Figure 2G), demonstrated a binding affinity of 1.9 μM in the SPR single-cycle kinetics experiment (Figure 2H). The computational model for this design makes several key contacts at the interface that are similar to interactions observed in native MDM2-p53 complex structures (Figure 2I and S10)^23^. Despite different overall structures, all three hits from the SPR screen have a similar binding motif composed of phenylalanine, tryptophan, and either leucine or methionine from the helical segment of the macrocycle. Together, these data highlight the promising accuracy of the RFpeptides pipeline to design diverse macrocyclic binders for selected targets of interest.

### *De novo* design and characterization of macrocyclic binders to GABARAP

We next set out to design binders against a target with a binding site that is structurally different from MCL1 and MDM2, and formed by a mix of α-helices and β-strands (in contrast to all α-helical pockets of MCL1 and MDM2). We selected to target the Gamma-aminobutyric acid type A receptor-associated protein (GABARAP), a protein responsible for mediating autophagy through its role in autophagosome biogenesis and recruitment of cargo, resulting in lysosomal degradation of damaged or surplus proteins and organelles^24^. Peptide modulators against GABARAP could have therapeutic applications in the treatment of late-stage cancers^25^ or as chimeric peptides for autophagy-mediated targeted protein degradation (AutoTACs)^26^. Our target binding site for GABARAP, which is also the binding site for the native LC3-interacting region (LIR) or Atg8-interacting motif (AIM)^27^, is formed by a mix of β-strand and α-helix secondary structures (Figure 3 and S13). For designing macrocyclic binders against the human GABARAP, we used a similar pipeline as described above for MCL1 and MDM2 (see Methods), but we doubled the number of generated designs and defined six hotspot residues–Tyr49, Leu50, Lys48, Lys46, Phe60, and Leu63–to guide the macrocycle backbone generation to a specific site on the target (Figure 3A,D). We generated 20,000 macrocycle backbones and designed the amino acid sequences using ProteinMPNN and Rosetta Relax protocols. Out of the resulting 80,000 design models, we selected 335 macrocyclic designs based on the AfCycDesign (iPAE < 0.13) and Rosetta interface metrics (ddG < -30 kcal/mol, CMS > 300) (Figure S4). Instead of trying to synthesize and characterize all 335 cyclic peptides (which would have required significant time and experimental resources), we clustered the 335 designs into 80 different clusters based on their three-dimensional structures and selected representative designs from diverse clusters for further biochemical characterization. We selected 13 diverse macrocycles of 12-17 residues for synthesis and experimental validation (Table S3 and Figure S11). Unlike the design candidates described above for MCL1 and MDM2, several of the selected macrocycles for GABARAP show cyclic β-sheet structures with several edge-strand interactions with the target (Figure S11).

**Figure 3:**
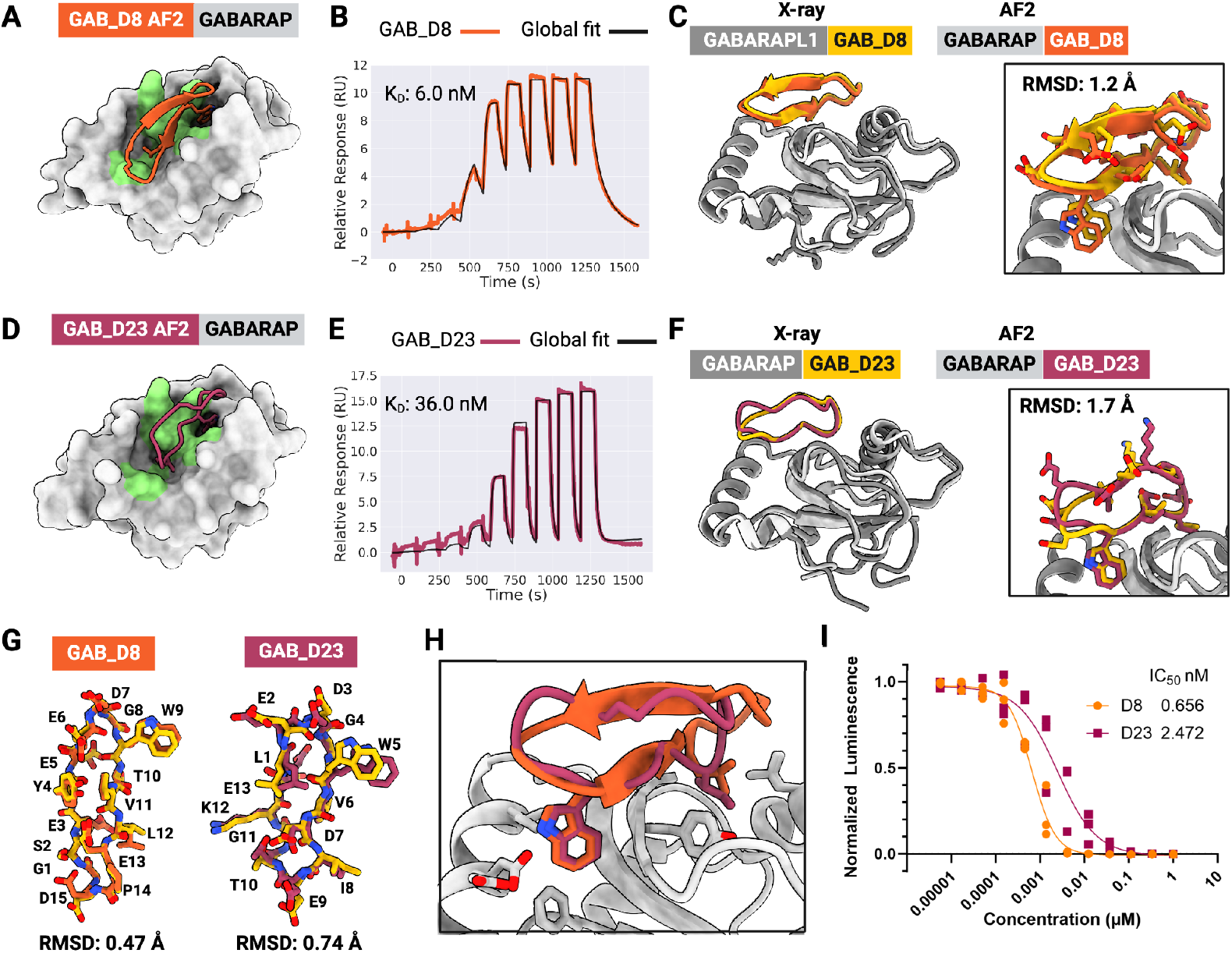
*De novo* design of high-affinity macrocycle binders to GABARAP. (A) AfCycDesign predicted model for design GAB_D8 bound to GABARAP shown as surface, with hotspot residues highlighted in green. (B) Affinity determination of GAB_D8 using SPR. SPR sensorgram from 9-point single-cycle kinetics experiments (5-fold dilution, highest concentration 20 μM). Experimental data are shown in orange and global fits are shown with black lines. The dissociation constant (K_D_) is also shown on the plot. (C) Superposition of chains E and F from the X-ray crystal structure of GAB_D8 bound to GABARAPL1 and the AfCycDesign model. (D) AfCycDesign predicted model for design GAB_D23 bound to GABARAP shown as surface, with hotspot residues highlighted in green. (E) Affinity determination of GAB_D23 using SPR. SPR sensorgram from 9-point single-cycle kinetics experiments (5-fold dilution, highest concentration 20 μM). Experimental data are shown in pink and global fits are shown with black lines. The dissociation constant (K_D_) is also shown on the plot. (F) Alignment of chains A and B from the X-ray crystal structure of GAB_D23 bound to GABARAP and the AfCycDesign model. (G) Alignments of GAB_D8 and GAB_D23 macrocycle models to X-ray crystal structures show close matches. (H) Comparison of GAB_D8 and GAB_D23 binding modes in the design models. (I) Competitive AlphaScreen response vs. concentration plot, IC_50_ from the average of three experiments. Donor and acceptor beads in the assay are bound to GABARAP and GABARAP-binding peptide K1, respectively.

We successfully synthesized six designs with high purity (> 90%) and tested them for binding to GABARAP using SPR (Figure S12). Two designs, GAB_D8 and GAB_D23, showed binding affinities of 6 nM and 36 nM, respectively (Figure 3B,E). To further characterize the binding of GAB_D8 and GAB_D23, we tested the ability of these designs to disrupt the interaction of GABARAP with linear peptide K1 (a previously described binder to this site^28^) in AlphaScreen assays. GAB_D8 and GAB_D23 demonstrated IC_50_ of 0.7 nM and 2.5 nM in the AlphaScreen assay, respectively (Figure 3I). To our knowledge, GAB_D8 is the most potent macrocyclic GABARAP binder to date.

In crystallization trials, we did not obtain crystals of sufficiently high quality for GAB_D8 bound to GABARAP. However, the X-ray crystal structure for GAB_D8 bound to GABARAPL1, a GABARAP homolog with 86% sequence identity, matches very closely with the design model, with a Cα RMSD of 1.2 Å over the macrocycle when aligned by the target protein to the closest of the four copies in the asymmetric unit (Figure 3C and Figure S13) and a Cα RMSD of 0.47 Å when aligned by macrocycle alone (Figure 3G). Notably, the X-ray structure of the GAB_D8–GABARAPL1 complex shows two different bound conformations of GAB_D8, one that closely matches the design model and a second one that partially deviates from the design model (Figure S14), with a register shift nucleated by Thr10 from the macrocycle forming main chain and side chain mediated hydrogen bonds with Lys48 on the target. GAB_D23 crystallized readily with GABARAP and also closely matches the design model with Cα RMSD of 1.7 Å when aligned by the target (Figure 3F) and Cα RMSD of 0.74 Å across the macrocycle alone (Figure 3G). The X-ray crystal structure confirms the key designed interactions, such as Trp5 and Ile8, with the main difference between the design model and the X-ray structure being the switch from a type I β-turn from Leu1 to Gly4 in the design model to a less regular conformation in the crystal structure, with a tendency for a type I’ β-turn from Glu2 to Trp5. While our original design models were predicted with single sequences as inputs to AF2, we retrospectively predicted the GAB_D8–GABARAPL1 and GAB_D23–GABARAP complex structures with multiple sequence alignment (MSA) inputs. These MSA-based predictions of the designs matched even more closely with the X-ray crystal structures, with Cα RMSDs of 0.5 Å and 0.9 Å for the GAB_D8–GABARAPL1 and GAB_D23–GABARAP complexes, respectively, when aligned by the target structure (Figure S15). Overall, these data demonstrate the ability of our *de novo* design pipeline to identify high-affinity binders against targets with diverse pocket shapes and surfaces without requiring library-scale screening.

### *De novo* design of macrocyclic binders against targets with unknown structures

Given the high accuracy and binding affinity of macrocycles designed against selected targets, we next set out to design macrocyclic binders against targets without any experimentally determined structure. We reasoned that the high accuracy of RFpeptides could mitigate the inherent risk of designing against a predicted target structure. We designed macrocycles against Rhombotarget A (RbtA), a recently identified cell surface protein from the ESKAPE pathogen, *Acinetobacter baumannii*. There are no experimentally determined structures available for this protein, and sequence-based searches against the Protein Data Bank do not return significant matches to other protein structures. We predicted the structure of the 617 amino acid full-length protein using AF2 and RF2; both methods predicted similar overall structures (Cα RMSD of 0.4 Å over 509 residues that exclude the signal peptide and transmembrane domain) with high confidence (pLDDT > 90), (Figure S16). AF2 and RF2 both predict two distinct extracellular domains: an N-terminal β-helix domain and a C-terminal Ig-like domain (Figure S16). While there were some differences in the predicted structures from AF2 and RF2, we decided to focus our binder design calculations on regions that were predicted nearly identically and with high confidence by AF2 and RF2. Based on our preliminary design runs without hotspots to guide the diffusion, we identified a patch in the N-terminal domain to pursue in our large-scale design calculations against this target and defined hotspots Leu144, Phe202, Phe204, Tyr206, Val208, Leu231, and Ala269 for peptide backbone generation(Figure 4A). In contrast to the concave pockets targeted for MDM2 and MCL1, this selected patch for RbtA is considerably flatter and difficult to target with conventional computational and experimental approaches (Figure S17). We generated 20,000 backbones for macrocycle binders and designed four amino acid sequences for each backbone using iterative rounds of ProteinMPNN and Rosetta Relax. Designs were filtered using AfCycDesign confidence metrics and Rosetta interface metrics, as described in earlier sections (Figure S4). Based on these *in silico* metrics, we selected 26 designs for biochemical and structural characterization with AfCycDesign iPAE < 0.28, ddG < -40 kcal/mol, RMSD between the design model and AfCycDesign prediction < 1.5 Å, and CMS more than 300. The selected designs cover diverse sizes (13-18 amino acids), sequences, shapes, and secondary structures (Figure S18 and Table S4). We expressed the Avi-tagged version of the RbtA N-terminal domain (residues 20-458) and used it for binding screens using SPR. Four out of 11 designs that were synthesized in sufficient quantity and purity showed a binding signal at 100 μM in our screens (Figure S19). Based on further binding experiments with SPR, we determined the dissociation constant of the best binder, RBB_D10, to be 9.4 nM (Figure 4B). The design model for RBB_D10 shows extensive contacts to the target with several side chain-mediated polar contacts and hydrophobic interactions (Figure 4F-H).

**Figure 4:**
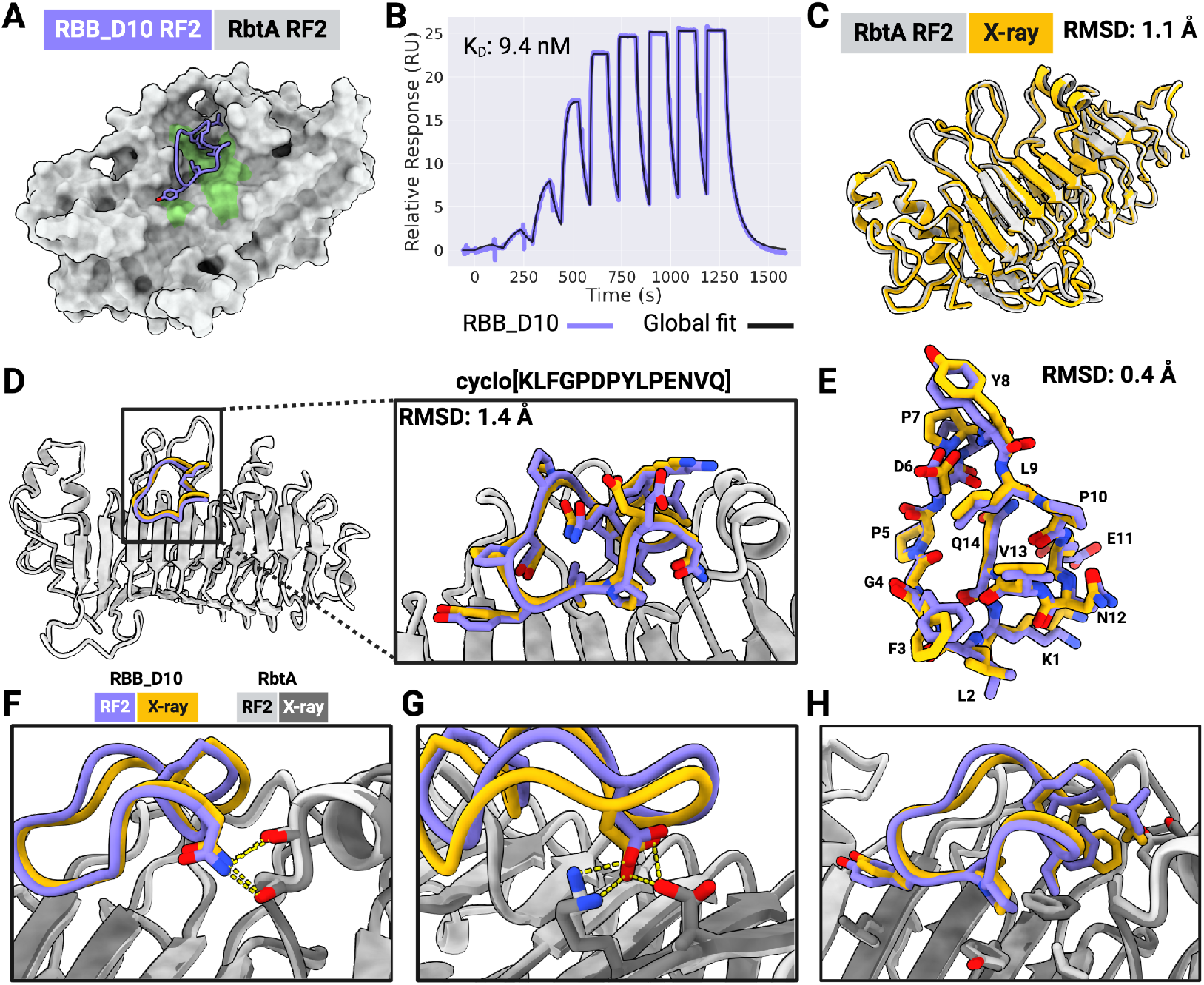
Accurate *de novo* design of a high-affinity cyclic peptide binder against the predicted structure of RbtA from *A. baumannii*. (A) AfCycDesign prediction of design RBB_D10 (violet cartoon) bound to the AF2 predicted β-helix domain of RbtA shown in gray surface. Hotspot residues from RbtA used during the backbone design step are shown in green. (B) SPR sensorgram from 9-point single-cycle kinetics experiment (5-fold dilution, highest concentration 20 μM). K_D_ determined from the SPR experiment also denoted on the plot. (C) Close agreement of the RF2-predicted structure of RbtA (gray) with the X-ray structure (gold) of the RbtA N-terminal domain determined here confirms the predicted structure of the target used for the macrocycle design calculations. (D) Alignment of the design model of RbtA-bound RBB_D10 (violet/gray) to the X-ray structure (in gold) shows a close match between the design model and the experimentally determined structure (Cɑ RMSD for macrocycle: 1.4 Å). Close-up view of the RbtA-bound RBB_D10 with side chains shown as sticks. (E) Overlay of RBB_D10 design model (after the AfCycDesign prediction step) aligned to the X-ray structure without RbtA demonstrates a nearly identical match for backbone coordinates and side chain rotamers (Cɑ RMSD: 0.4 Å). The design model and X-ray structure are shown in violet and gold, respectively. (F-H) Close-up views of the macrocycle-bound RbtA structure and the design model showing the accurate design of electrostatic and hydrophobic interactions at the binding interface.

To confirm the structures of RbtA and RBB_D10 and the binding mode between them, we determined the high-resolution X-ray crystal structure of apo and macrocycle-bound RbtA using X-ray crystallography at 2 Å and 2.6 Å resolution, respectively. The apo structure of the RbtA N-terminal domain–which is also the first experimentally determined structure from this class of bacterial proteins–matches our AF2/RF2 predictions for this target very closely, with an overall Cα RMSD of 1.2 Å and 1.1 Å between the X-ray structure of the RbtA N-terminal domain and the AF2/RF2 predicted structures, respectively (Figure 4C). The complex structure also confirms the structure and binding mode of our designed macrocycle, RBB_D10, with the X-ray structure matching the design model with an RMSD of 1.4 Å (Figure 4D). Notably, the conformation adopted by the macrocycle in the X-ray structure, including the side chain rotamers involved in interactions with the target, is almost identical to the design model with an RMSD of 0.4 Å (Figure 4E-H). Together, these data highlight the high accuracy and success rates provided by RFpeptides even while designing macrocycles against targets without deep pockets or targets with no known structures.

## DISCUSSION

Here, we describe RFpeptides, a generative DL pipeline for precise *de novo* design of macrocycle binders against a wide range of protein targets. The power of the approach is highlighted by the high affinities (Kd < 10 nM) of the designed macrocyclic binders to GABARAP and RbtA, and the nearly identical X-ray crystal structures and design models of the macrocycle-bound MCL1, GABARAP, and RbtA (Cα RMSDs of 0.7 Å, 1.2 Å, and 1.4 Å, respectively). The RFpeptides approach offers several advantages over traditional methods. First, the design approach should enable faster and more efficient discovery of macrocyclic binders. Despite testing less than 20 designed candidates per target (in contrast to trillions of peptides tested in traditional library-based approaches), we achieve high-affinity binders for two targets without requiring any further experimental optimization; to our knowledge, this is a considerably higher success rate than achieved with any previous method. Second, in contrast to the untargeted nature of the random library-based approaches, RFpeptides can be used for designing custom binders to specific patches and sites, as demonstrated for GABARAP and RbtA. Finally, the atomically accurate nature of the design models enables structure-guided optimization for properties beyond target binding (as well as further increases in affinity), bypassing the bottleneck of complex structure determination, which has hindered the optimization of leads from library screening. Combined with the design principles for membrane traversal, RFpeptides could enable the design of peptides simultaneously optimized for target binding and cell permeability or oral bioavailability.

RFpeptides also has considerable advantages over previous computational peptide design methods. Information on known ligands and/or binding partners is not required to initiate design: RFpeptides can design macrocycles completely *de novo* from just the structure or sequence (as in the case of RbtA) of the target, enabling design against molecular targets intractable with previous methods. RFpeptides is not limited to generating macrocycles with particular motifs or topologies: the diffusion process generates macrocycles with diverse shapes and sizes, and selects the topologies appropriate for the protein being targeted. Among the four targets tested here, binders for MCL1 and MDM2 have helical motifs, binders for GABARAP have a β-sheet topology, and binders for RbtA sample loop-like conformations that make extensive contacts with the flat surface of this target.

We anticipate RFpeptides will enable the rapid design of custom macrocyclic binders against a wide range of molecular targets, accelerating efforts to develop peptides for diverse functional applications. With the rapid advances in DL methods and frameworks, including the recent development of all-atom diffusion models, we aim to extend the approach to generative design of macrocycles with non-canonical amino acids, crosslinkers, and cyclization chemistries.

## ACKNOWLEDGEMENTS

We thank Lynda Stuart, Lance Stewart, Kandise VanWormer, Luki Goldschmidt, Madison Kennedy, Ian Haydon, Joshua Woodward, Xinting Li, Zoe Taylor, Heather Osterstock, Guangfeng Zhou, Gizem Gökçe, Jonathan Palmer, Kris Lindenauer, Katelyn Campbell, Matthias Gloegl, Robert Ragotte, and Martin Sadilek for their helpful feedback, guidance, and support. We also thank the IPD Peptide, Crystallography, and Biologics, Vaccines, and Process Development (BioVaxPD) core labs, and the UW Chemistry Mass Spectrometry Facility for providing instrumentation support and expertise. This work was supported by funds from the DARPA Harnessing Enzymatic Activity for Lifesaving Remedies (HEALR) program HR001120S0052 contract HR0011-21-2-0012 (GB, DB, JDM, ML), the Defense Threat Reduction Agency grant no. HDTRA1-19-1-0003 (DB, GB, SAR), HHMI Emerging Pathogens Initiative (JM, GB, VA), start-up funds from the University of Washington’s Department of Medicinal Chemistry and Institute for Protein Design (GB), the Audacious Project (GB, AKB, AK), C19 HHMI Initiative grant (AKB, AK), NIH R35 grant no. GM148407 (JAK, ANL), Marie Skłodowska-Curie Actions grant no. 101059124 (YFB), an NIH R01 grant no. R0AI160052 (AKB, AK), Deutsche Forschungsgemeinschaft (DFG, German Research Foundation)–Project-ID 267205415–SFB 1208 (JAW, AU, OHW, DW), and Bill and Melinda Gates Foundation grant GR047983 (DJ). All plots in this manuscript were generated using matplotlib or seaborn^29,30^. Peptide structures were rendered using ChimeraX 1.8^31^ or PyMOL 2.5.4. All figures in this manuscript were created using BioRender. This research used resources (FMX/AMX) of the National Synchrotron Light Source II, a U.S. Department of Energy (DOE) Office of Science User Facility operated for the DOE Office of Science by Brookhaven National Laboratory under Contract No. DE-SC0012704. The Center for Bio-Molecular Structure (CBMS) is primarily supported by the NIH-NIGMS through a Center Core P30 Grant (P30GM133893), and by the DOE Office of Biological and Environmental Research (KP1607011). This publication resulted from data collected using the beamtime obtained through NECAT BAG proposal # 311950. Finally, we would like to thank the staff of the ESRF and EMBL Grenoble for assistance and support in using beamline BM07 under proposal number MX-2587.

## CONTRIBUTIONS

S.A.R., D.J., V.A., F.D., and G.B. conceived the study. D.J. and F.D. implemented the cyclic relative positional offsets into RF2 and RFdiffusion. S.A.R., D.J., V.A., and G.B. developed the protocol for generating and filtering designs. A.L. and J.D.M. identified RbtA as a surface-exposed target in *A. baumannii*. S.A.R., V.A., M.B., P.M.L., M.S., and V.V. synthesized the designs. Y.F.B., Q.Z., M.B., S.R.G., A.L., J.A.W., A.U., and A.M. expressed and purified the target proteins. S.A.R., V.A., and A.N.L., biophysically characterized designed macrocyclic peptides. A.K.B., J.A.W., A.U., A.K., E.B., O.H.W., determined the X-ray crystal structures of the designed macrocycle peptides bound to their targets. S.O., O.H.W., D.W., J.A.K., J.D.M., D.B., F.D., and G.B. offered supervision throughout the project. S.A.R., D.J., V.A., D.B., and G.B. wrote the manuscript. All authors read and contributed to the manuscript. S.A.R., D.J., and V.A. agree that the order of their respective names may be changed for personal pursuits to best suit their interests.

## DATA AVAILABILITY

The design models and sequences are available in the Supplementary Information. Crystal structures of MCB_D2 bound to MCL1, GAB_D8 bound to GABARAPL1, GAB_D23 bound to GABARAP, RBB_D10 bound to RbtA, and apo RbtA have been deposited in the RCSB Protein Data Bank.

## CODE AVAILABILITY

The code and scripts for running the RFpeptides pipeline will be released upon publication. The scripts used for this work are also available as Supplementary Files.

