## supplementary_information for "Accurate *de novo* design of high-affinity protein binding macrocycles using deep learning": biorxiv_SI.pdf

**Table of Contents**

1. Figures S1 to S19 and Tables S1 to S4
2. Materials and Methods
    - 2.1. Macrocycle monomer design with RFpeptides
        - 2.1.1. Implementation of the cyclic relative position encodings for RoseTTAFold and RFdiffusion
        - 2.1.2. Monomeric macrocycle prediction with RFpeptides
        - 2.1.3. Monomeric macrocycle design with RFpeptides
        - 2.1.4. Visualization and structural clustering of monomeric designs
    - 2.2. Macrocycle Binder Design with RFpeptides
        - 2.2.1. Macrocycle Backbone Generation
        - 2.2.2. Sequence Design
    - 2.3. Target-macrocycle complex structure prediction using AfCycDesign
    - 2.4. Filtering designs based on physics-based metrics of interface quality
    - 2.5. Structure-based clustering for binder design campaigns
    - 2.6. Prediction of RbtA Structure
3. Experimental Methods
    - 3.1. Peptide Synthesis
    - 3.2. Protein Expression and Purification
        - 3.2.1. MDM2 and MCL1
        - 3.2.2. GABARAP for Surface Plasmon Resonance
        - 3.2.3. GABARAP and GABARAPL1 for Crystallography
        - 3.2.4. RbtA
    - 3.3. Determination of Binding Affinity by Surface Plasmon Resonance
    - 3.4. Determination of GABARAP Activity by AlphaScreen assay
    - 3.5. Crystallization of Protein-Cyclic Peptide complexes
        - 3.5.1. MCL1 with Cyclic Peptide
        - 3.5.2. GABARAPL1 with Cyclic Peptide
        - 3.5.3. GABARAP with Cyclic Peptide
        - 3.5.4. RbtA with Cyclic Peptide and apo RbtA
    - 3.6. Data Collection and Refinement Statistics
4. Peptide Analytical Characterization Data
5. References

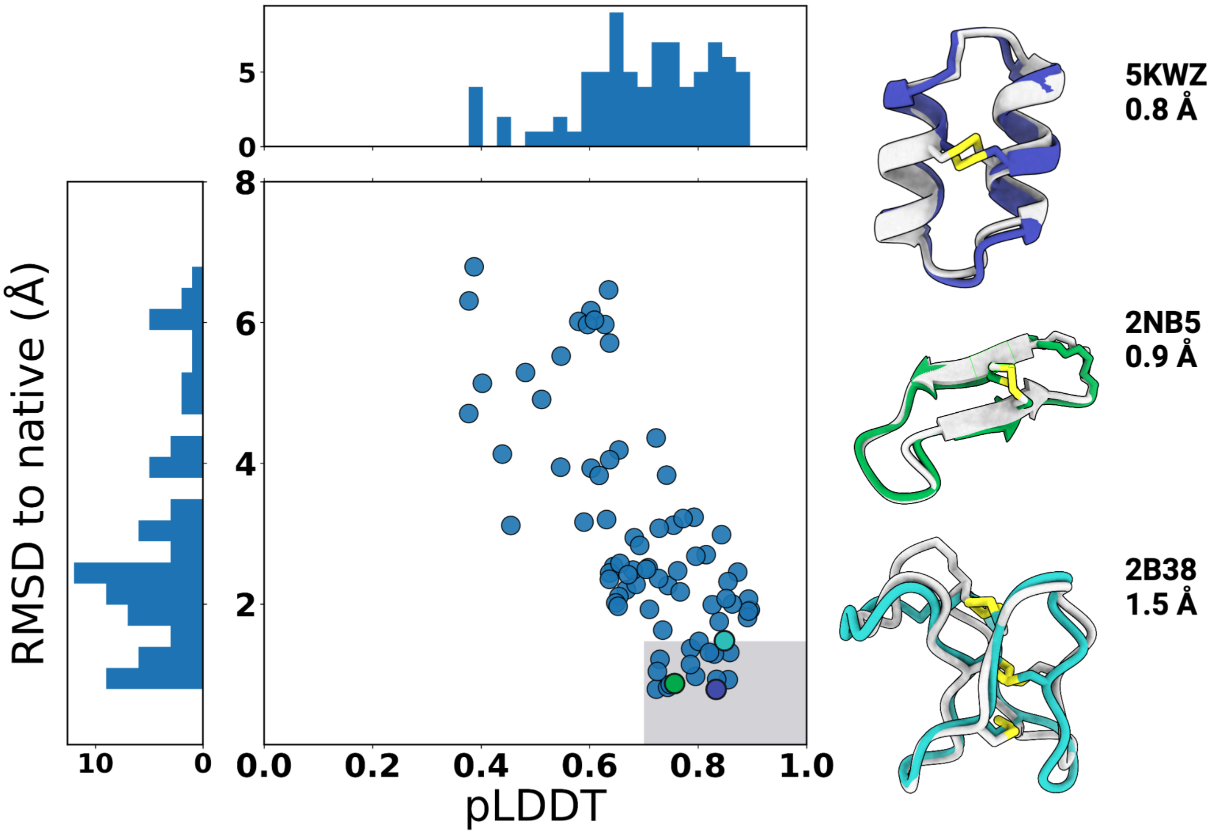

**Figure S1: Structure prediction of cyclic peptides using RoseTTAFold2 (RF2) modified with cyclic positional encoding.**

Structures for 80 cyclic peptides from the Protein Data Bank were predicted using RF2 modified with cyclic positional encoding. Each PDB entry is an NMR structure. The lowest pLDDT prediction was compared to all members of the NMR ensemble and the lowest backbone RMSD is plotted. Three selected examples from the benchmark set are shown with the NMR structure in gray and the predicted structure colored corresponding to its position in the plot. The boxed gray area encloses 17 high-confidence and accurate designs that were predicted within 1.5 Å of the NMR structure and with pLDDT of 0.7 or above.

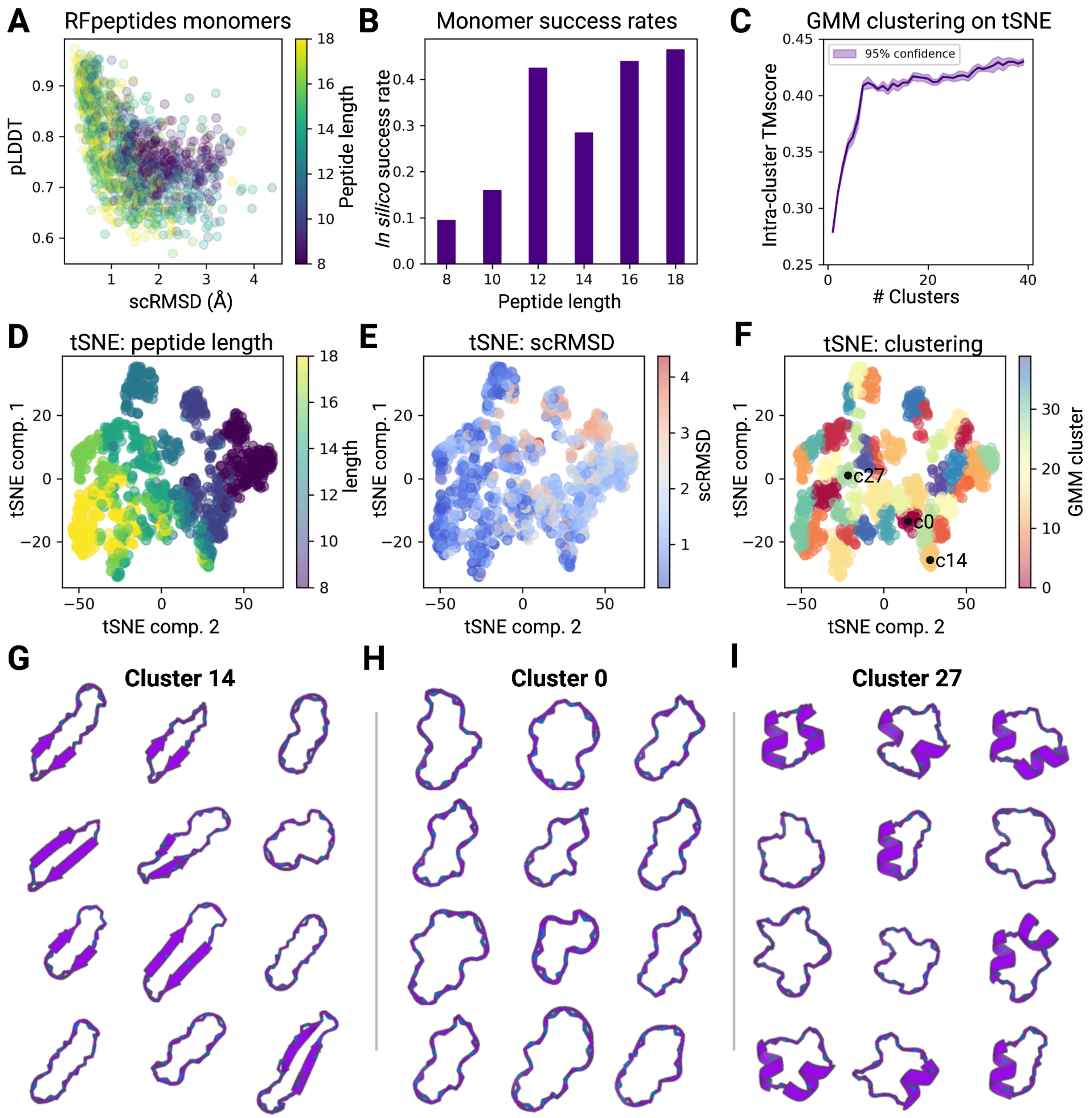

**Figure S2: *In silico* analysis of monomeric cyclic peptide generation by RFpeptides.** For each length in [8,10,12,14,16,18] amino acids, 200 cyclic peptide backbones were generated with RFpeptides. 8 amino acid sequences per backbone were then designed with LigandMPNN, and their structures predicted with AfCycDesign. (A) scRMSD ("self-consistency" RMSD) was calculated as the RMSD between the RFdiffusion output and AfCycDesign prediction (best of 8 LigandMNN sequences), and plotted against AfCycDesign pLDDT. (B) *In silico* success rates for peptides of varying length generated by the RFpeptides pipeline. A design is successful if its scRMSD < 2.0 Å and pLDDT > 0.8. This panel is reproduced with data from Figure 1 panel C for ease of reference and clarity. (C) Plot of intra-cluster mean TMscore versus number of clusters used when performing Gaussian mixture model (GMM) cluster assignment on 2D tSNE (see Methods 1.1.4 for more details). The TMscore metric reported here is the average of the query- and target-normalized TMscores output from TMalign. (D,E,F) tSNE components 1 and 2 plotted, and colored by (D) peptide length, (E) scRMSD and (F) cluster assignment after GMM clustering (reproduced from Fig 1B for ease of comparison). (G,H,I) Representative examples of generated backbones with < 2.0 Å scRMSD from (G) cluster 14, (H) cluster 0 and (I) cluster 27. We note two observations on the panels (G,H,I): First, it can be seen that internally to a given cluster, the structures are qualitatively more similar to each other than to structures in other clusters. E.g., Samples from cluster 14 are mostly extended, strand-pairing peptides with little-to-no alpha-helical structure, while samples from clusters (H) and (I) have little beta character. Second, although intra-cluster similarity is higher than inter-cluster similarity, there is still a wide range of significantly differing structures within each cluster. We attribute the “coarseness” of these clusters to the low overall number of Gaussians used (40 total) in the Gaussian mixture model. As can be seen in Fig. S2 C, using more Gaussians in the GMM results in higher intra-cluster TMscore, suggesting that if more clusters were used (e.g., hundreds), the clusters would become increasingly structurally uniform.

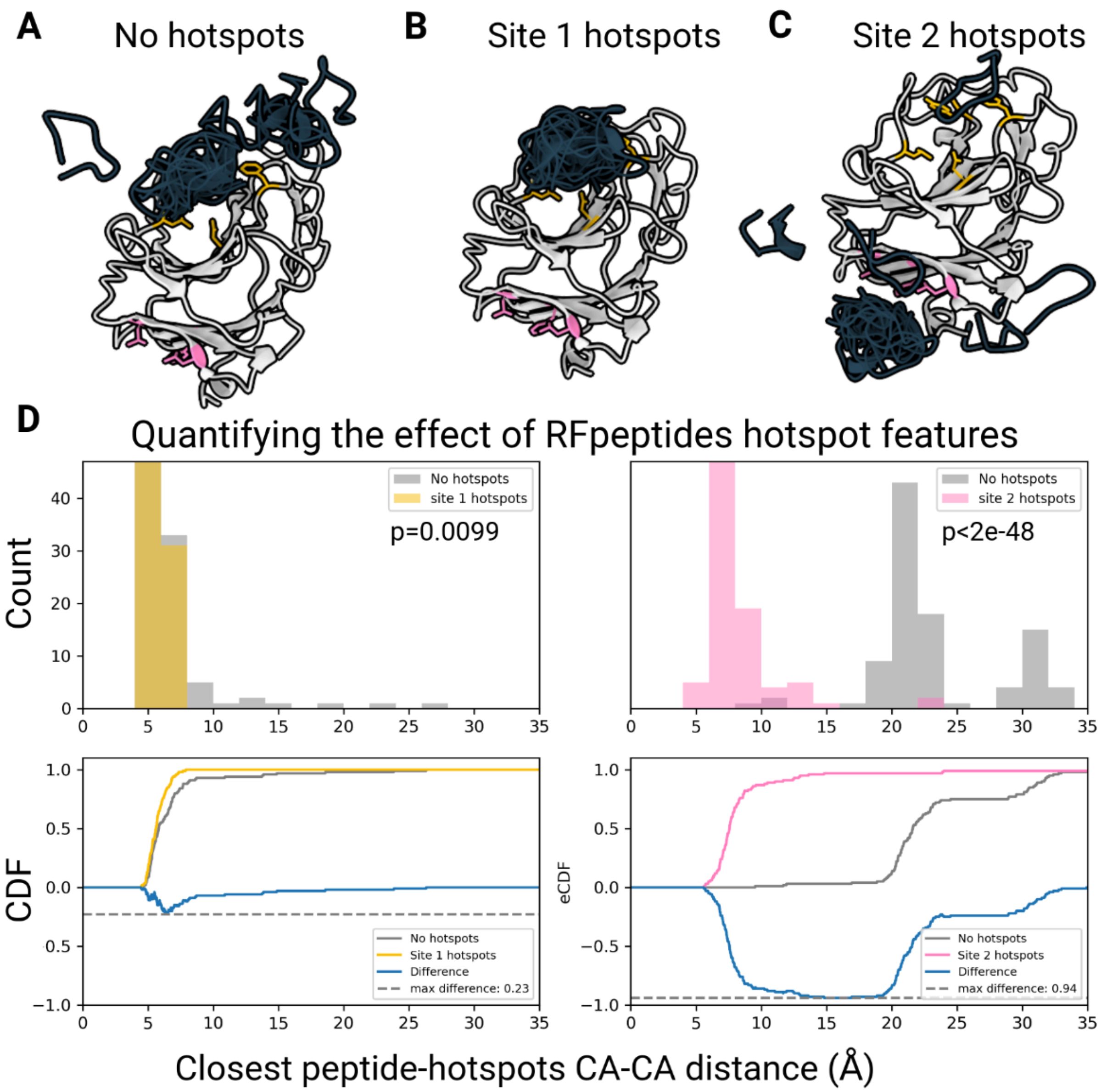

**Figure S3: Impact of specifying hotspot residues for cyclic peptide binder design**

RFpeptides macrocyclic peptide diffusion was sampled N=100 times using (A) no hotspot features, (B) hotspot features targeting residues Asp77, Tyr50, Phe171, Leu93 ("site 1" in yellow), (C) hotspot features targeting residues Phe131, Val138, Leu150 ("site 2" in pink). (D) For each of sites 1-3, with and without hotspot features provided, we computed the distribution of minimum Cα-Cα distances between any residue in the generated peptide and either of the two residues in the site (top). A two-sided Kolmogorov-Smirnov test was performed for each site, testing the hypothesis that the addition of hotspot features changes the distribution of minimum distances between the generated peptides and the epitope to be targeted. The null hypothesis ("hotspot features have no effect on the minimum distance distribution") is rejected with p-value 0.0099 for site 1, and $p < 2e^{-48}$ for site 2.

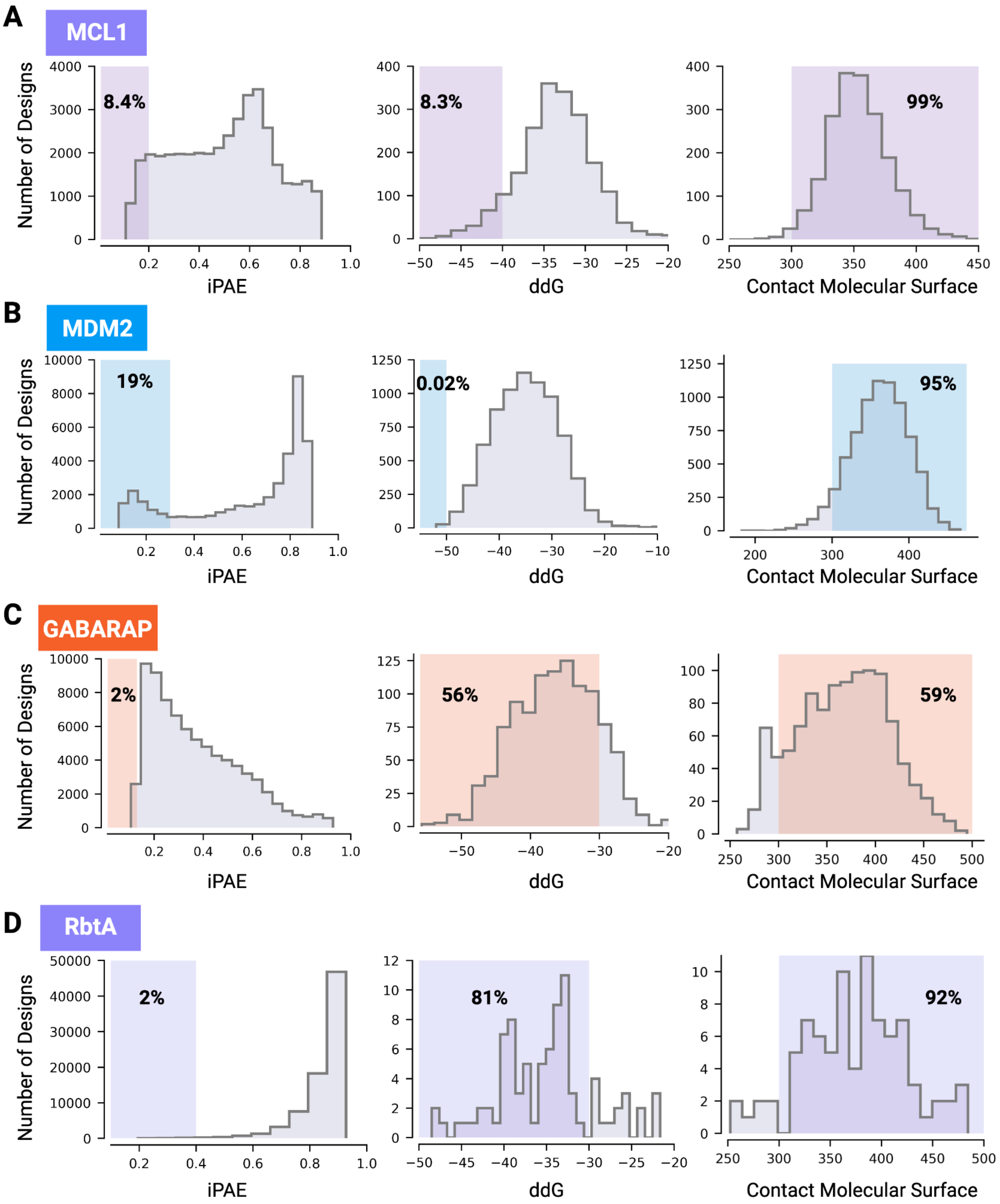

**Figure S4: Distribution of *in silico* scores used in downselection of design models**
Distribution of the interface PAE (iPAE) score from AfCycDesign predictions (left), Rosetta ddG (middle) and the contact molecular surface area (right) for the computationally generated macrocyclic binders against (A) MCL1, (B) MDM2, (C) GABARAP, and (D) RbtA. For all four targets, designs were first filtered by normalized iPAE from the AfCycDesign prediction of the

macrocycle-bound target complex, followed by further filtering using Rosetta ddG and the contact molecular surface area. The percentage of total designs passing the cutoff values (highlighted region) denoted in each plot.

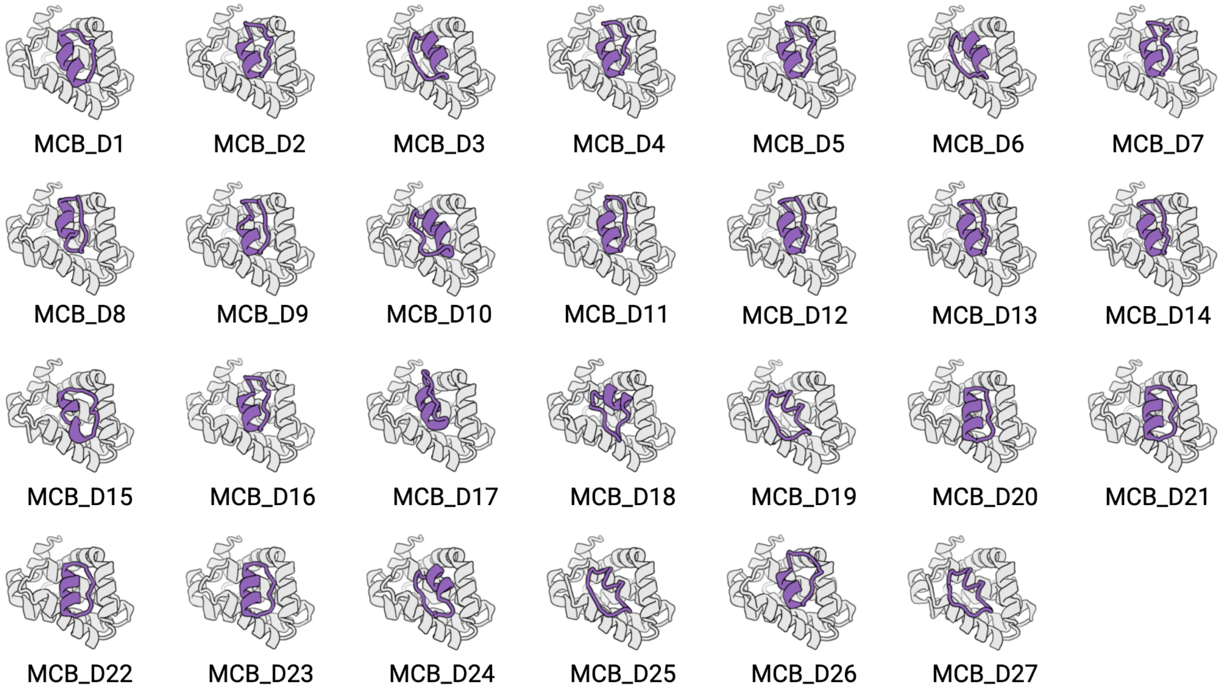

**Figure S5: Structural diversity of the selected MCL1 binding designs**

Computational models for 27 putative MCL1-binding macrocycles selected after filtering using AfCycDesign (normalized iPAE < 0.2), RoseTTAFold (iPAE < 6 and Cα RMSD to the design model less than 1 Å) and Rosetta interface metrics (ddG < -40, SAP < 35, CMS > 300). Two rank-ordered lists of candidates were generated, one using Rosetta metrics and the other using iPAE < 0.13. The two lists were merged, leaving 27 designs spanning those with the best Rosetta metrics, those with the best AfCycDesign metrics, and those with both.

**Table S1: Sequences and interacting residues for selected MCL1 designs**

| Design | Sequence | Length | Interacting Residues (< 4 Å) |
|---|---|---|---|
| MCB_D1 | GAPEILKKMADMVGYE | 16 | H224,T226,A227,F228,G230,M231,R233,K234,L235,V249,M250,H252,V253,F254,D256,G257,N260,G262,R263,T266,L267,F270 |
| MCB_D2 | PPEIAWLADAVGLKDA | 16 | V220,H224,A227,F228,G230,M231,K234,L235,V249,H252,V253,D256,R263,T266,L267,F270 |
| MCB_D3 | GLETDDPVVKPLADAV | 16 | H224,A227,F228,M231,K234,L235,V249,H252,V253,D256,G257,R263,T266,L267,F270 |
| MCB_D4 | APPEIKALADAVGLED | 16 | V220,H224,A227,F228,G230,M231,K234,L235,V249,H252,V253,R263,T266,L267,F270 |
| MCB_D5 | APESIKWLADAVGLKD | 16 | H224,A227,F228,G230,M231,K234,L235,V249,H252,V253,D256,R263,T266,L267,F270 |
| MCB_D6 | SLIKPLADAIGVETDD | 16 | H224,A227,F228,M231,K234,L235,V249,H252,V253,D256,R263,T266,L267,F270 |
| MCB_D7 | SSTVAFLQEAVGMPVT | 16 | V220,H224,A227,F228,G230,M231,K234,L235,V249,H252,V253,D256,R263,T266,L267,F270 |
| MCB_D8 | EAVGLESDSELIKKLA | 16 | V220,H224,A227,F228,G230,M231,K234,L235,V249,H252,V253,D256,G262,R263,V265,T266,L267,F270 |
| MCB_D9 | KPLAEAVGVETDIPTV | 16 | V220,H224,A227,F228,G230,M231,K234,L235,V249,H252,V253,R263,T266,L267,F270 |
| MCB_D10 | PAESELIAPLAEAVGI | 16 | H224,A227,F228,M231,K234,L235,V249,M250,H252,V253,S255,D256,R263,T266,L267,F270 |
| MCB_D11 | LEVPDTIVKPLAEAIG | 16 | V220,H224,A227,F228,G230,M231,K234,L235,V249,H252,V253,G262,R263,V265,T266,L267,F270 |
| MCB_D12 | SVDDPLIRKLAEAVGL | 16 | V220,H224,A227,F228,G230,M231,K234,L235,V249,H252,V253,D256,G262,R263,V265,T266,L267,F270 |
| MCB_D13 | SIDDPLLKKLAEAVGL | 16 | V220,H224,A227,F228,G230,M231,K234,L235,V249,H252,V253,D256,G262,R263,T266,L267,F270 |
| MCB_D14 | SIDDPLLRPLAEAVGL | 16 | H224,A227,F228,G230,M231,K234,L235,V249,H252,V253,G262,R263,T266,L267,F270 |
| MCB_D15 | ADALKRLTVPKNLEEI | 16 | V220,H224,T226,A227,F228,G230,M231,K234,L235,V249,H252,V253,S255,D256,R263,T266,L267,F270 |
| MCB_D16 | PPEVAFLADAVGLKDA | 16 | V220,H224,A227,F228,G230,M231,K234,L235,V249,H252,V253,R263,T266,L267,F270 |
| MCB_D17 | ERDDIIGPLVKAILGE | 16 | V220,H224,A227,F228,G230,M231,K234,L235,V249,H252,V253,G262,R263,V265,T266,L267,F270,F319 |
| MCB_D18 | KPQPDIDPAFAEIVGF | 16 | H224,A227,F228,G230,M231,K234,V249,H252,V253,S255,D256,R263,T266,L267,F270 |
| MCB_D19 | LAKIVGVETDDPVFAD | 16 | H224,A227,F228,M231,K234,L235,L246,V249,M250,H252,V25 |

| | | | |
|---|---|---|---|
| | | | 3,F254,D256,G257,R263,T266,L267,F270 |
| MCB_D20 | EDTLEGIARGLLTGKV | 16 | A227,F228,G230,M231,K234,L235,S245,R248,V249,H252,V253,R263,T266,L267,F270 |
| MCB_D21 | YDTEEGIAEGLLTGKV | 16 | H224,T226,A227,F228,G230,M231,K234,L235,S245,L246,R248,V249,H252,V253,R263,T266,L267,F270 |
| MCB_D22 | ADTEEGIAKGLLTGKV | 16 | H224,T226,A227,F228,G230,M231,K234,L235,R248,V249,H252,V253,R263,T266,L267,F270 |
| MCB_D23 | ADTIEGIAKGLLTGEV | 16 | H224,T226,A227,F228,G230,M231,K234,L235,S245,R248,V249,H252,V253,R263,T266,L267,F270 |
| MCB_D24 | PEIAEDSRQIFGVDLD | 16 | H224,A227,F228,G230,M231,K234,L235,V249,M250,H252,V253,S255,D256,R263,T266,L267,F270 |
| MCB_D25 | LAKIVGVETSDEKIED | 16 | H224,A227,F228,M231,K234,L235,L246,V249,M250,H252,V253,F254,D256,G257,R263,T266,L267,F270 |
| MCB_D26 | GSPEIRWLMDAFGVDE | 16 | N223,H224,T226,A227,F228,G230,M231,R233,K234,L235,L246,V249,H252,V253,D256,R263,T266,L267,F270 |
| MCB_D27 | LAKIVGVTDAPEEIAD | 16 | H224,A227,F228,M231,K234,L235,L246,V249,M250,H252,V253,F254,D256,G257,R263,T266,L267,F270 |

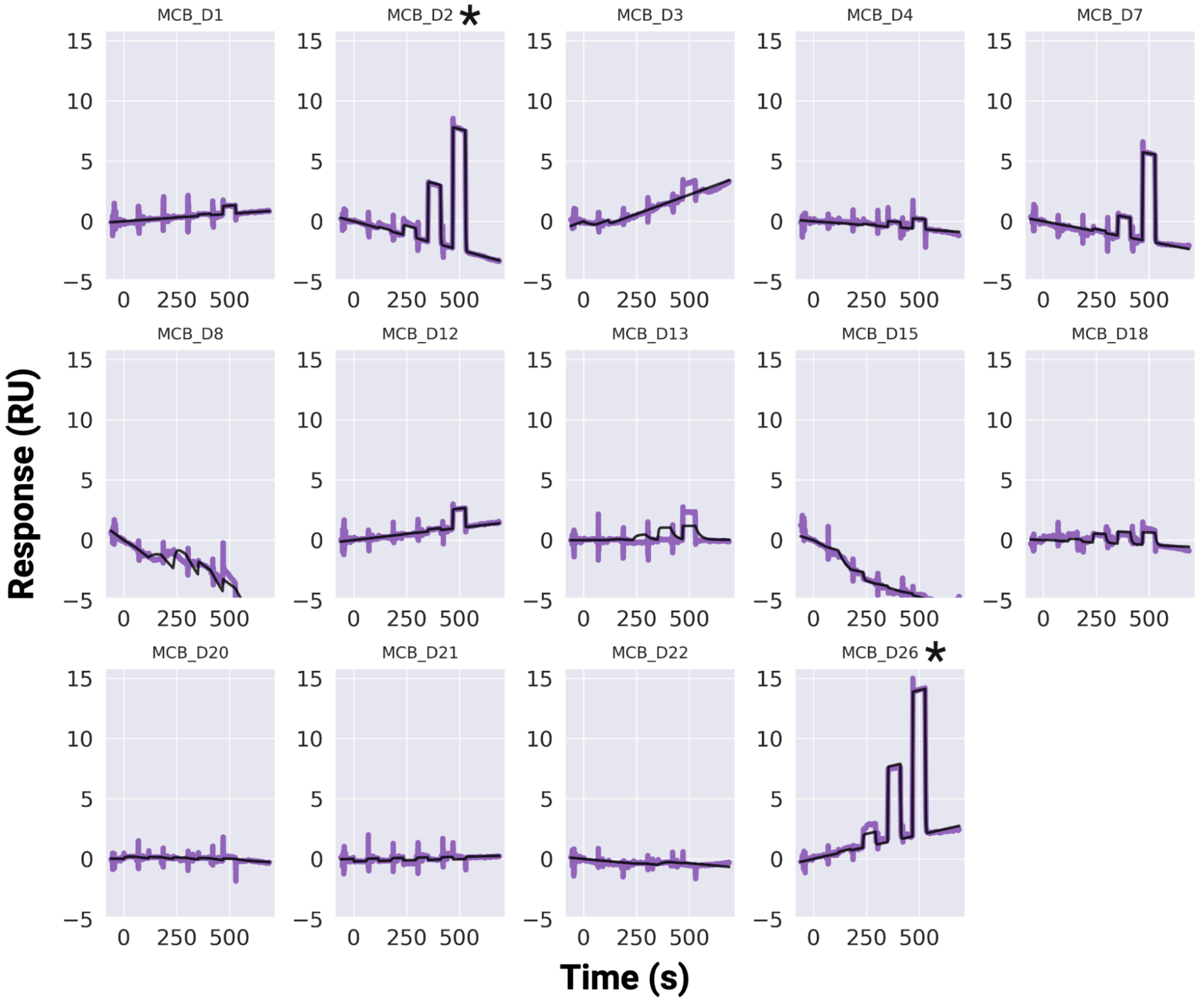

**Figure S6: SPR-based binding screen of the selected MCB designs against MCL1**

Single cycle kinetics SPR binding screen for selected MCB designs, applying a 5-point 10-fold dilution with 100 μM as the highest concentration. Three designs (MCB_D2, MCB_D7, and MCB_D26) demonstrate detectable binding signals at or below 100 μM. MCB_D2 and MCB_D26 denoted with * were chosen for further characterization.

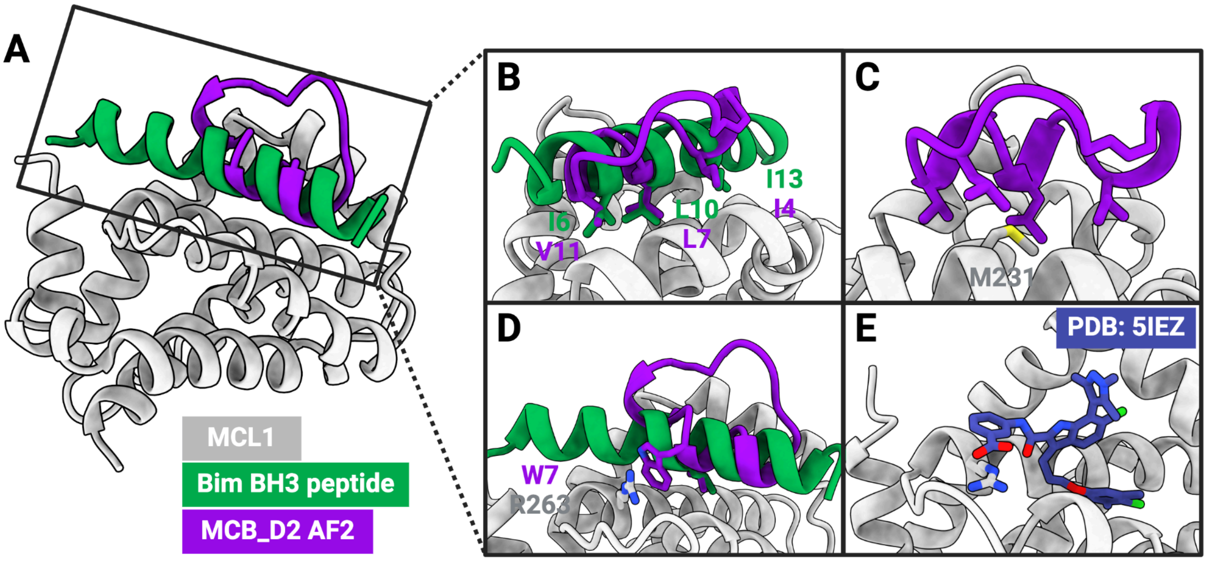

**Figure S7: Comparison between the MCL1–MCB_D2 interactions and interactions made by previously described MCL1 binders**

(A) Alignment of Bim-BH3 (PDB: 2PQK[1]) to the design model for MCB_D2 demonstrates that the direction of the interacting helix is opposite across the two cases. (B) Some of the side chain placements at the interface of the Bim-BH3–MCL1 complex are mimicked in the design model of MCB_D2. (C) The helix and loop regions of MCB_D2 make hydrophobic contacts with MCL1 (D) Cation-π interaction between W7 of MCB_D2 and R263 of MCL1 not present in native Bim-BH3. (E) A small-molecule ligand bound to MCL1 (PDB: 5IEZ[2]) shows a similar cation-π interaction with R263.

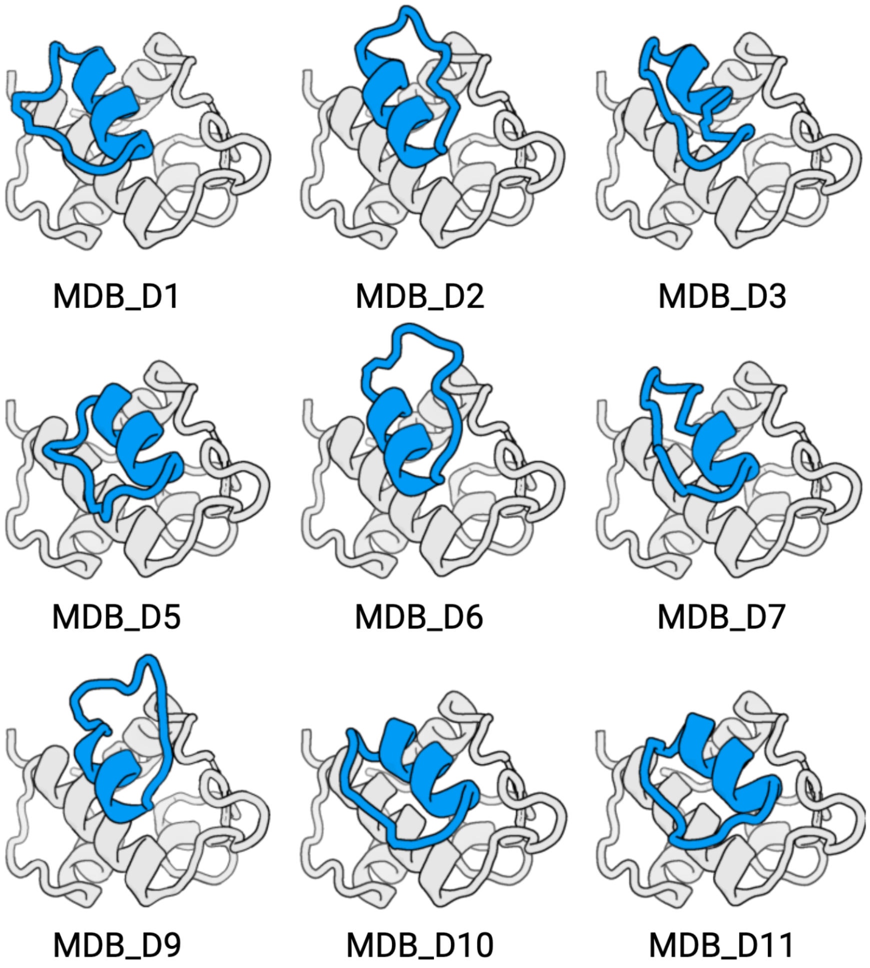

**Figure S8: Structural diversity of the selected MDM2 binding designs**

Computational models for 11 putative MDM2-binding macrocycles selected after filtering using AfCycDesign (normalized iPAE < 0.3) and Rosetta interface metrics (ddG < -50, SAP < 35, CMS > 300). The designs that passed the computational filters were sorted by ddG, and the top 11 designs were chosen for synthesis.

**Table S2: Sequences and interacting residues for selected MDM2 designs**

| Design | Sequence | Length | Interacting Residues (< 4 Å) |
|---|---|---|---|
| MDB_D1 | KKYNWMVDELVSMVGKPE | 18 | M50,K51,L54,F55,L57,G58,Q59,I61,M62,Y67,Q72,H73,V75,F91,V93,K94,H96,I99,Y100,I103 |
| MDB_D2 | EESEDEDFRVFERVLGI | 17 | M50,K51,L54,F55,L57,G58,Q59,I61,M62,Y67,Q72,H73,V75,F91,V93,K94,H96,I99,Y100,I103 |
| MDB_D3 | RELMEMVGEKYDQNFIL | 17 | M50,K51,L54,F55,L57,G58,Q59,I61,M62,Y67,Q72,H73,V75,F91,V93,K94,H96,I99,Y100,I103 |
| MDB_D4 | IESDDDFKVLTDVMGLD | 17 | M50,K51,L54,F55,L57,G58,Q59,I61,M62,Y67,Q72,H73,V75,F91,V93,K94,H96,I99,Y100,I103 |
| MDB_D5 | DGSEFSKIWKSLGGDQDL | 18 | K51,L54,F55,L57,G58,Q59,I61,M62,Y67,Q72,H73,V75,F86,F91,V93,K94,H96,I99,Y100,I103 |
| MDB_D6 | EDYQVLHDVLGLPLEFDD | 18 | M50,K51,L54,F55,L57,G58,Q59,I61,M62,Y67,Q72,H73,V75,F91,V93,K94,H96,I99,Y100,I103 |
| MDB_D7 | MAMVGEELEGADFVLREL | 18 | M50,K51,L54,F55,L57,G58,Q59,I61,M62,Y67,Q72,H73,V75,F91,V93,K94,H96,I99,Y100,I103 |
| MDB_D8 | SRKAKNKFEELWNELIDP | 18 | K51,L54,F55,L57,G58,I61,M62,Y67,Q72,H73,V75,F91,V93,K94,H96,I99,Y100 |
| MDB_D9 | FKVLDEVLGVDVEDADEL | 18 | M50,K51,L54,F55,L57,G58,Q59,I61,M62,Y67,Q72,H73,V75,F91,V93,K94,H96,I99,Y100,I103 |
| MDB_D10 | PGTPFAKEWSKMAGGAPL | 18 | K51,L54,F55,L57,G58,Q59,I61,M62,Y67,Q72,H73,V75,F86,F91,V93,K94,H96,I99,Y100,I103 |
| MDB_D11 | GLDLETDSEFAKEWQKMV | 18 | K51,L54,F55,L57,G58,Q59,I61,M62,Y67,Q72,H73,V75,F91,V93,K94,H96,I99,Y100,I103 |

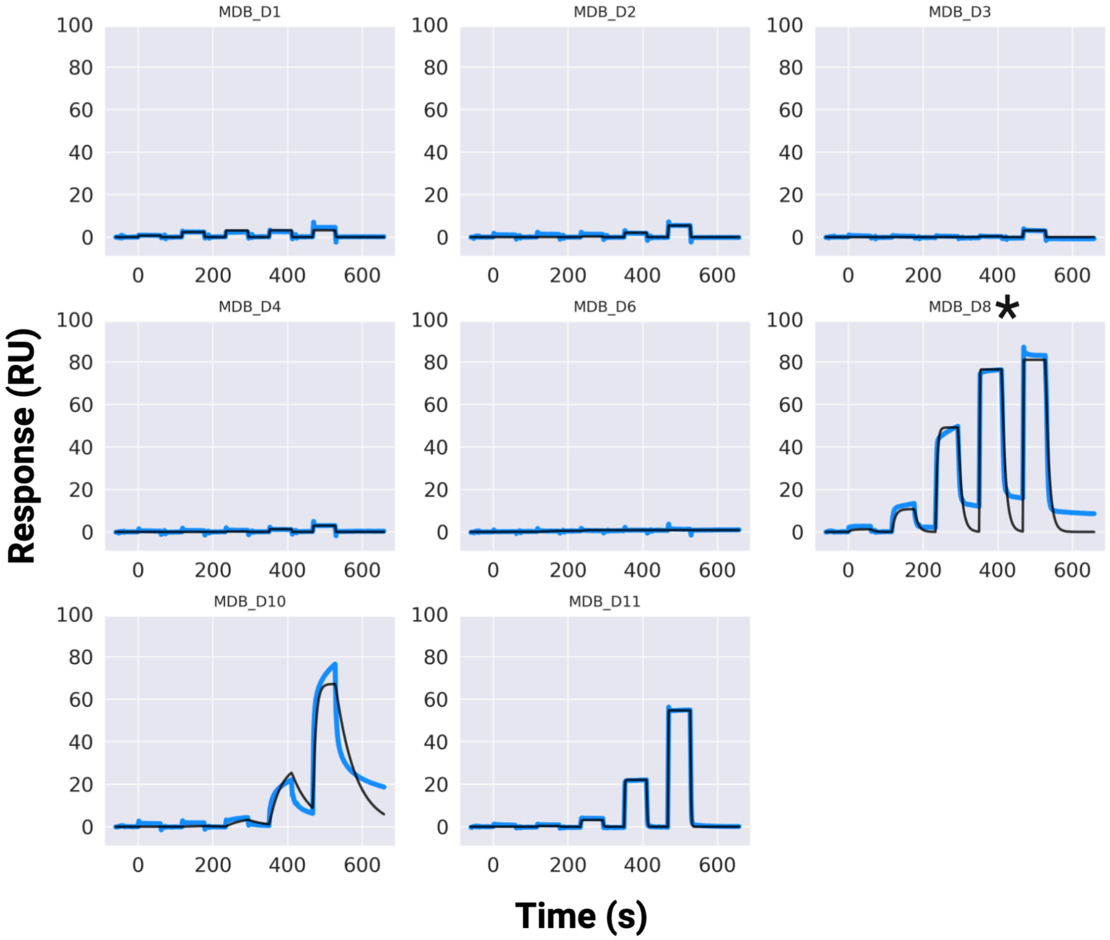

**Figure S9: SPR-based binding screen of the selected MDB designs against MDM2**

Sensorgrams from an SPR single cycle kinetics screen for the selected MDB designs, applying a 5-point 10-fold dilution with 100 µM as the highest concentration. Three designs (MDB_D8, MDB_D10, and MDB_11) show detectable binding at or below 100 µM. MDB_D8 denoted with * chosen for further characterization.

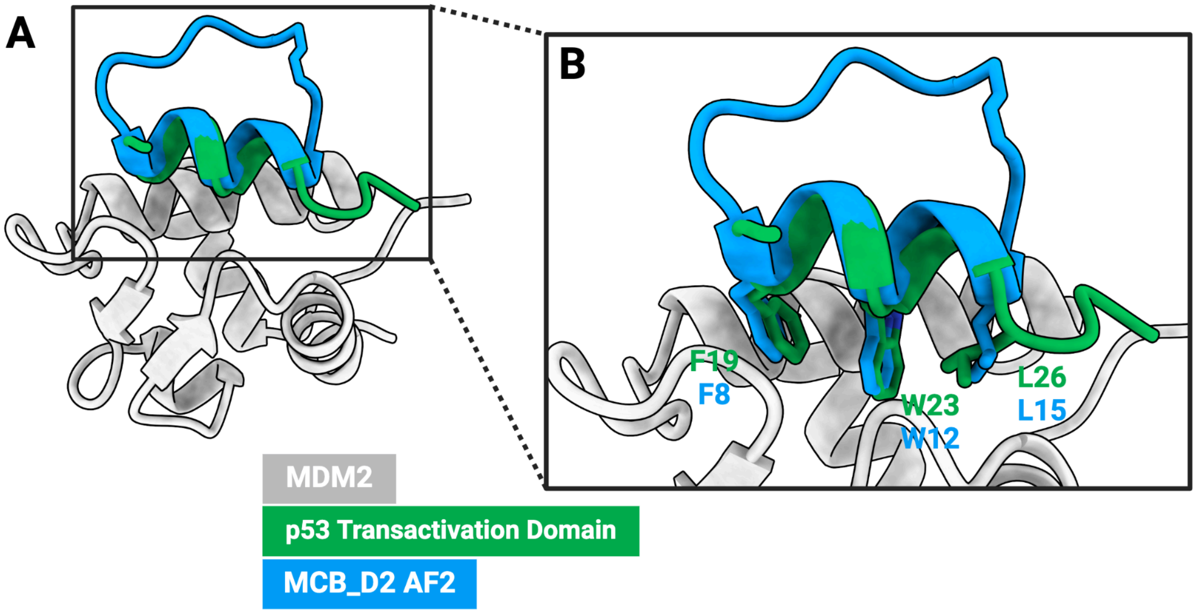

**Figure S10: MDB_D8 mimics p53 interactions**

(A) Alignment of the MDB_D8 design model to the crystal structure of p53 bound to MDM2 (PDB: 1YCR[3]). (B) Positions and identities of interfacial side chains in MDB_D8 match those in the native ligand.

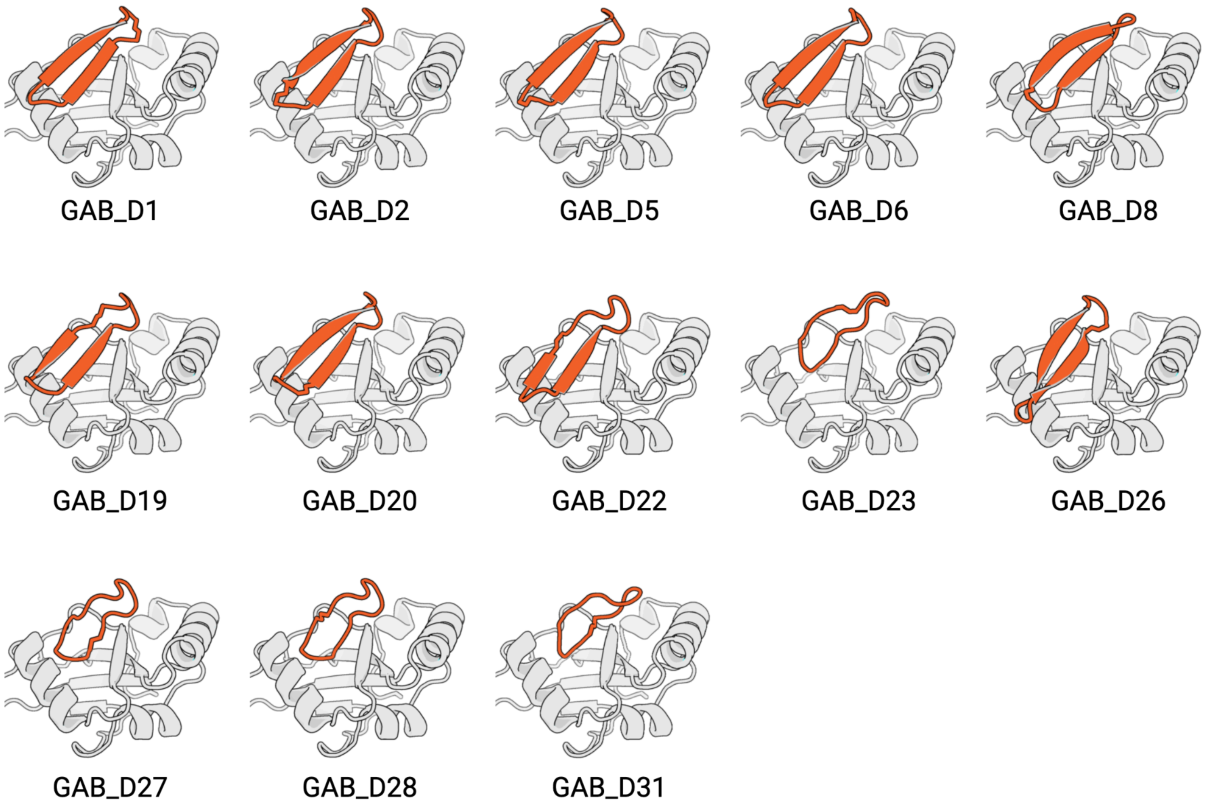

**Figure S11: Structural diversity of the selected GABARAP binding designs**
Computational models for 13 putative GABARAP-binding macrocycles selected after filtering using AfCycDesign (normalized iPAE < 0.13) and Rosetta interface metrics (ddG < -30, SAP < 35, CMS > 300). Post filtering, designs were clustered and 13 different designs from diverse clusters were selected for synthesis and experimental screening.

**Table S3: Sequences and interacting residues for selected GABARAP designs**

| Design | Sequence | Length | Interacting Residues (< 4 Å) |
|---|---|---|---|
| GAB_D1 | ETGEVENIDGVEIIYP | 16 | I23,R26,K48,K50,Y51,L52,V53,P54,F64,L65,K68 |
| GAB_D2 | DGVTIIDIDTGEEEVL | 16 | I23,R26,K48,K50,Y51,L52,P54,L57,F64,L65,K68,R69 |
| GAB_D5 | GIEIIELETGEVVVID | 16 | I23,R26,K48,K50,Y51,L52,P54,L57,L65,K68,R69 |
| GAB_D6 | GEVEVIDGVEIIDLET | 16 | I23,R26,K48,K50,Y51,L52,P54,L57,L65,K68,R69 |
| GAB_D8 | GSEYEEDGWTVLEPD | 15 | I23,K26,Y27,R30,K48,K50,Y51,L52,P54,Q61,F62,L65,R69,F102 |
| GAB_D19 | KEKWDENIEIINIETGE | 17 | E19,I23,R30,K48,K50,Y51,L52,P54,L57,L65,K68 |
| GAB_D20 | IEIIDLDTGEVEVWSPD | 17 | I23,R30,K48,K50,Y51,L52,P54,L57,F62,L65,R69 |
| GAB_D22 | MELDGVKVLDIDTEEVV | 17 | I23,K26,Y27,R30,K48,K50,Y51,L52,V53,P54,L57,F64,L65,R69 |
| GAB_D23 | LEDGWVDIETGKE | 13 | E19,I23,K26,Y27,R30,P32,V33,K48,K50,Y51,L52,V53,P54,F62,F106 |
| GAB_D26 | LDTGEVYKAPNGQEVIE | 17 | I23,R30,K48,K50,Y51,L52,V53,P54,D56,L57,Q61,F64,L65,R69 |
| GAB_D27 | IDIDTEEEVMPGV | 13 | I23,K26,R30,K48,K50,Y51,L52,V53,P54,L65 |
| GAB_D28 | VMPGIIDIDTEEE | 13 | I23,K26,R30,K48,K50,Y51,L52,P54,F62,L65 |
| GAB_D31 | DIVTDEELDGYV | 12 | E19,K26,Y27,R30,K48,K50,Y51,L52,V53,P54,F62,R69,F106 |

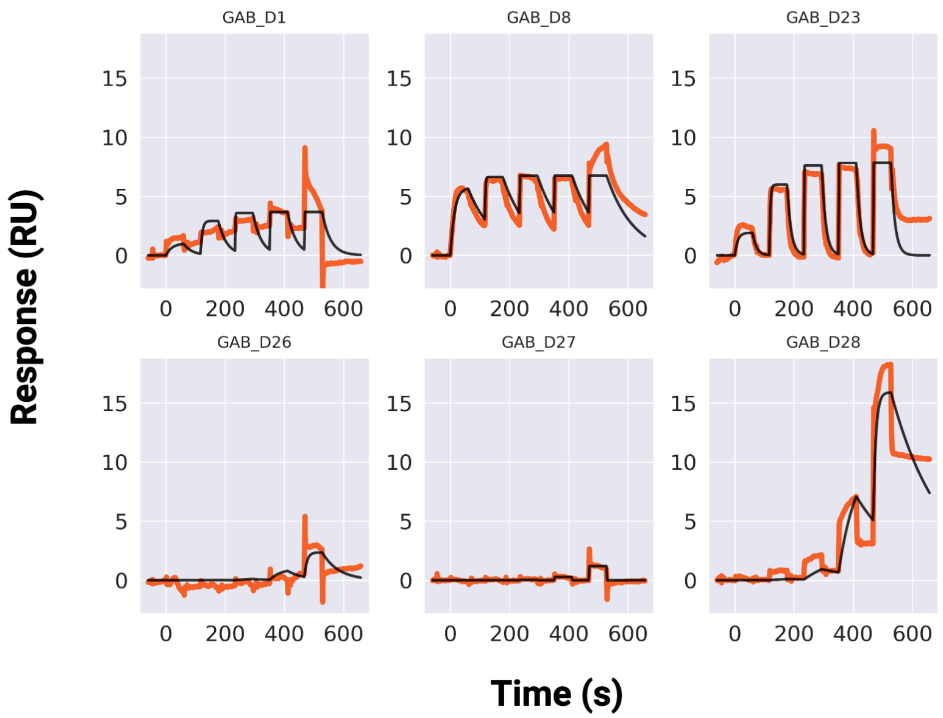

**Figure S12: SPR-based screening of the selected GAB designs against GABARAP**
Sensorgrams from an SPR single cycle kinetics screen for the selected GAB designs, applying a 5-point 10-fold dilution with 100 µM as the highest concentration. Four designs (GAB_D1, GAB_D8, GAB_D23, and GAB_D28) show detectable binding at or below 100 µM. GAB_D8 and GAB_D23 were chosen for further characterization.

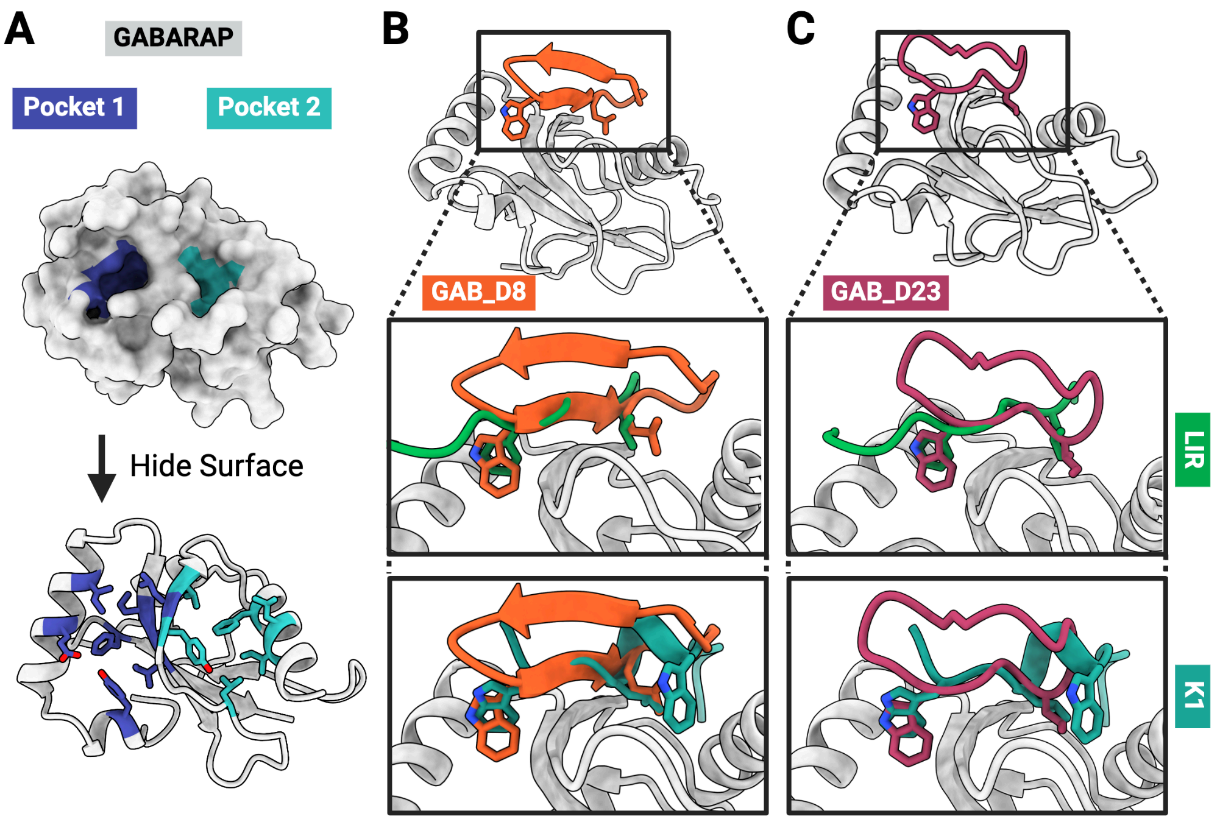

**Figure S13: GAB_D8 and GAB_D23 span both known pockets on GABARAP**
(A) GABARAP has two hydrophobic pockets that are targeted by typical LIR peptides (exemplified by PDB entry 5LXI[1]) as well as the artificial ligand K1 (PDB: 3D32[4]). Both GAB_D8 (B) and GAB_D23 (C) interact with pocket 1 via an aromatic residue (tryptophan) and with pocket 2 via an aliphatic residue (leucine and isoleucine, respectively), thus mimicking the canonical binding mode found in native ligands. In contrast, the K1 peptide engages in the reverse orientation and inserts a second tryptophan side chain, in addition to a leucine, into pocket 2.

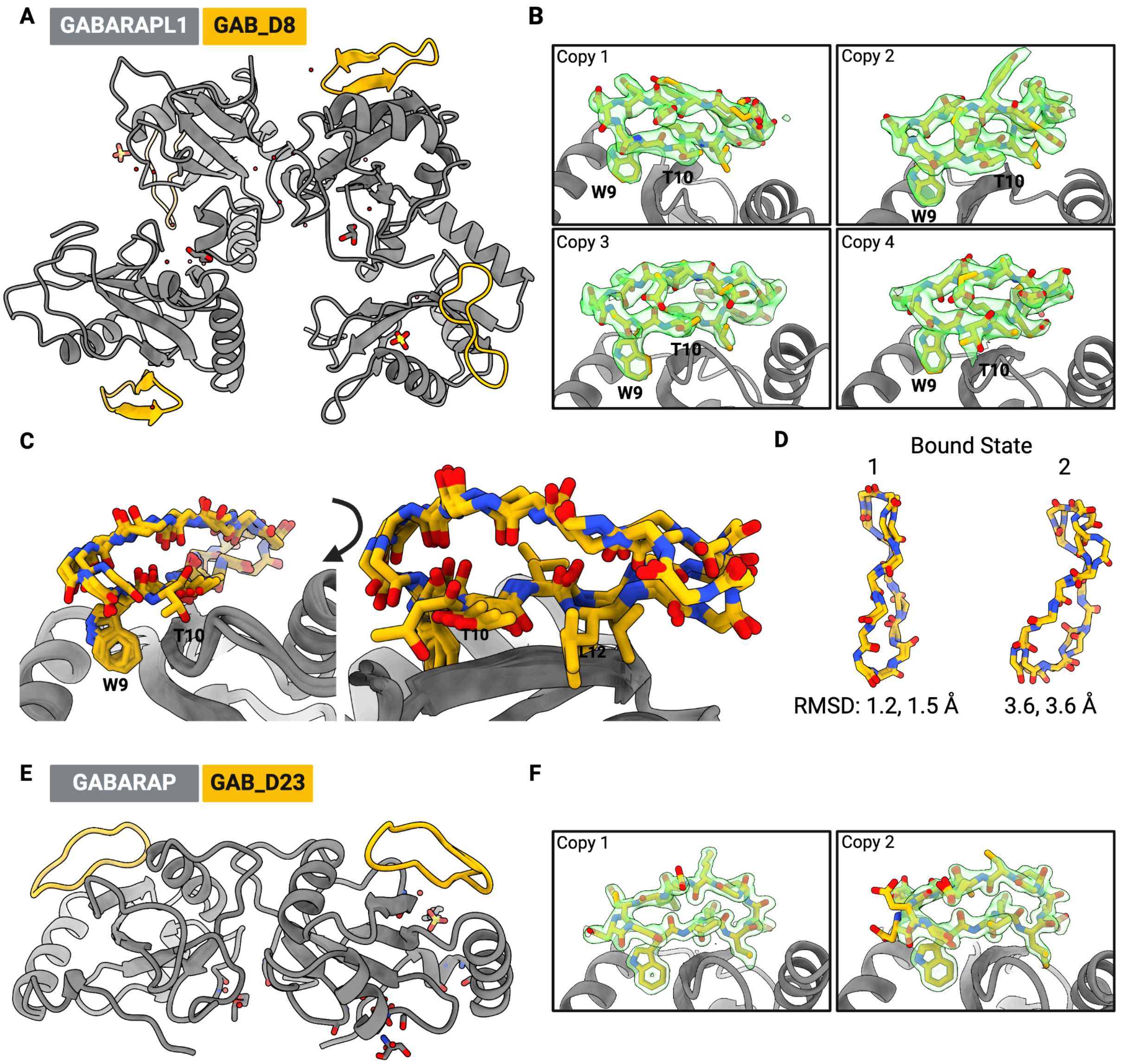

**Figure S14: GAB_D8 and GAB_D23 crystal structures display secondary conformations.** (A) The asymmetric unit contains four copies of GAB_D8 bound to GABARAPL1. (B) Electron densities (2Fo-Fc map at 1 σ contour level, green) are shown for each copy of the peptide. Note that copy 4 deviates from the other copies, presumably as a result of lattice restraints. Specifically, Thr10 and Val11 bulge out, establishing a new side chain-to-backbone hydrogen bond to GABARAPL1 and causing a register shift in the strand pairing. (C) Alignment of all four models. The difference at Thr10 for copy 4 can be seen, as well as the register shift leading to Leu12 moving to the other side of the strand. (D) The four copies can be grouped into two bound states. Both state 1 copies closely match the design with Cα RMSDs < 2 Å, but state 2 differs from the designed fold and contains copy 4 with its alternate binding mode. (E) The asymmetric unit contains two copies of GAB_D23 bound to GABARAP. (F) Electron densities (2Fo-Fc map at 1 σ contour level, green) are shown for each copy of the peptide. Note that copy 2 contains a second conformer to more completely account for the features of the electron density.

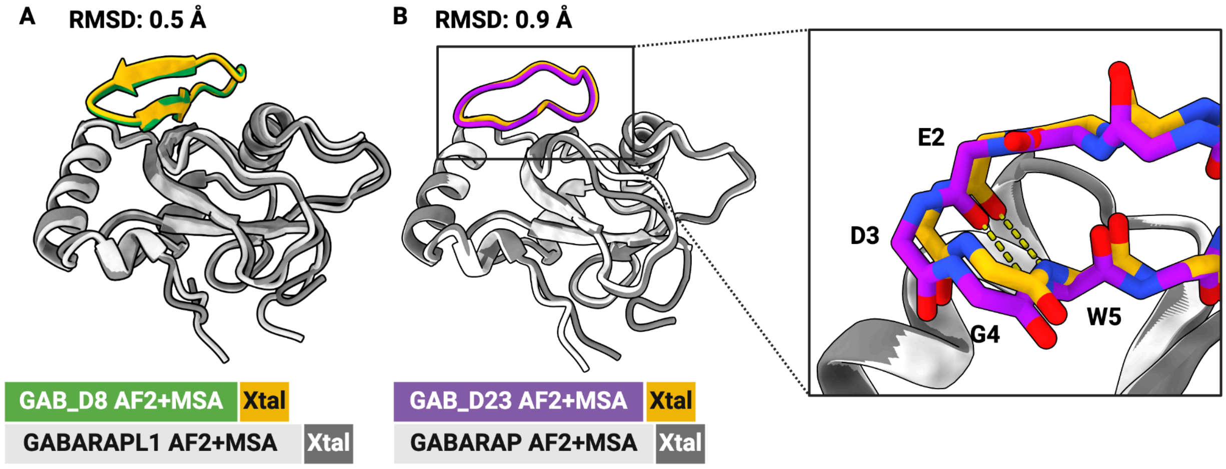

**Figure S15: Refolding of GAB_D8 with GABARAPL1 and GAB_D23 with GABARAP using multiple sequence alignment (MSA) improves accuracy of the design models**

(A) The GAB_D8-GABARAPL1 complex predicted with MSA aligned to the crystal structure. Cα RMSD calculated for the peptide aligned by the target chain. (B) The GAB_D23-GABARAP complex predicted with MSA aligned to the prevailing conformer found in the crystal structure. Cα RMSD calculated for the peptide aligned by the target chain. The close-up view shows capture of the type I' β-turn from the crystal structure in the MSA prediction.

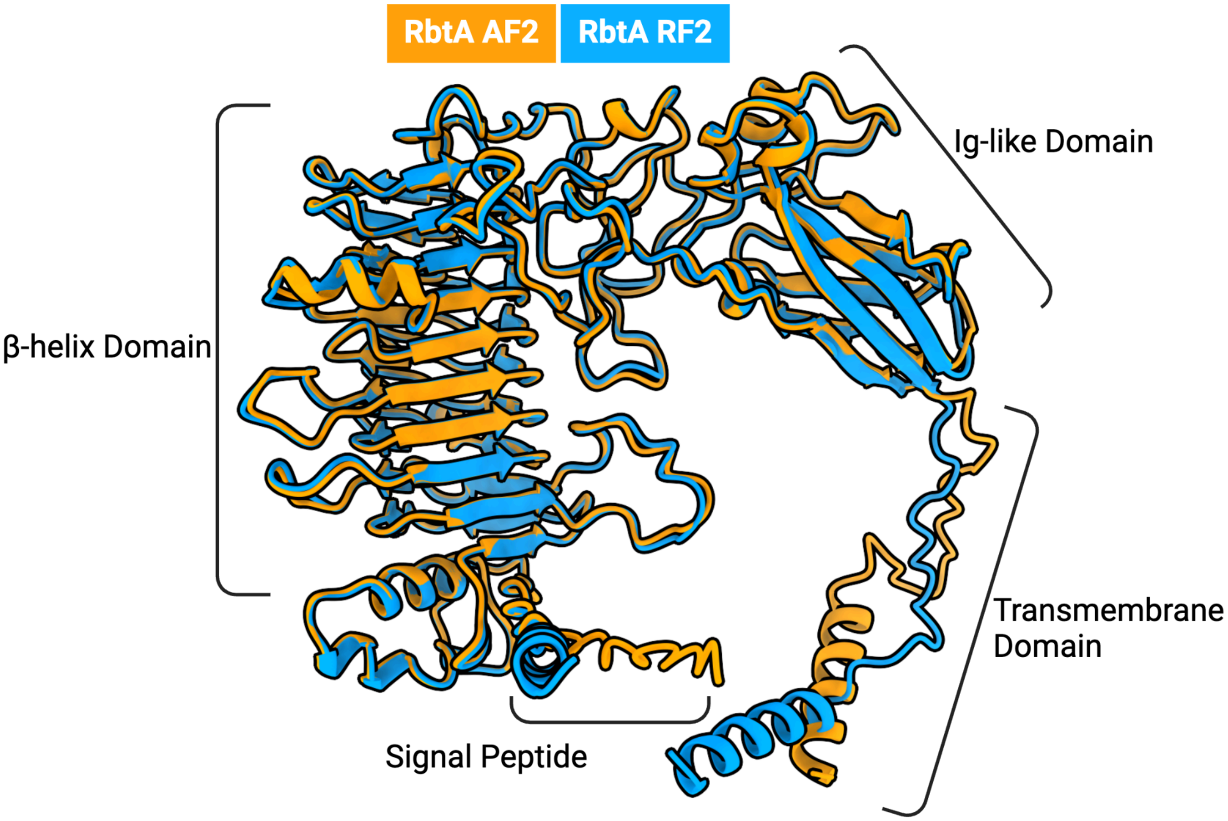

**Figure S16: Alignment of the RbtA structures predicted by RF2 and AF2**

Alignment of RF2 (blue) and AF2 (orange) predictions of full length RbtA from *Acinetobacter baumannii.* The predicted structures from AF2 and RF2 are very similar, with a Cα RMSD excluding the transmembrane domain and signal peptide of 0.5 Å.

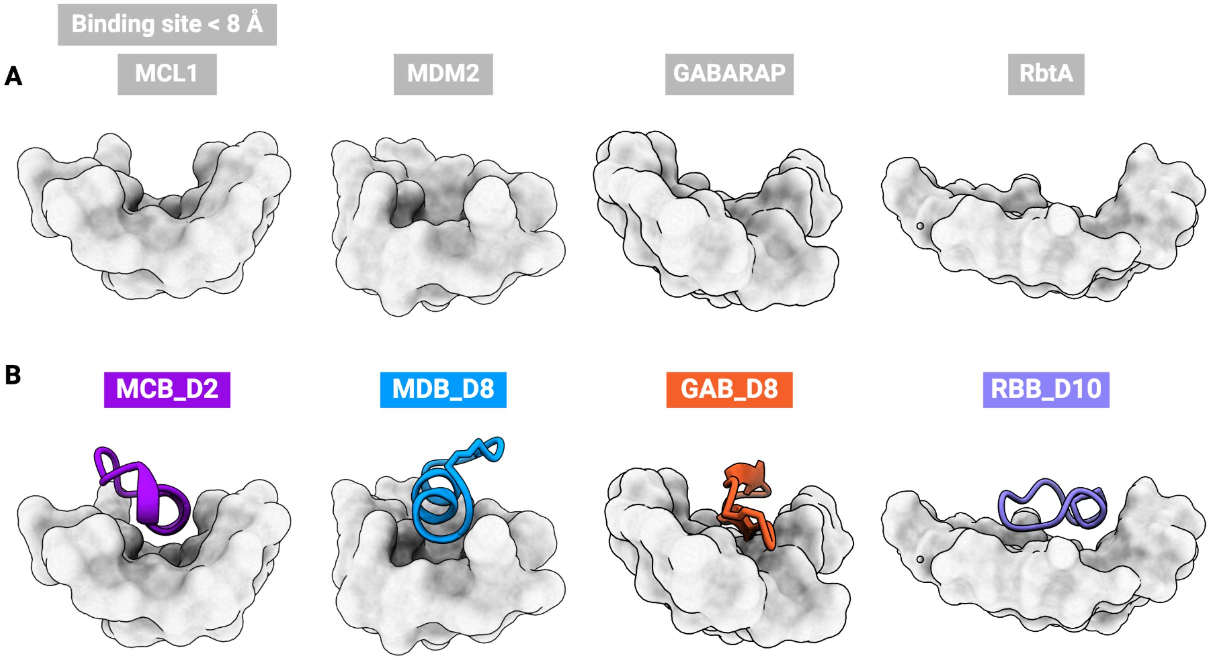

**Figure S17: Macrocycles designed by RFpeptides engage diverse target pockets with with appropriately shape-matched macrocycle comformations**
(A) Atoms within 8 Å of the binder represented as surface for each of the targets described in this work. MCL1 and MDM2 have deep concave pockets, GABARAP has a relatively more open binding site, and RbtA has a very flat surface at the binding site. (B) Computationally-designed macrocycle binders with highest affinity shown in cartoon mode. The deep concave pockets of MCL1 and MDM2 are engaged by helix-containing macrocycles, while the more open, flat surfaces of GABARAP and RbtA are engaged by macrocycles with β-sheet and loop-like conformations, respectively.

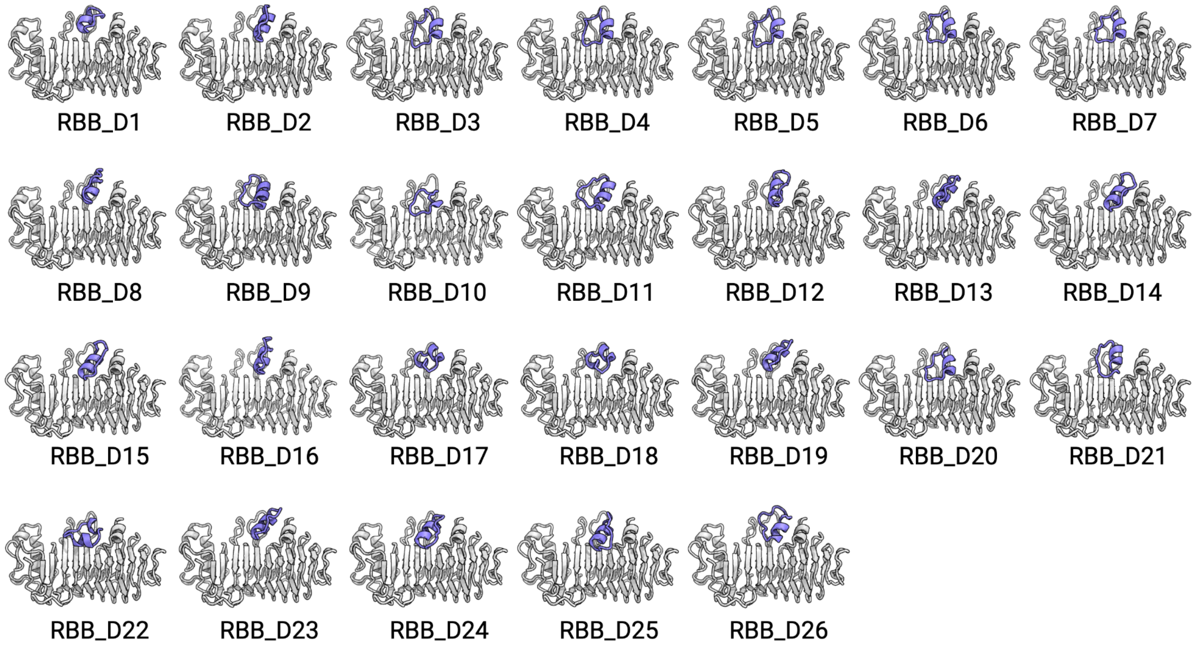

**Figure S18: Structural diversity of the selected RbtA binding designs**
We diffused macrocyclic peptides against RbtA with hotspots L144, F202, F204, Y206, V208, L231, and A269 (based on NCBI GenBank entry PXA52927.1). Of the 80,000 design models, 147 were predicted to fold and bind as designed with high confidence by AfCycDesign (normalized iPAE < 0.4 and 1.5 Å Cα RMSD to the design model). We next used Rosetta to calculate binding affinity (ddG), spatial aggregation propensity (SAP) of the designed macrocycle, and the molecular surface area of the interface contacts (CMS). We generated two lists of candidates, one using Rosetta metrics  ddG < -40, SAP < 35, CMS > 300, and the other of designs with iPAE < 0.28. The two lists were merged leaving 26 designs spanning those with the best Rosetta metrics, those with the best AfCycDesign metrics, and those with both.

**Table S4: Sequences and interacting residues for selected RbtA designs**

| Design | Sequence | Length | Interacting Residues (< 4 Å) |
|---|---|---|---|
| RBB_D1 | SSLPEYVRDVLGV | 13 | A139,S140,K170,A195,S196,K197,D199,R200,F202,F204,L229,D258,S260,E263,Y264,Q267,A269,I270 |
| RBB_D2 | GKPGDSPWEKIESVVFG | 17 | S140,K170,A195,S196,K197,D199,R200,F202,F204,Y206,L229,L231,S260,E263,Y264,Q267,R268,A269,I270 |
| RBB_D3 | PDPAIPENVQKVFKEHML | 18 | A139,S140,S142,L144,K170,T172,A195,K197,R200,F202,F204,Y206,L229,L231,E263,Y264,Q267,R268,A269,I270 |
| RBB_D4 | PDPAFPENVQKVFKDKIE | 18 | A139,S140,S142,L144,K170,L171,T172,A195,K197,R200,F202,F204,Y206,L229,L231,D258,E263,Y264,Q267,R268,A269,I270 |
| RBB_D5 | PDPAFPESIHKVFKDKII | 18 | A139,S140,S142,L144,K170,L171,T172,A195,S196,K197,R200,F202,F204,Y206,L229,L231,D258,E263,Y264,Q267,R268,A269,I270 |
| RBB_D6 | FSKYEPVDDSYPEGIRKV | 18 | A139,S140,S142,K170,L171,T172,A195,S196,K197,R200,T201,F202,F204,Y206,L229,L231,D258,S260,E263,Y264,Q267,A269,I270 |
| RBB_D7 | FEKYDPMDESYPENIRKV | 18 | A139,S140,S142,K170,L171,T172,A195,S196,K197,R200,T201,F202,F204,Y206,L229,L231,F256,D258,S260,E263,Y264,Q267,R268,A269,I270 |
| RBB_D8 | VEGNDKGSDIYEFIRDK | 17 | A139,S140,K170,A195,S196,K197,R200,T201,F202,G203,F204,L229,L231,F256,D258,S260,E263,Y264,Q267,A269,I270 |
| RBB_D9 | VDDAKEQVLAWAKDPTVV | 18 | A139,S140,K170,A195,S196,K197,D199,R200,F202,F204,L229,L231,S260,E263,Y264,Q267,R268,A269,I270 |
| RBB_D10 | KLFGPDPYLPENVQ | 14 | T97,S142,L144,K170,T172,F202,F204,Y206,L229,L231,E263,Y264,Q267,R268,A269,I270 |
| RBB_D11 | SISGEPEVFKKVYSIAGI | 18 | A139,S140,S142,K170,L171,T172,A195,S196,K197,D199,R200,T201,F202,F204,Y206,L229,L231,S260,E263,Y264,Q267,R268,A269,I270 |
| RBB_D12 | VEGKKATSKDPWKVLEDY | 18 | S140,K170,A195,S196,K197,D199,R200,F202,F204,Y206,Q227,L229,L231,F256,D258,S260,E263,Y264,Q267,R268,A269,I270 |
| RBB_D13 | VEASWRAREVLEFLGKS | 17 | S142,K170,L171,T172,A195,S196,K197,D199,R200,F202,F204,Y206,L229,L231,F256,D257,D258,S260,E263,Y264,Q267,R268,A269,I270 |
| RBB_D14 | SVAKEIAEWIGIPSKVPP | 18 | S142,V143,K170,L171,T172,A195,F202,F204,Y206,L229,L231,E263,Y264,V266,Q267,R268,A269,I270 |
| RBB_D15 | SLAKEIADWIGIPSSVPP | 18 | S142,V143,K170,L171,T172,A195,F202,F204,Y206,L |

| | | | |
|---|---|---|---|
| | | | 229,L231,F256,E263,Y264,Q267,R268,A269,I270 |
| RBB_D16 | VRGEKTTPDPWEFIYNL | 17 | A139,S140,K170,A195,S196,K197,S198,R200,T201,F202,F204,L229,L231,F256,D258,S260,E263,Y264,Q267,A269,I270 |
| RBB_D17 | FAPVFAKYDPNLAD | 14 | A139,S140,F141,K170,A195,S196,K197,R200,T201,F202,F204,Y206,L229,L231,D258,S260,E263,Y264,Q267,A269,I270 |
| RBB_D18 | FAPVFAKYDPNLAE | 14 | A139,S140,F141,K170,T172,A195,S196,K197,R200,T201,F202,F204,Y206,L229,L231,D258,S260,E263,Y264,Q267,A269,I270 |
| RBB_D19 | FEASSEVAKQLGEVVG | 16 | S140,K170,A195,S196,K197,R200,F202,F204,L229,F256,D258,S260,E263,Y264,Q267,A269,I270 |
| RBB_D20 | FDKYDPLDDLYPENIRKV | 18 | A139,S140,S142,L144,K170,L171,T172,A195,S196,K197,R200,T201,F202,F204,Y206,L229,L231,F256,D258,S260,E263,Y264,Q267,R268,A269,I270 |
| RBB_D21 | FRLIDKKLGIEDNDPIEA | 18 | A139,S140,S142,K170,L171,T172,A195,S196,K197,D199,R200,F202,F204,Y206,L229,L231,S260,E263,Y264,Q267,A269,I270 |
| RBB_D22 | RKPVEEILKKNEITYEDV | 18 | P95,A139,S140,F141,S142,K170,T172,A195,S196,K197,R200,F202,F204,Y206,L229,L231,D258,S260,E263,Y264,Q267,R268,A269 |
| RBB_D23 | KPPASSEVARKVMEVFG | 17 | S140,K170,A195,S196,K197,D199,R200,F202,F204,L229,L231,F256,D258,S260,E263,Y264,Q267,R268,A269,I270 |
| RBB_D24 | KAKNWFTEQLSEVIPGI | 17 | A139,S142,K170,T172,A195,S196,K197,D199,R200,F202,F204,Y206,L229,L231,F256,S260,E263,Y264,Q267,R268,A269,I270 |
| RBB_D25 | SGNPVLQIAAELDPNVKG | 18 | S140,S142,K170,L171,T172,A195,K197,F202,F204,Y206,L229,L231,T233,F256,S260,E263,Y264,Q267,R268,A269,I270 |
| RBB_D26 | EFARLLAKGDFEGWKDSK | 18 | A139,S140,K170,A195,S196,K197,D199,R200,F202,F204,L229,F256,D258,S260,E263,Y264,Q267,A269,I270 |

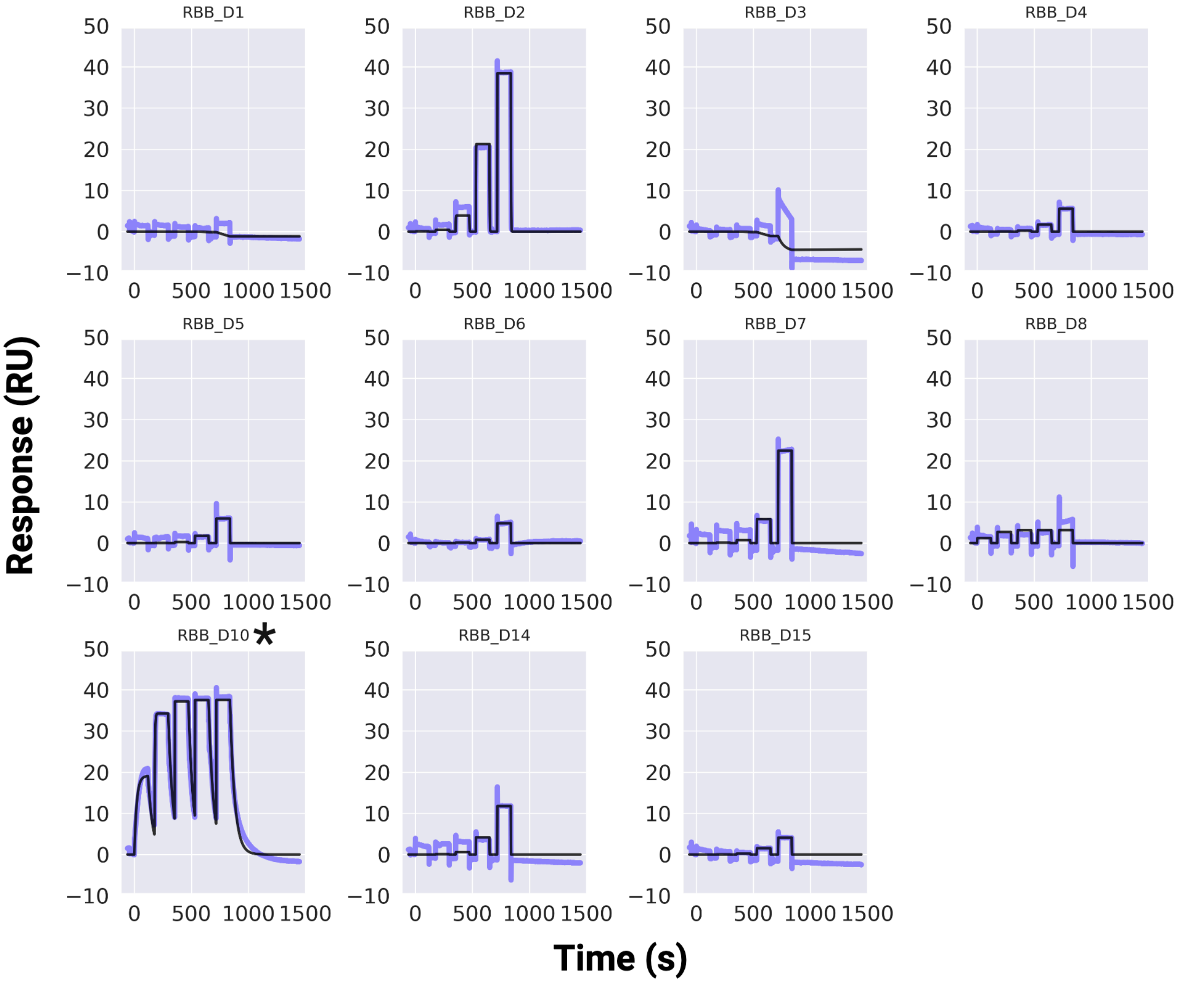

**Figure S19: SPR-based screening of the selected RBB designs against RbtA**

Sensorgrams from an SPR single cycle kinetics screen for the selected RBB designs, applying a 5-point 10-fold dilution with 100 μM as the highest concentration. Four designs (RBB_D2, RBB_D7, RBB_D10, and RBB_D14) show detectable binding at or below 100 μM. RBB_D10 denoted with * chosen for further characterization.

### 2. Materials and Methods

#### 2.1.1. Implementation of the cyclic relative position encodings for RoseTTAFold and RFdiffusion

While conceptually similar to cyclic positional encodings in AfCycDesign[5], our implementation of the cyclic relative position encodings differs in two ways. First, as shown in Figure 1A (e.g., first row), the relative position encoding for a starting residue ‘i’, and its increasingly distant neighbors ‘j’ going in the N→C direction increments by positive 1 at every next position. This positive increment occurs until the difference between i and j is such that j is more than halfway around the peptide from i, at which point the relative position encoding switches sign. With the switched sign, residue j is now seen by residue i to be to the “left” (N-terminal from i) instead of to the right (C-terminal from i). Upon switching the sign, the relative position encoding continues to be incremented by +1, encoding an increasingly close distance from j→i approaching from the left/N-term. When applied to all residues in the forward pass of the network, this manifests a polypeptide chain in which there are no termini and the backbone is cyclized. The second, more practical, difference between the relative position encodings demonstrated in Rettie et. al.[5] and those shown here is the way in which they are introduced to the neural network. In addition to introducing the cyclized offsets into the network pairwise representation, as in Rettie et. al., we include the same offsets into the edge features of the graph fed into the 3D track of RoseTTAFold. This then enables RF’s SE3Transformer to also see that the peptide should be cyclized.

All the required changes for implementing RFpeptides are now incorporated into the RFdiffusion codebase. The latest version of RFdiffusion can be downloaded from https://github.com/RosettaCommons/RFdiffusion.

#### 2.1.2. Monomeric macrocycle prediction with RFpeptides

With cyclic relative positional encoding implemented in RoseTTAFold we used the same set of 80 NMR structures from the ProteinDataBank that was used in our previous work to validate predictions of diverse cyclic structures[5]. PDB entries were identified by searching for monomeric peptides less than 40 amino acids in length, containing only L amino acids, and with CONECT records between the terminal N and C atoms.

1bh4, 1df6, 1hvz, 1jbl, 1qvk, 1qvl, 1r1f, 1ski, 1skk, 1vb8, 1za8, 1znu, 2atg, 2b38, 2eri, 2gj0, 2k7g, 2kch, 2knm, 2kuk, 2kux, 2kvx, 2lam, 2ll5, 2lur, 2lws, 2lwt, 2lwu, 2lwv, 2lye, 2lyf, 2lzi,

2m77, 2m78, 2m79, 2m9o, 2mh1, 2mn1, 2mso, 2mt8, 2mw0, 2n07, 2nb5, 2ndl, 2ndm, 2ndn, 2ns4, 2otq, 2ox2, 2po8, 5h1h, 5h1i, 5kwz, 5kx1, 5wov, 5wow, 6dny, 6pin, 6pio, 6pip, 6u7q, 6u7r, 6u7s, 6wpv, 7f32, 7k7x, 7l53, 7l54, 7l55, 7lhc, 7m25, 7m27, 7m28, 7m29, 7m2a, 7m2b, 7m2c, 7m3u, 7rn3, 7s55

Multiple sequence alignments were generated by

```
python RoseTTAFold/examples/make_protein_msa.sh example.fasta
```

and the resulting `example.a3m` used for the input for prediction using RoseTTAFold

```
python RoseTTAFold/network/predict.py -inputs example.a3m -n_recycles 48 -cyclize
```

The outputs from RoseTTAFold are the highest pLDDT models for each input sequence and MSA. To determine their backbone heavy atom RMSD to the native structure the following RosettaScript was used with Rosetta v2024.16.

---

Example RosettaScript for calculating backbone heavy atom RMSD

---

```
<ROSETTASCRIPTS>
<SIMPLE_METRICS>
      <RMSDMetric name="RMSD"
            rmsd_type="rmsd_protein_bb_heavy"
            use_native="true"
            super="true"/>
</SIMPLE_METRICS>
<MOVERS>
      <PeptideCyclizeMover name="pcm"/>
      <RunSimpleMetrics name="run_metric" metrics="RMSD"/>
</MOVERS>
<PROTOCOLS>
      <Add mover="close" />
```

```
    <Add mover="run_metric" />
</PROTOCOLS>
</ROSETTASCRIPTS>
```

---

In cases where the native structure was an NMR ensemble, all states of the NMR structures were compared to the predicted model and the lowest RMSD was plotted in Supplementary Figure S1.

**2.1.3. Monomeric macrocycle design with RFpeptides**

To design a monomeric ordered macrocycle (i.e., not binder design) we simply add the cyclic positional encoding to RFdiffusion as described in 1.1.1 during an otherwise standard execution of RFdiffusion. Then, we use a sequence design model, such as ProteinMPNN or LigandMPNN to design the sequences. Finally, AfCycDesign[5] is used for structure prediction of the designed peptides.

For the experiment shown in Figures 1 B,C and S2, we first generated 200 backbones for macrocycles of length 8, 10, 12, 14, 16, 18. An example of the execution to generate the peptide backbones is:

```
python rf_diffusion/run_inference.py
inference.output_prefix=/path/to/folder/file_prefix
inference.num_designs=200 contigmap.contigs=[\"8-8\"]
inference.cyclic=True inference.cyc_chains="a" diffuser.T=50
```

The new flags `inference.cyclic=True` and `inference.cyc_chains=”a”` trigger the addition of cyclized offset features to the network. For each of these backbones we then used LigandMPNN with sampling temperature of 0.1 to design 8 sequences per backbone. AfCycDesign was then used to predict the structure of each of these sequences.

**2.1.4. tSNE dimensionality reduction and gaussian mixture model clustering**

For the experiment described in the section 2.1.3, we computed an all-by-all (1200x1200) matrix of TMscores, in which the (i,j) entry is the average of the target- and query-normalized TMscores output from TMalign[6] between peptides i and j. We then used tSNE[7] implemented in Scikit-Learn[8] to reduce the dimensionality of this matrix to 1200x2. We then plotted this matrix, colored by various quantities of interest (e.g., peptide length, scRMSD; Fig S2 D,E). We clustered the points in this matrix using Scikit-Learn's Gaussian mixture model clustering implementation[8]. To ensure meaningful clustering assignments, we computed intra-cluster mean TMscores as a function of the number of clusters being fit to the data (Fig S2 C). Intuitively, as the peptides are split into more and more clusters, the intra-cluster mean TMscore rises.

#### 2.1.5 Measuring self-consistency of designs and MaxCluster clustering

A useful benchmark for polypeptide backbone diffusion models is a measurement of the frequency with which a sampled polypeptide can be assigned an amino acid sequence (usually with ProteinMPNN[9]) that, when "re-folded" using a protein structure prediction network (such as AlphaFold[10], ESMFold[11] or RoseTTAFold[12]), recapitulates the original sampled backbone with high confidence. Such computational filtering criteria have been shown to yield experimentally active designs for a variety of functions[13,14], and is commonly known as the "self-consistency" test. Furthermore, it is useful to quantify the number of *unique* protein or peptide "folds" produced by a generative model that achieve the aforementioned metrics. This additional criteria separates models which consistently produce a single or few unique solutions to a problem from models which produce a multitude of high quality solutions.

To compute this benchmark for RFpeptides, we employed the following pipeline: Using the structures and sequences generated in section 2.1.3 above, we computed the fraction of backbones at each length which have at least 1 of the 8 LigandMPNN sequences re-fold with AfCycDesign to within 2.0 Å RMSD and with at least 0.8 (out of 1.0) pLDDT. In Figure 1C, this statistic is denoted in the panel legend as "all" in blue. Additionally, to quantify the rate of unique structural clusters at each length which are self-consistent, we used MaxCluster[15,16] to structurally cluster all 1200 (6 lengths, 200 structures each) backbones sampled. We then redundancy-reduced the set of passing backbones on a per-length basis according to MaxCluster structural cluster assignments, so as not to double count multiple successes from the same peptide structural family. This new statistic is at most equivalent to the aforementioned "raw" success rate (but likely lower) and is denoted in the Figure 1C panel legend as "unique @ TM 0.5" in orange. It can be interpreted as the probability that a backbone sampled from RFpeptides will be a structurally unique, self-consistent design when sampling a total of 200 backbones at a given length. We used a TMscore cutoff of 0.5 and a hierarchical clustering strategy, executed on the command line as follows:

```
/path/to/maxcluster -l $list_of_pdbs -in -TM -C 2 -Tm 0.5
```

#### 2.2. Macrocycle Binder Design with RFpeptides

Generally, designing macrocyclic binders to selected targets with RFpeptides is a four-step process. First, we generate macrocyclic peptide backbones with diverse sizes (e.g., sampling a number of designs from length 8 to 18 uniformly) against the selected target protein. This backbone process can also be guided towards a specified list of 'hotspot' residues on the target. Next, amino acid sequences are designed for each generated backbone using iterative rounds of ProteinMPNN[9] and Rosetta FastRelax[17,18]. Third, the designed sequences are re-predicted using AfCycDesign and RF2 to confirm if the designed sequences fold and bind in the designed conformations. Finally, the downselected designs from AfCycDesign and RF2[5,19] are further filtered using Rosetta-based interface metrics.

##### 2.2.1 Macrocycle Backbone Generation

To prepare the target structure for macrocycle backbone generation, the PDB files corresponding to the selected target were downloaded from the Protein Data Bank and stripped of all water and ligands, leaving just the target protein atoms. For medium to large-sized target proteins (> 300 amino acids), the target protein can also be truncated to keep just the relevant target domains to reduce the computational costs. An example script for macrocycle backbone generation is shown below. It generates 400 macrocyclic peptides (controlled by the `inference.num_designs`) ranging from 12 to 18 amino acids against residues 1-113 of the target protein defined as chain A. The `contigmap.contigs` option defines the size range as 12-18 residues for the diffused cyclic peptide against residues 1-113A of a target protein as chain A from `inference.input_pdb`. The `inference.cyclic` option implements cyclic relative positional encoding and `inference.cyc_chains='a'` applies it just to the diffused peptide that will be chain A in the output pdb file. The number of time steps are controlled using `diffuser.T=50` setting them to 50. The `ppi.hotspot_res` option defines the target hotspot residues to guide the macrocycle generation. In the example below, the macrocycles are generated to bind around the residue numbers 68 and 70 of the target chain A. This line can be omitted if no hotspots are to be used. The final output will have a diffused cyclic peptide 12-18 residues as chain A and the residues 1 to 113 of the input target protein shifted to chain B.

---

Example script for generating macrocyclic binders against a target protein

```
prefix=./outputs/diffused_binder_cyclic
pdb='./target.pdb'
num_designs=400
script="rf_diffusion/run_inference.py"
python $script --config-name base \
      inference.output_prefix=$prefix \
      inference.num_designs=$num_designs \
      contigmap.contigs=[\'12-18,0\ A1-113,0\'] \
      inference.input_pdb=$pdb \
      inference.cyclic=True \
      diffuser.T=50 \
      +inference.cyc_chains='a' \
      +ppi.hotspot_res=[\'A68\',\'A70\'] \
```

For MCL1 and MDM2 designs, we generated 10,000 backbones without defining any hotspot residues. For targeting GABARAP and RbtA, we generated 20,000 backbones with hotspots specified. For designing against GABARAP, we defined six hotspot residues: Tyr51, Leu52, Lys50, Lys48, Phe62, and Leu65 (residue numbering the same as PDB ID: 7ZKR[20]). For targeting RbtA, we used the following seven hotspot residues: Leu144, Phe202, Phe204, Tyr206, Val208, Leu231, and Ala269 (residue numbering based on NCBI GenBank entry PXA52927.1). We explored macrocycles of length 16 residues against MCL1, lengths 16-18 residues for MDM2, and 12-18 for GABARAP and RbtA. Example scripts and files are available as Supplementary files.

#### 2.2.2. Sequence Design

Amino acid sequences for the designed backbones were designed using four iterative rounds of ProteinMPNN[9] followed by Rosetta FastRelax[18]. ProteinMPNN can be downloaded from https://github.com/dauparas/ProteinMPNN. An example Rosetta FastRelax script is shown below and is also available in Supplementary Files. This example uses the output from the example script for generating macrocyclic binders against a target protein, `diffused_binder_cyclic_1.pdb`

.

We used the following ProteinMPNN options:

```
--pdb_path ./diffused_binder_cyclic_1.pdb
--pdb_path_chains A
--temperature 0.0001
--backbone_noise 0
--omit_AAs C
--num_seq_per_target 1
--path_to_model_weights vanilla_model_weights/v_48_020.pt
```

We chose to omit cysteine from our designs to streamline synthesis and validation of the peptides as disulfides or singular cysteines would require additional steps in synthesis. After designing the sequence with ProteinMPNN, the input model was updated with the new sequence, and the structure was minimized with FastRelax. During the relax step, distance, angle, and dihedral constraints were applied to maintain the terminal bond using Rosetta PeptideCyclizeMover as described previously[21,22]. The relax step was implemented as a PyRosetta4[23] (Version 2022.41) script that imports the PeptideCyclizeMover and FastRelax as XmlObjects from the following RosettaScripts.

---

An example RosettaScripts fast_relax.xml used for PeptideCyclizeMover and FastRelax movers

---

```
<ROSETTASCRIPTS>
<SCOREFXNS>
<ScoreFunction name="beta" weights="beta_nov16">
        <Reweight scoretype="metalbinding_constraint" weight="1.0" />
        <Reweight scoretype="arg_cation_pi" weight="3" />
                <Reweight scoretype="approximate_buried_unsat_penalty" weight="5”/>
                <Set approximate_buried_unsat_penalty_burial_atomic_depth="3.5" />
                <Set approximate_buried_unsat_penalty_hbond_energy_threshold="-0.5" />
                <Reweight scoretype="coordinate_constraint" weight="1" />
                <Reweight scoretype="atom_pair_constraint" weight="1" />
                <Reweight scoretype="dihedral_constraint" weight="1" />
                <Reweight scoretype="angle_constraint" weight="1" />
        </ScoreFunction>
<RESIDUE_SELECTORS>
        <Chain name="chA" chains="A"/>
        <Neighborhood name="int" selector="chA" distance="8"/>
        <Not name="target" selector="chA"/>
```

```
        <And name="target_int" selectors="target,int" />
        <Not name="not_target_int" selector="target_int" />
        <And name="non_interface_target" selectors="not_target_int,target"/>
</RESIDUE_SELECTORS>
<TASKOPERATIONS>
        <OperateOnResidueSubset name="move_interface" selector="target_int" >
                <RestrictToRepackingRLT/>
        </OperateOnResidueSubset>
        <OperateOnResidueSubset name="hold_non_interface" selector="non_interface_target" >
                <PreventRepackingRLT/>
        </OperateOnResidueSubset>
</TASKOPERATIONS>
<MOVERS>
        <PeptideCyclizeMover name="pcm" residue_selector="chA"/>
        <FastRelax name="full_relax_complex"
                repeats="3"
                scorefxn="beta"
                min_type="dfpmin_armijo_nonmonotone"
                ramp_down_constraints="false"
                task_operations="move_interface,hold_non_interface">
                <MoveMap name="specifics">
                        <Jump number="1" setting="true" />
                        <ResidueSelector selector="non_interface_target" chi="0" bb="0" />
                        <ResidueSelector selector="target_int" chi="1" bb="0" />
                </MoveMap>
        </FastRelax>
</MOVERS>
<PROTOCOLS>
        <Add mover="pcm" />
        <Add mover="full_relax_complex"/>
</PROTOCOLS>
<OUTPUT />
</ROSETTASCRIPTS>
```

---

---

PyRosetta script to relax the macrocycle-bound target protein

---

```
from pyrosetta import *
init('-beta_nov16')

xml = 'fast_relax.xml'
```

```
objs = protocols.rosetta_scripts.XmlObjects.create_from_file(xml)
fr = objs.get_mover('full_relax_complex')
pcm = objs.get_mover('pcm')
pose = pose_from_pdb('diffused_binder_cyclic_1_mpnn.pdb')
pcm.apply(pose)
fr.apply(pose)
pcm.apply(pose)
pose.dump_pdb('diffused_binder_cyclic_1_mpnn1.pdb')
```

---

After relaxing the design with the ProteinMPNN generated sequence, the new model was designed again using ProteinMPNN, and this cycle was repeated. Together, this process generated four sequences for each backbone generated by the diffusion step. Overall, for MCL1 and MDM2, we generated 39,860 and 40,000 designs, respectively. 80,000 designed sequences were generated for GABARAP and RbtA.

**2.3. Target-macrocycle complex structure prediction using AfCycDesign**

We used AfCycDesign version 1.1.1 to predict the structure of designed cyclic peptides in complex with their target. An example script to run AfCycDesign for macrocycle-bound target is described below and can also be run using ColabDesign at https://colab.research.google.com/github/sokrypton/ColabDesign/blob/main/af/examples/af_cyc_design.ipynb

In the example below, the target protein is chain A. No templates were used for the designed macrocycle. The structure of the target chain in the input model was used as a template while predicting the co-complex structure. The outputs from this step is a pdb file of the predicted binding mode of the designed peptide, and a score file score.sc containing the interface predicted aligned error (iPAE) and Cα RMSD of the peptide to the design model. Normalized iPAE (iPAE/32) cutoff values for filtering were chosen based on the iPAE distribution per target. 0.3 and 0.2 for MDM2 and MCL1, 0.13 and 0.4 for GABARAP and RbtA respectively. iPAE values over 0.4 were often nonoptimal interfaces or the peptide not interacting with the target in the designed conformations.

---

**An example script for target-macrocycle complex structure prediction using AfCycDesign**

```
pdb = 'diffused_binder_cyclic_1_mpnn1'
model = mk_afdesign_model('binder')
model.prep_inputs(f'{pdb}.pdb',
     binder_chain='B',
     target_chain='A',
     use_binder_template=False,
     use_multimer=True,
     use_initial_guess=True)
add_cyclic_offset(model, offset_type=2)
model.set_seq(mode='wildtype')
model.set_opt(num_recycles=1)
model.predict(models=[0,1], verbose=False)
model.save_pdb(f'diffused_binder_cyclic_1_mpnn1_prediction.pdb')
rmsd = model.aux['losses']['rmsd']
ipae = model.aux['all']['losses']['i_pae'][0]
with open(f'score.sc','a') as outfile:
     outfile.write(f'{pdb},{ipae},{rmsd}\n')
```

#### 2.4. Filtering designs based on physics-based metrics of interface quality

After filtering design models based on the confidence metrics from DL-based structure prediction of design models (AfCycDesign and RF2), we used Rosetta (version 2021.12) to calculate the binding affinity (ddG), spatial aggregation propensity (SAP), and the contact molecular surface area (CMS) of the designed target macrocycle complexes. In the example below, target is defined as chain A and macrocycle is defined as chain B.

**An example script for calculating physics-based interface metrics using Rosetta**

```
<ROSETTASCRIPTS>
```

```
<SCOREFXNS>
	<ScoreFunction name="sfxn_cart" weights="beta_nov16_cart" >
		<Reweight scoretype="coordinate_constraint" weight="1" />
		<Reweight scoretype="atom_pair_constraint" weight="1" />
		<Reweight scoretype="dihedral_constraint" weight="1" />
		<Reweight scoretype="angle_constraint" weight="1" />
	</ScoreFunction>
</SCOREFXNS>
<RESIDUE_SELECTORS>
	<Chain name="chainA" chains="1"/>
	<Chain name="chainB" chains="2"/>
	<Neighborhood name="interface_chA" selector="chainB" distance="14.0" />
	<Neighborhood name="interface_chB" selector="chainA" distance="14.0" />
	<And name="AB_interface" selectors="interface_chA,interface_chB" />
	<Not name="Not_interface" selector="AB_interface" />
</RESIDUE_SELECTORS>
<TASKOPERATIONS>
	<ProteinInterfaceDesign name="pack_long"
		design_chain1="0"
		design_chain2="0"
		jump="1"
		interface_distance_cutoff="15"/>
	<OperateOnResidueSubset name="restrict_to_interface" selector="Not_interface">
		<PreventRepackingRLT/>
	</OperateOnResidueSubset>
</TASKOPERATIONS>
<MOVERS>
	<PeptideCyclizeMover name="pcm" residue_selector="chain%%chain%%"/>
	<TaskAwareMinMover name="minimize_interface"
		scorefxn="sfxn_cart"
		tolerance="0.01"
		cartesian="true"
		task_operations="restrict_to_interface"
		jump="0" />
	<TaskAwareMinMover name="min"
		scorefxn="sfxn"
		bb="0"
		chi="1"
		task_operations="pack_long" />
<FILTERS>
	<Ddg name="ddg"
		threshold="50"
		jump="1"
		repeats="5"
		repack="1"
		relax_mover="min"
		confidence="0"
		scorefxn="sfxn"
```

```
                extreme_value_removal="1" />
        <ContactMolecularSurface name="contact_molecular_surface"
                distance_weight="0.5"
                target_selector="chainA"
                binder_selector="chainB"
                confidence="0" />
<SIMPLE_METRICS>
        <SapScoreMetric name="sap_score" score_selector="chainB" />
</SIMPLE_METRICS>
<PROTOCOLS>
        <Add mover="pcm" />
        <Add mover="minimize_interface" />
        <Add mover="pcm" />
        <Add filter="ddg" />
        <Add metrics="sap_score" />
        <Add filter="contact_molecular_surface" />
    </PROTOCOLS>
</ROSETTASCRIPTS>
```

---

**2.5. Structure-based clustering for binder design campaigns**

To cluster structurally similar designed macrocycle binders, we first extracted them from their complexes. Then, each design was circularly permuted to generate all possible sequence representations of the macrocycle. At an arbitrarily selected example in the list of all designs to be clustered, C$\alpha$ RMSD was calculated against all members of the list and their cyclic permutations. If C$\alpha$ RMSD was less than 0.5 Å then that member and its permutations were removed from the list and clustered with the design currently looping through the list. This process was repeated until all members were placed into their respective clusters.

**2.6. Prediction of RbtA Structure**

The structure of RbtA was predicted using both RoseTTAFold2 and AlphaFold2. For both prediction methods, MSAs were originally generated following the procedure from the original RoseTTAFold publication[12], where hhblits was used to search both the Uniprot and BFD protein sequence databases at increasing e value cutoffs until at least 10000 sequences are found[10,24]. The resulting MSAs were then used as inputs to RoseTTAFold2 and AlphaFold2 (using CollabFold)[10,19,25]. In both cases, 5 models were generated (all 5 AlphaFold2 weights were used, and the single RoseTTAFold weights were used with 5 different random starting configurations); the model of the five with highest predicted accuracy (pLDDT) was chosen as the design target.

Both AlphaFold2 and RoseTTAFold2 yielded models with pLDDTs > 0.9, indicating very high confidence; the models were in very good agreement with one another.

### 3. Experimental Methods

#### 3.1. Peptide Synthesis

Macrocyclic peptides described here were either purchased from Wuxi AppTec at greater than 90% purity or synthesized in-house using the Fmoc-based solid-phase peptide synthesis. Peptides were typically synthesized on preloaded 2-chlorotrityl chloride (CTC) resin. The resin was swollen in dichloromethane (DCM) followed by iterative deprotection with 20% piperidine in dimethyl formamide (DMF) and coupling with either O-(benzotriazol-1-yl)-N,N,N′,N′-tetramethyluronium hexafluorophosphate (HBTU, Sigma) or 7-azabenzotriazol-1-yloxy)trispyrrolidinophosphonium hexafluorophosphate (PyAOP, Novabiochem), and N,N-Diisopropylethylamine (DIEA, Sigma). The linear peptides were cleaved from the resin using either 2% TFA in DCM or 20% 1,1,1,3,3,3-hexafluoro-2-propanol (HFIP, Oakwood Chemical) in DCM. The solvent was removed by rotary evaporation and linear protected peptides were cyclized in either DCM, DMF, or a mixture of both depending on the solubility of the peptide, using 2 equivalents of PyAOP and 5 equivalents of DIEA overnight. The protecting groups are removed using a cocktail of 95:2.5:2.5, trifluoroacetic acid:water:triisopropylsilane for 2.5 hours. The crude peptides were precipitated using cold diethyl ether. The precipitate containing the crude cyclization reaction was dissolved in a mixture of water and acetonitrile for purification using reverse-phase high-performance liquid chromatography (RP-HPLC). Peptide identities were confirmed by mass spectrometry. Purities for all synthesized and tested macrocyclic peptides are also summarized in tables S7-10. The mass spectrogram and analytical LC chromatogram for all purified peptides are shown in section 4.

#### 3.2. Protein Expression and Purification

##### 3.2.1. MDM2 and MCL1

The amino acid sequences of MCL1 (PDB ID: 2PQK[1]) and MDM2 (PDB ID: 4HFZ[26]) were retrieved from the Protein Data Bank. The optimized genes were then cloned into a Novogen pRSF-DUET plasmid (Sigma, Cat No. 71341-3), incorporating a 6xHistidine-tag at the N-terminus, followed by an Avi-tag and a Tobacco Etch Virus (TEV) protease cleavage site. The resulting constructs were codon-optimized for *E. coli* expression and synthesized by Genscript (Piscataway NJ, USA). For propagation, the plasmids were transformed into *E. coli* NEBα cells

(NEB, Cat No. C2987), and for protein expression, into *E. coli* BL21 DE3 cells (NEB, Cat No. C2527). A single sequence verified colony was cultured in 50 ml of Kanamycin (50 µg/ml) selective Luria Broth (LB) media. This culture was incubated at 37°C with shaking at 200 rpm for 16 hours overnight. Subsequently, 50 optical density at 600 nanometers ($OD_{600}$) units of the overnight culture were transferred to 1 L of fresh Kanamycin (50µg/ml) selective LB media. The culture was grown at 37°C with shaking at 200 rpm for 2 hours (until it reached an $OD_{600}$ of 0.4-0.5), at which point the temperature was decreased to 20°C. The culture was grown until reaching an $OD_{600}$ of 0.7-0.8, and protein expression was induced by adding 1 mM Isopropyl β-D-1-thiogalactopyranoside (IPTG) and left to grow overnight for 14 h.

Cells were harvested by centrifugation at 5,000 x g for 10 minutes at 4°C, resulting in a cell pellet with a density of 5 g/L. The pellet was immediately flash-frozen and stored at -20°C for later use. For lysis, the pellet was thawed on ice and resuspended in 5 ml of lysis buffer per gram of pellet. This lysis buffer contained 50 mM Tris-HCl, 300 mM NaCl, and 10 mM Imidazole, and was supplemented with 1X BugBuster Protein Extraction Reagent (Sigma-Aldrich, Cat No. 70921), 200 µg/ml Lysozyme (Sigma-Aldrich, Cat No. L6876), 25 U/ml Benzonase Nuclease (Sigma-Aldrich, Cat No. E8263), and 1X cOmplete™ EDTA-free Protease Inhibitor Cocktail (Sigma-Aldrich, Cat No. 11836170001). The buffer was filter-sterilized using a 0.2 µm filter prior to the addition of Benzonase, mixed by inversion, and kept on ice until use. Cells were completely resuspended in the lysis buffer using a homogenizer at low speed and incubated for 30 minutes at room temperature (22-25°C). Following incubation, the suspension was sonicated using a Q500 Sonicator equipped with a 4-tip probe. Sonication was conducted for 2-3 minutes using pulses of 10-15 seconds on followed by 10-15 seconds off, at 70% amplitude. The lysate was clarified by centrifugation at 16,000 x g for 20 min.

Ni-NTA agarose resin (QIAGEN, Cat No. 30210) was equilibrated with 20 column volumes (CV) of ultrapure water, followed by 20 CV of equilibration buffer (50 mM Tris-HCl, 300 mM NaCl, 10 mM Imidazole). 4 ml of 50% resin suspended in equilibration buffer were used to bind His-tagged proteins from 25 ml of clarified lysate. All immobilized metal affinity chromatography (IMAC) steps were conducted at 4°C. The lysate-resin mixture was incubated for 60 minutes on a rotary shaker set to a slow speed. After incubation, the resin was transferred to a 20 ml gravity column and allowed to completely settle. The resin was first washed with 20 CV of wash buffer 1 (20 mM Tris-HCl, 250 mM NaCl, 10 mM Imidazole, 5 mM beta-mercaptoethanol), followed by another 20 CV of wash buffer 2 (20 mM Tris-HCl, 500 mM NaCl, 35 mM Imidazole). The bound proteins were then eluted with 8 ml of elution buffer (20 mM Tris-HCl, 250 mM NaCl, 350 mM Imidazole, 2 mM Dithiothreitol). Aliquots of the eluate were collected and analyzed using SDS-PAGE gels.

The eluate was loaded onto a pre-equilibrated Superdex 75 10/300 GL column (25 mM Tris-HCl, 250 mM NaCl, 2 mM Dithiothreitol) and run at a flow rate of 0.6 ml/min using an ÄKTA pure system for size exclusion chromatography (SEC). 1 ml fractions were collected from the 8 to 16 ml elution volume, and those corresponding to peaks in the absorbance at 280 nm between 10 and 13 ml elution volume were assessed with SDS-PAGE gels. Fractions confirming the expected molecular weight were pooled and concentrated by centrifugation at 4,000 x g for 30 min at 4°C using Amicon Ultra-4 concentrators with a 3 kDa cutoff (Millipore SIGMA, Cat No. UFC800308) to a final volume of 500 µl. The identity and purity of the eluted proteins were confirmed via mass spectrometry using an Agilent 6230 LC/MS Time-of-Flight (TOF) system.

Verified protein samples were processed for further applications: biotinylation for surface plasmon resonance (SPR) analysis or tag removal via TEV protease cleavage for crystallography. Biotinylation was performed using the BirA biotin-protein ligase standard reaction kit (Avidity LLC, BirA-500) according to the manufacturer's recommended conditions. The reaction was carried out at 4°C overnight on a slowly shaking platform. For TEV protease cleavage, the proteins were treated with a 25:1 protein to TEVd enzyme ratio[27]. Similarly, the mixture was incubated at 4°C overnight on a slowly shaking platform. Following these treatments, samples underwent a cleanup step using 1 ml Ni-NTA resin per 20 mg of protein. The resin was pre-equilibrated with 10 CV of ultra-pure water and 10 CV of a buffer containing 25 mM Tris-HCl, 250 mM NaCl, and 10 mM Imidazole. The pre-equilibrated resin was added to the protein mixture and incubated for 30 minutes on a rolling platform at 4°C. Subsequently, the mixtures were filtered through a 0.45 µm PVDF centrifugal filtering unit to remove the Ni-NTA bound substrates. The eluate was collected and dialysed in 2 L of 25 mM Tris-HCl, 250 mM NaCl, 2 mM Dithiothreitol using a Slide-A-Lyzer™ G3 Dialysis Cassettes, 3.5K molecular weight cut-off (Thermo Scientific, Cat No. A52966) overnight for 18h at 4°C stirring. The dialysed protein was concentrated to 0.2 - 0.5 ml (as required for downstream assays), using the Amicon ultra concentrators (as above), aliquoted, and flash-frozen. Fractions were analyzed by mass-spectroscopy for the efficacy of the biotinylation and TEV protease cleavage treatments, as previously described.

#### 3.2.2. GABARAP for Surface Plasmon Resonance

A synthetic cDNA was designed based on the amino acid sequence of GABARAP (UniProt: O95166) and optimized for expression in *E. coli* using Benchling software. The construct was devised to include an N-terminal Avi-tag and TEV protease cleavage site and was cloned into the Novogen pET-50b(+) plasmid. This plasmid configuration introduced a tandem arrangement of protein tags at the N-terminus: a 6xHistidine tag, followed by a NusA solubility tag, another 6xHistidine tag, and a Human Rhinovirus (HRV) 3C protease cleavage site. Therefore, the final construct sequence was as follows: 6xHis-NusA-6xHis-HRV3C-AviTag-TEV-Gabarap. NusA was specifically chosen as a solubility tag due to its known effectiveness in enhancing protein solubility in *E. coli*[28,29]. The construct was synthesized and cloned by Genscript (Piscataway NJ, USA).

As described above for MCL1 and MDM2 protein expression, the plasmids were introduced into *E. coli* NEBα cells and BL21 DE3 cells. A single sequence verified colony was cultured in 50 ml of Kanamycin (50 µg/ml) selective LB media for 16 hours at 37°C, shaking at 200 rpm. 50 optical $OD_{600}$ units of this culture were transferred to 1 L of fresh Kanamycin (100 µg/ml) selective Autoinduction media (TBM-5052: 1.2% (w/v) tryptone, 2.4% (w/v) yeast extract, 0.5% (v/v) glycerol, 0.05% (w/v) D-glucose, 0.2% (w/v) D-lactose, 25 mM $Na_2HPO_4$, 25 mM $KH_2PO_4$, 50 mM $NH_4Cl$, 5 mM $Na_2SO_4$, 2 mM $MgSO_4$, 10 µM $FeCl_3$, 4 µM $CaCl_2$, 2 µM $MnCl_2$, 2 µM $ZnSO_4$, 400 nM $CoCl_2$, 400 nM $NiCl_2$, 400 nM $CuCl_2$, 400 nM $Na_2MoO_4$, 400 nM $Na_2SeO_3$, 400 nM $H_3BO_3$). The culture was grown at 37°C with shaking at 200 rpm for 2 hours, at which point the temperature was decreased to 22°C and the culture left to grow for 16 hours.

Cells were harvested, lysed, and purified following the protocol outlined earlier for MCL1 and MDM2, with the following modifications: The cultures yielded a cell pellet amounting to 15 g/L. Lysis was completed using an IKA T18 Microfluidizer at 450 psi, followed by lysate clarification by centrifugation at 16,000 x g for 15 minutes. All IMAC steps were conducted at 22°C, except for the incubation of the lysate-resin mixture, which was performed at 4°C. Proteins bound to the resin were eluted with 5 ml of elution buffer (50 mM Tris-HCl (pH 8), 250 mM NaCl, and 300 mM imidazole). Size exclusion chromatography (SEC) was then performed using a Superdex 200 Increase 10/300 GL column (Cytiva) equilibrated with TBS (50 mM Tris-HCl (pH 8), 250 mM NaCl). Fractions confirmed by SDS-PAGE were pooled and concentrated using Amicon Ultra-15 concentrators with a 30 kDa cutoff (Millipore SIGMA, Cat No. UFC9030) to a final volume of 1 ml. Downstream processing for SPR analysis was performed as described previously, with the following modification: for biotinylation, the protein was first cleaved using HRV 3C protease using the reagents and protocol provided by the Pierce™ HRV 3C Protease Solution Kit (Thermo Scientific, Cat No. 88946). The digested samples were subsequently purified and verified, as outlined in earlier sections.

#### 3.2.3 GABARAP and GABARAPL1 for Crystallography

GABARAP and GABARAPL1 were expressed as glutathione S-transferase (GST) fusion proteins after transforming *E. coli* BL21 (DE3) T1 cells with pGEX4T2-GABARAP and pGEX4T2-GABARAPL1 plasmids, respectively. Bacteria were cultivated in LB media containing 100 µg/ml Ampicillin; gene expression was induced with 1 mM IPTG at an $OD_{600}$ of 0.6–0.8 and allowed to proceed for 20 hours at 25°C. Afterwards cells were harvested by centrifugation at 3000 × g for 30 min at 4°C. The bacterial pellet was washed with phosphate buffered saline (PBS; 137 mM NaCl, 2.7 mM KCl, 1.8 mM $KH_2PO_4$, 10 mM $Na_2HPO_4$) and resuspended in lysis buffer (PBS supplemented with 5% (v/v) glycerol, 0.01% (v/v) beta-mercaptoethanol, 10 µg/ml DNase (AppliChem, Cat No. A3778), cOmplete™ EDTA-free Protease Inhibitor Cocktail (Roche, Cat No. 11836170001)) before application to the cell disruptor (Constant Systems, model TS1.1) for 3 cycles with 1.9 kbar at 4°C. Lysates were cleared by centrifugation at 4°C with 45,000 × g for 45 min. The GST fusion proteins were purified from the supernatant by affinity chromatography using Glutathione Sepharose 4B (Cytiva, Cat No. 1705605). Cleavage with thrombin (Sigma-Aldrich; Cat No. 1.12374) during dialysis against 10 mM Tris-HCl, 150 mM NaCl, pH 7.0 at 4°C overnight yielded 119-aa proteins carrying an N-terminal glycine-serine extension in addition to the native residues of GABARAP/GABARAPL1. Subsequently, samples were applied to a Hiload 26/60 Superdex 75 preparatory grade size exclusion column (GE Healthcare) equilibrated with 10 mM Tris-HCl, 150 mM NaCl, pH 7.0. Protein purity was assessed by SDS-PAGE and Coomassie staining. Fractions containing the eluted proteins were concentrated to 3–5 mg/ml using Vivaspin 20 concentrators, 3 kDa cut-off (Sartorius), flash-frozen in liquid $N_2$ and kept at -80°C for long-term storage.

#### 3.2.4. RbtA β-helix domain

For heterologous expression of β-helix domain of RbtA (residues A20-I459) in *E. coli*, the gene was amplified and fused with a SNAC tag (GSHHWGS) at the C-terminus using the primers (forward: GCTGCCCAGCCGGCGATGGCCATGGGCGCTGATATTGAAGTCACAACTAC; reverse:

CAGTGGTGGTGGTGGTGGTGCTCGAGGCTGCCCCAATGATGGCTGCCGATATATTCAATTG CGCCTAAAT)[30]. The fragment was inserted to NcoI-digested and XhoI-digested pET-22b(+) by Gibson assembly to generate a construct with C-terminal 6xHistidine fusion. The construct was confirmed by sequencing and transformed into *E. coli* Rosetta (DE3).

To purify the β-helix domain of RbtA, overnight culture of Rosetta (DE3) carrying the construct was back-diluted 1:300 in 2xYT broth and grown at 37°C with shaking at 200 r.p.m. until the $OD_{600}$ reached 0.4. The incubation temperature was reduced to 18°C, IPTG was added to a final concentration of 0.3 mM and the culture was incubated for a total of 18 hours. Cells were then collected by centrifugation and resuspended in lysis buffer containing 200 mM NaCl, 50 mM Tris-HCl pH 7.5, 10% glycerol (v/v), 5 mM imidazole, 0.5 mg/ml lysozyme and 1 mU benzonase. Cells were then lysed by sonication and cellular debris was removed by centrifugation at 35,000 x g for 30 minutes at 4°C. The protein was purified from lysates using a 1 ml HisTrap HP column on an AKTA fast protein liquid chromatography (FPLC). Column bound protein was eluted using a linear imidazole gradient from 5-500 mM. Protein purity was assessed by SDS-PAGE and Coomassie staining. The fractions with high purity were concentrated using 30 kDa cut-off Amicon filter and then further purified by FPLC using a HiLoad 16/600 Superdex 200 pg column (GE Healthcare) equilibrated with sizing buffer (500 mM NaCl, 50 mM Tris-HCl pH 7.5 and 10% glycerol (v/v)). The fractions with high purity were concentrated and used for evaluation of macrocyclic binders or determination of X-ray structure.

For determination of the X-ray crystal structure of RbtA, the C-terminal 6xHistidine tag was removed by chemical cleavage at the SNAC tag. In Brief, the buffer of the concentrated protein was exchanged to cleavage buffer (0.1 M N-cyclohexyl-2-aminoethanesulfonic acid, 0.1 M NaCl, 0.1 M acetone oxime, 5 mM Fos-choline-12 pH 8.6). The protein solution was diluted to 1 mg/ml, followed by the addition of 1 mM TCEP and 1 mM $NiCl_2$. The mixture was vortexed and incubated at room temperature for 16 hours. The precipitation was removed by centrifugation at 35,000 x g for 30 minutes at 4°C. The supernatant was concentrated and exchanged to Tris buffer (50 mM Tris-HCl pH 7.5, 200 mM NaCl). The protein solution was incubated with 1 ml bed volume of Ni-NTA beads to extract the cleaved 6xHistidine tag. The resulting fraction was concentrated and then further purified by FPLC using a HiLoad 16/600 Superdex 200 pg column.

**3.3. Determination of Binding Affinity by Surface Plasmon Resonance**

Surface plasmon resonance experiments were performed using a Cytiva Biacore 8K in HBS-EP+ buffer from Cytiva. Measurements were obtained by immobilization of biotinylated target protein using the Biotin Capture Kit from Cytiva. Binding screens were performed by single-cycle kinetics (SCK) experiments using the standard protocol in the Biacore 8K control software at 30 μl/min with serial injections of 10 nM, 100 nM, 1 μM, 10 μM and 100 μM, association time of 60 seconds and dissociation time of 120 seconds. For MCL1 designs a dissociation time of 150 seconds was used. To evaluate the affinity of successful designs, a 9-point SCK experiment was performed with an association time of 90 seconds and dissociation time of 300 seconds. Dilution series for MCB_D2 was 2-fold starting at 20 μM, MDB_D8 was 5-fold starting at 50 μM, GAB_D8, GAB_D23, and RBB_D10 were 5-fold starting at 20 μM. Reported measurements were analyzed using Biacore Insight Evaluation Software, sensorgrams were double-referenced and fit with a 1:1 binding kinetics fit model.

#### 3.4. Determination of GABARAP Activity by AlphaScreen assay

We used AlphaScreen assay as described in Leveille *et. al.*[31] to measure the inhibition of GABARAP-K1 interaction by the computationally-designed macrocycles. K1 is a previously described GABARAP binder with $K_D$ of 10 nM[20]. Biotin-labeled peptide K1 was used at a final concentration of 10 nM and incubated with 10 nM (final concentration) of 6xHis-GABARAP in a final reaction volume of 50 μL. Computationally-designed inhibitor peptides were serially diluted with 1:3 dilutions using the highest final concentration of 50 μM and added to the reaction mixture. The buffer used was 25 mM HEPES pH 7.3, 150 mM NaCl, 0.01% Tween, 1 mg/mL BSA, and 0.5% DMSO. The plate was covered in foil, centrifuged at 1500 rpm for 2 minutes, and incubated for 150 mins at room temperature with shaking. 20 μg/mL (final concentration) of the Streptavidin donor beads and nickel chelate acceptor beads were added in the dark and incubated for another 45 minutes. Data was collected on a Tecan plate reader using excitation at 680 nm and emission at 520-620 nm. Data were normalized to 0% (buffer only) and 100% (protein and tracer peptide, no inhibitor) controls. $IC_{50}$ values were obtained from curve fits using GraphPad Prism 9 software, using the equation $Y = \frac{100}{(1+(\frac{X}{IC_{50}})^h)}$ where X is the concentration of inhibitor and h is the Hill coefficient. At least three independent replicates were used to calculate the average $IC_{50}$ and the standard error of the mean (SEM).

#### 3.5. Crystallization of Protein-Cyclic Peptide Complexes

##### 3.5.1. MCL1 with Cyclic Peptide

MCL1 (18.5 mg/ml) and macrocycle MCB_D2 were mixed in 1:2 molar ratio and incubated for 30 min at room temperature. Upon addition of the MCB_D2 to the protein, we observed some precipitation. This precipitant was removed by centrifugation prior to crystallographic screening. Crystallization experiments for the MCL1–MCB_D2 complex were conducted using the sitting drop vapor diffusion method. Initial crystallization trials were set up in 200 nL drops using 96-well crystallization plates. Crystal drops were imaged using the UVEX crystal plate hotel system by JANSi. Diffraction quality crystals for the complex appeared in 0.2 M Sodium chloride, 0.1 M Bis-Tris pH 6.5, and 25% (w/v) Polyethylene glycol 3,350 (Index, Hampton Research) in 2 weeks.

#### 3.5.2. GABARAP and GABARAPL1 with Cyclic Peptides

Cyclic peptides GAB_D8 and GAB_D23 were dissolved in 10 mM Tris-HCl, 150 mM NaCl, pH 7.0 and were each mixed with both GABARAP and GABARAPL1, targeting a peptide:protein molar ratio of 3:2. After incubation for 10 min at room temperature any insoluble components were removed by centrifugation (10 min at 20,000 × g and 4°C). The protein-peptide complexes were concentrated using Amicon Ultra-0.5 centrifugal filter units, 3 kDa cut-off (Merck) until a final protein concentration of 6–8 mg/ml (GABARAPL1-GAB_D8) or 13–15 mg/ml (GABARAP-GAB_D23) was reached. Samples were once again cleared of particles by centrifugation (30 min at 20,000 × g and 4°C) prior to application in crystallization experiments. Search for crystallization conditions was performed by the sitting-drop vapor diffusion method using robotic systems Freedom Evo (Tecan) and Mosquito LCP (SPT Labtech) with commercially available screening sets. Experiments were set up by combining 200 nl of protein-peptide complex with 100 nl (for GABARAPL1-GAB_D8) or 200 nl (for GABARAP-GAB_D23) of reservoir solution, and plates were incubated at 20°C. Crystals appeared for a number of conditions, which were subjected to optimization as appropriate. Diffraction-quality samples used for X-ray structure determination developed with reservoir solutions containing 0.17 M ammonium sulfate, 25.5% (w/v) PEG 4,000 and 15% (v/v) glycerol for GABARAPL1-GAB_D8, and 0.1 M MES pH 5.0, 30% (w/v) PEG 6,000 in the case of GABARAP-D23. Diffraction data were collected at 100 K on beamline BM07 of the European Synchrotron Radiation Facility (ESRF; Grenoble, France) tuned to an X-ray wavelength of 0.9795 Å, using a Pilatus 6M detector (DECTRIS). Data processing was carried out with XDS and XSCALE[32] and included reflections up to a diffraction limit of 1.5 Å for GABARAP-GAB_D23 and 2.5 Å for GABARAPL1-GAB_D8. The GABARAP-GAB_D23 structure featuring space group C2 was determined by molecular replacement (MR) using MOLREP[33] with the structure of

GABARAP from its K1 peptide complex (PDB ID: 3D32[4]) as template. For the GABARAPL1-GAB_D8 complex, initial evaluation suggested tetragonal symmetry but with strong indications of twinning. Data integration in maximal *translationengleiche* subgroups followed by MR search via MoRDa[34] revealed $P2_12_12_1$ as true space group, with near-perfect pseudomerohedral twinning accounting for apparent Laue group 4/mmm. In order to avoid bias in cross-validation, this pseudosymmetry of the data was explicitly accounted for in flag assignment. The solution obtained for GABARAPL1-GAB_D8 was subjected to a round of automated rebuilding in phenix.autobuild[35]. In either case, model refinement was performed with phenix.refine[36], alternating with interactive rebuilding in COOT[37], which included stepwise introduction of cyclic peptides GAB_D8 and GAB_D23. According to validation using MolProbity[38] as well as the wwPDB validation system (https://validate-rcsb-2.wwpdb.org), both models feature good geometry. For detailed statistics of data collection and refinement refer to Table S6.

**3.5.3. RbtA with Cyclic Peptide and apo RbtA**

RbtA (10 mg/ml) and RBB_D10 were mixed in a 1:5 molar ratio and incubated for 30 min at room temperature. Initial crystallization trials were set up in 200 nl drops using 96-well crystallization plates and the experiments were conducted by sitting drop vapor diffusion method. Crystal drops were imaged using the UVEX crystal plate hotel system by JANSi. Diffraction-quality crystals for the RbtA–RBB_D10 complex appeared in 0.2 M Lithium sulfate, 0.1 M Tris pH 8.5 and 40 % (v/v) PEG 400 (JCSG Plus, Hampton Research). Additionally, we soaked the crystals in 22.32 mg/ml RBB_D10 for 5 minutes before flash freezing. Crystals for RbtA alone (18.7 mg/ml) were grown in 0.1 M Bis-Tris pH 6.5 and 20 % (v/v) PEG 5,000 MME (SG1, Molecular Dimensions). All crystals were flash-cooled in liquid nitrogen before shipping to the synchrotron for data collection.

Diffraction data were collected at the NSLS2 beamline AMX/FMX (17-ID-1/17-ID-2). X-ray intensities and data reduction were evaluated and integrated by XDS[32] and merged/scaled by Pointless/Aimless in the CCP4i2 program suite[39]. The X-ray crystal structure was determined by molecular replacement using the designed model for phasing by Phaser[40]. Next, the structure obtained from the molecular replacement was improved and refined by Phenix[36]. Model building was performed by COOT[37] in between the refinement cycles. The final model was evaluated by MolProbity[38]. Data collection and refinement statistics were reported in Supplementary Table S5.

Crystal structures have been deposited in the RCSB Protein Data Bank.

**3.6. Data Collection and Refinement Statistics**

**Table S5: MCL-1-MCB_D2, RbtA-RBB_D10 and RbtA**

| | **MCL-1-MCB_D2** | **RbtA-RBB_D10** | **RbtA** |
|---|---|---|---|
| Resolution range (Å) | 33.43 - 2.1 (2.21 - 2.1) | 34.56 - 2.61 (2.75 - 2.61) | 34.09 - 2.00 (2.05 - 2.00) |
| Space group | P $2_1$ $2_1$ 2 | P $2_1$ $2_1$ 2 | P 1 $2_1$ 1 |
| Unit cell | 52.48, 75.36, 43.37; 90, 90, 90 | 102.78, 169.84, 59.47; 90, 90, 90 | 57.16, 106.48, 71.31; 90.00, 107.04, 90.00 |
| Unique reflections | 10245 (1470) | 32591 (4662) | 55105 (4079) |
| Multiplicity | 7.8 (7.7) | 4.6 (4.8) | 7.0 (7.2) |
| Completeness (%) | 97.25 (99.93) | 99.9 (100.0) | 100.0 (100.0) |
| Mean I/sigma(I) | 11.85 (2.30) | 6.6 (1.0) | 6.2 (1.1) |
| Wilson B-factor (Å$^2$) | 41.81 | 58.62 | 26.72 |
| R-merge | 0.093 (0.860) | 0.151 (1.478) | 0.208 (1.616) |
| R-pim | 0.035 (0.324) | 0.088 (0.844) | 0.092 (0.728) |
| $CC_{1/2}$ | 0.998 (0.850) | 0.994 (0.409) | 0.994 (0.559) |
| Reflections used in refinement | 10241 (1470) | 32432 (2262) | 55054 (2928) |
| R-work | 0.2164 (0.3240) | 0.2123 (0.3198) | 0.1954 (0.2942) |
| R-free | 0.2719 (0.4208) | 0.2587 (0.3860) | 0.2344 (0.3416) |
| Number of non-hydrogen atoms | 1350 | 6901 | 7217 |
| macromolecules | 1319 | 6774 | 6715 |

| | | | |
|---|---|---|---|
| solvent | 31 | 78 | 485 |
| Protein residues | 166 | 893 | 884 |
| RMS(bonds) | 0.003 | 0.002 | 0.002 |
| RMS(angles) | 0.490 | 0.490 | 0.510 |
| Ramachandran favored (%) | 96.30 | 93.39 | 94.04 |
| Ramachandran allowed (%) | 3.70 | 6.39 | 5.96 |
| Ramachandran outliers (%) | 0.00 | 0.23 | 0.00 |
| Average B-factor (Å$^2$) | 55 | 67 | 34 |
| macromolecules | 55 | 67 | 34 |
| solvent | 49 | 55 | 35 |

**Table S6: GABARAPL1-GAB_D8 and GABARAP-GAB_D23**

| | **GABARAPL1-GAB_D8** | **GABARAP-GAB_D23** |
|---|---|---|
| **Data collection** | | |
| Resolution range (Å) | 58.42–2.52 (2.59–2.52) | 20.02–1.50 (1.54–1.50) |
| Space group | P $2_1$ $2_1$ $2_1$ | C 1 2 1 |
| Unit cell: a, b, c (Å)<br>α, β, γ (°) | 51.29, 116.81, 116.83;<br>90.0, 90.0, 90.0 | 108.63, 43.79, 67.52;<br>90.0, 128.38, 90.0 |
| Unique reflections | 24336 (1803) | 39046 (2844) |
| Multiplicity | 13.3 (13.8) | 6.85 (6.62) |
| Completeness (%) | 99.4 (99.8) | 97.0 (95.5) |
| Mean I/sigma(I) | 10.01 (0.73) | 13.34 (0.58) |

| | | |
|---|---|---|
| Wilson B-factor (Å$^2$) | 58.53 | 32.70 |
| R-merge | 0.229 (3.146) | 0.062 (3.089) |
| R-pim | 0.065 (0.871) | 0.026 (1.280) |
| $CC_{1/2}$ | 0.996 (0.291) | 0.999 (0.346) |
| **Refinement** | | |
| Resolution range (Å) | 58.42–2.52 (2.63–2.52) | 18.42–1.50 (1.54–1.50) |
| No. reflections | 24328 (2996) | 38949 (2685) |
| R-work | 0.2147 (0.3228) | 0.1878 (0.4255) |
| R-free | 0.2586 (0.3257) | 0.2220 (0.4350) |
| Number of non-hydrogen atoms | 4542 | 2714 |
| polypeptides | 4498 | 2519 |
| ligands | 22 | 35 |
| solvent | 22 | 160 |
| Polypeptide residues | 536 | 264 |
| RMSD bonds (Å) | 0.002 | 0.006 |
| RMSD angles (°) | 0.410 | 0.836 |
| Ramachandran favored (%) | 95.95 | 96.88 |
| Ramachandran allowed (%) | 3.85 | 3.12 |

| | | |
|---|---|---|
| Ramachandran outliers (%) | 0.19 | 0.00 |
| Avg. B-factor (Å$^2$) | 58 | 38 |
| polypeptides | 58 | 38 |
| ligand | 68 | 53 |
| solvent | 50 | 44 |

Values for the highest-resolution shell are shown in parentheses.

**4. Peptide Analytical Characterization Data**

**Table S7: Purity of the chemically synthesized MCB designs**

| Design | Sequence | Purity (%) |
|---|---|---|
| MCB_D1 | GAPEILKKMADMVGYE | 87.12 |
| MCB_D2 | PPEIAWLADAVGLKDA | 97.30 |
| MCB_D3 | GLETDDPVVKPLADAV | 96.50 |
| MCB_D4 | APPEIKALADAVGLED | 96.10 |
| MCB_D7 | SSTVAFLQEAVGMPVT | 99.10 |
| MCB_D8 | EAVGLESDSELIKKLA | 91.23 |
| MCB_D12 | SVDDPLIRKLAEAVGL | 92.65 |
| MCB_D13 | SIDDPLLKKLAEAVGL | 85.24 |
| MCB_D15 | ADALKRLTVPKNLEEI | 97.10 |
| MCB_D18 | KPQPDIDPAFAEIVGF | 96.00 |
| MCB_D20 | EDTLEGIARGLLTGKV | 96.87 |
| MCB_D21 | YDTEEGIAEGLLTGKV | 92.38 |
| MCB_D22 | ADTEEGIAKGLLTGKV | 96.84 |
| MCB_D26 | GSPEIRWLMDAFGVDE | 96.32 |

**Table S8: Purity of the chemically synthesized MDB designs**

| Design | Sequence | Purity (%) |
|---|---|---|
| MDB_D1 | KKYNWMVDELVSMVGKPE | 95.04 |
| MDB_D2 | EESEDEDFRVFERVLGI | 97.00 |
| MDB_D3 | RELMEMVGEKYDQNFIL | 88.06 |
| MDB_D4 | IESDDDFKVLTDVMGLD | 95.14 |
| MDB_D6 | EDYQVLHDVLGLPLEFDD | 91.19 |
| MDB_D8 | SRKAKNKFEELWNELIDP | 95.09 |
| MDB_D10 | PGTPFAKEWSKMAGGAPL | 90.64 |
| MDB_D11 | GLDLETDSEFAKEWQKMV | 96.51 |

**Table S9: Purity of the chemically synthesized GAB designs**

| Design | Sequence | Purity (%) |
|---|---|---|
| GAB_D1 | ETGEVENIDGVEIIYP | 99.00 |
| GAB_D8 | GSEYEEDGWTVLEPD | 97.15 |
| GAB_D23 | LEDGWVDIETGKE | 99.46 |
| GAB_D26 | LDTGEVYKAPNGQEVIE | 99.00 |
| GAB_D27 | IDIDTEEEVMPGV | 95.87 |
| GAB_D28 | VMPGIIDIDTEEE | 92.15 |

**Table S10: Purity of the chemically synthesized RBB designs**

| Design | Sequence | Purity (%) |
|---|---|---|
| RBB_D1 | SSLPEYVRDVLGV | 93.77 |
| RBB_D2 | GKPGDSPWEKIESVVFG | 92.43 |
| RBB_D3 | PDPAIPENVQKVFKEHML | 92.43 |
| RBB_D4 | PDPAFPENVQKVFKDKIE | 98.04 |
| RBB_D5 | PDPAFPESIHKVFKDKII | 93.22 |
| RBB_D6 | FSKYEPVDDSYPEGIRKV | 92.11 |
| RBB_D7 | FEKYDPMDESYPENIRKV | 94.63 |
| RBB_D8 | VEGNDKGSDIYEFIRDK | 90.11 |
| RBB_D10 | KLFGPDPYLPENVQ | 97.44 |
| RBB_D14 | SVAKEIAEWIGIPSKVPP | 98.08 |
| RBB_D15 | SLAKEIADWIGIPSSVPP | 94.52 |

**MCB peptides Mass Spec and HPLC**

Analytical UPLC and mass spectrum of MCB_D1

Sequence: cyclo[APEILKKMADMVGYEG]

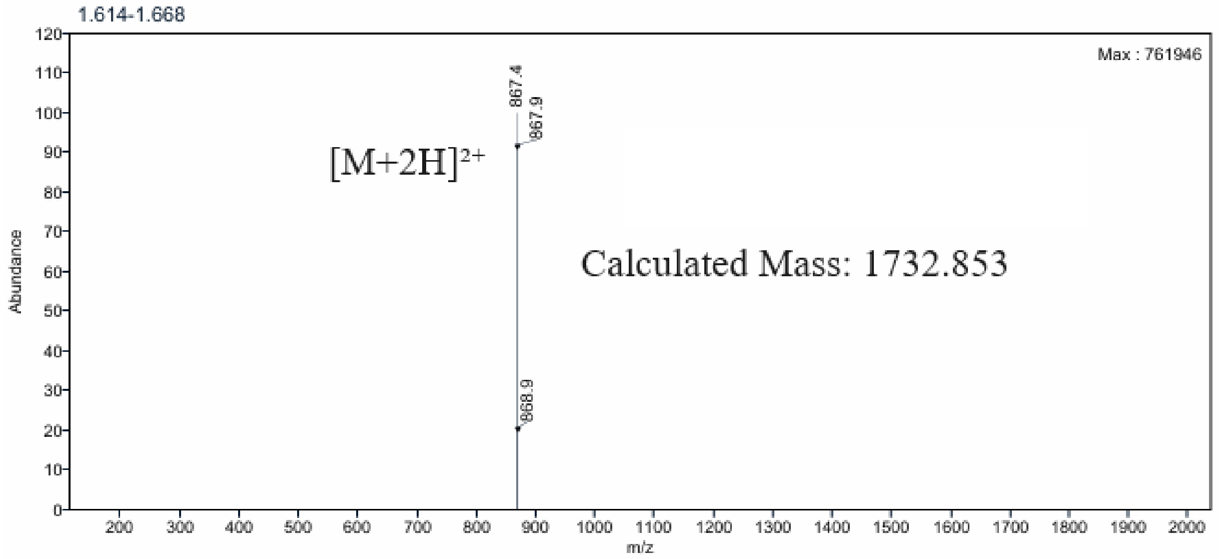

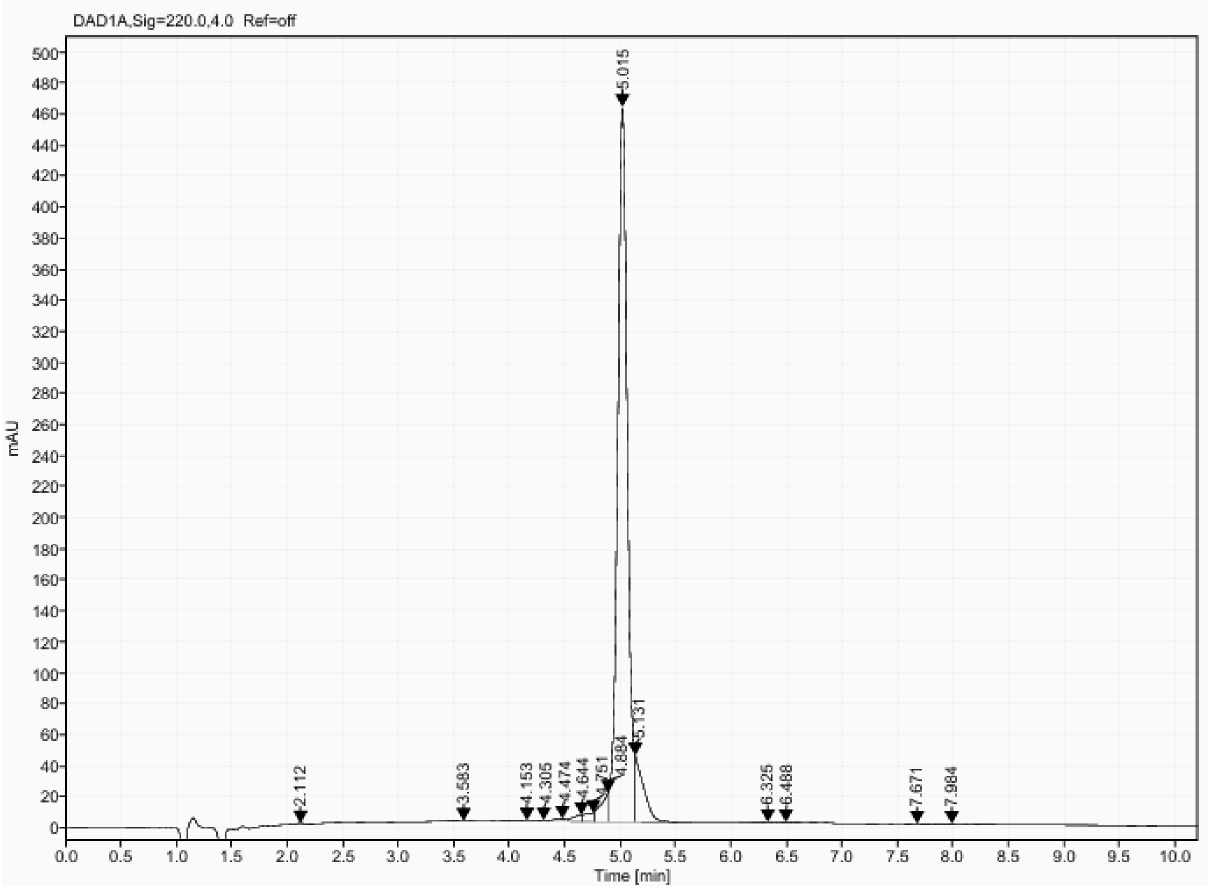

Analytical UPLC and mass spectrum of MCB_D2
Sequence: cyclo[LKDAPPEIAWLADAVG]

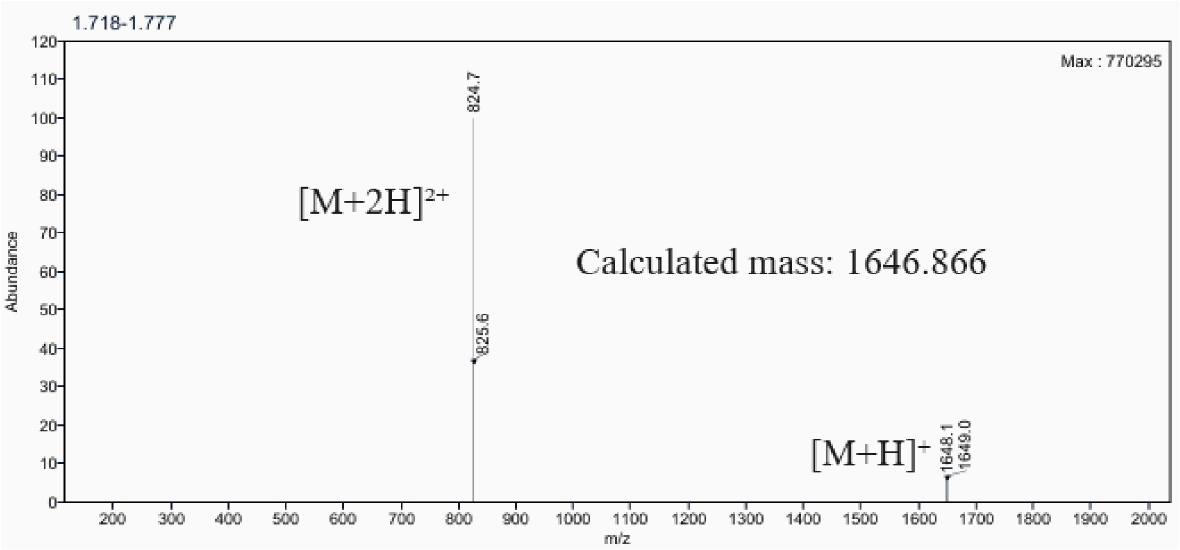

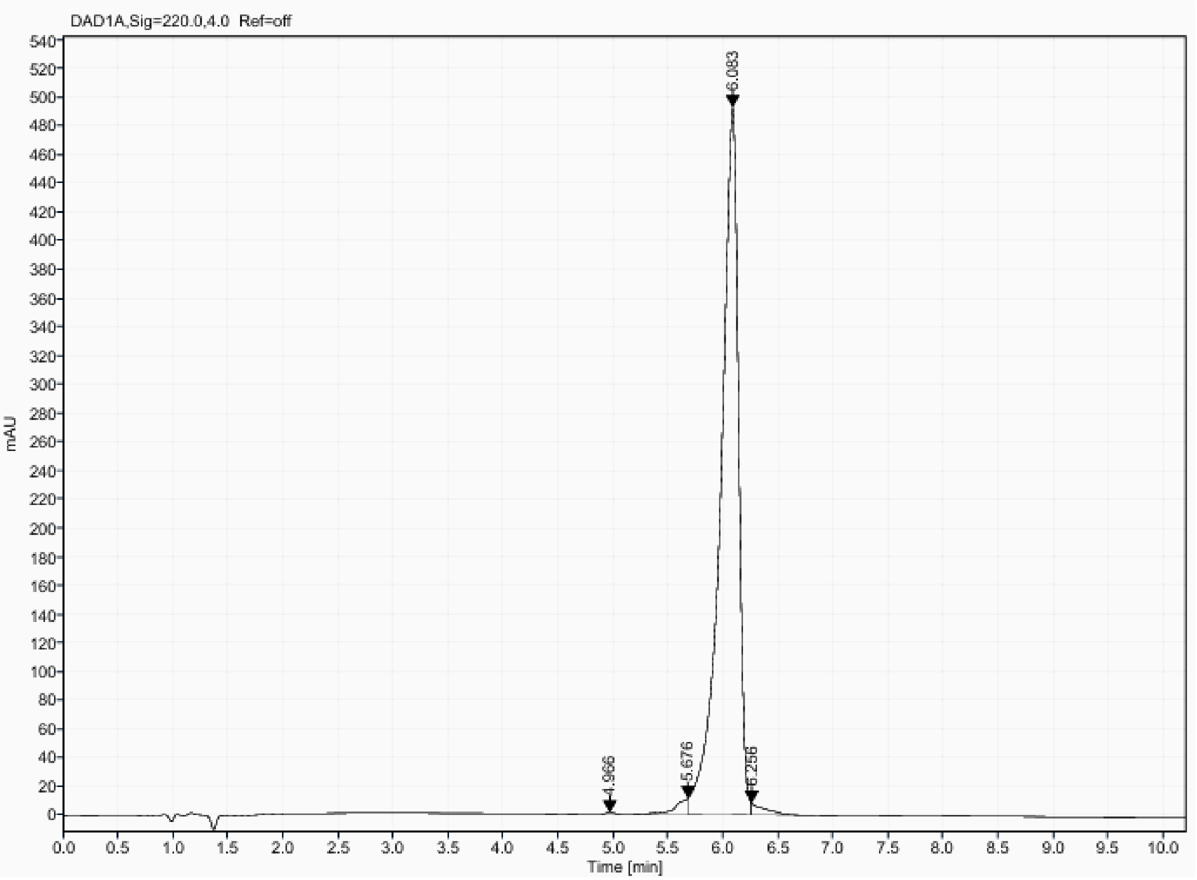

Analytical UPLC and mass spectrum of MCB_D3
Sequence: cyclo[LETDDPVVKPLADAVG]

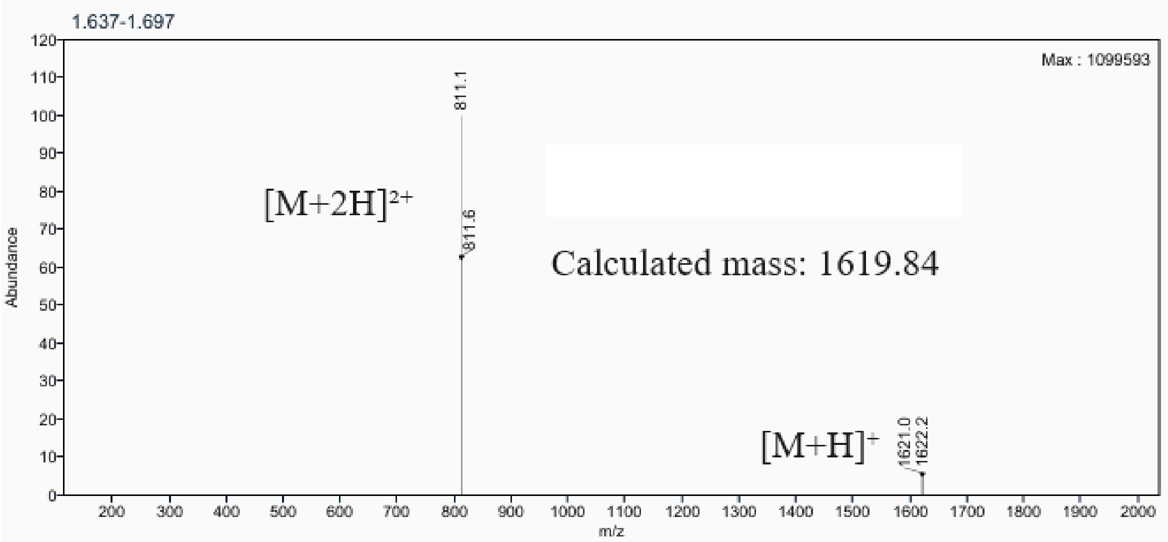

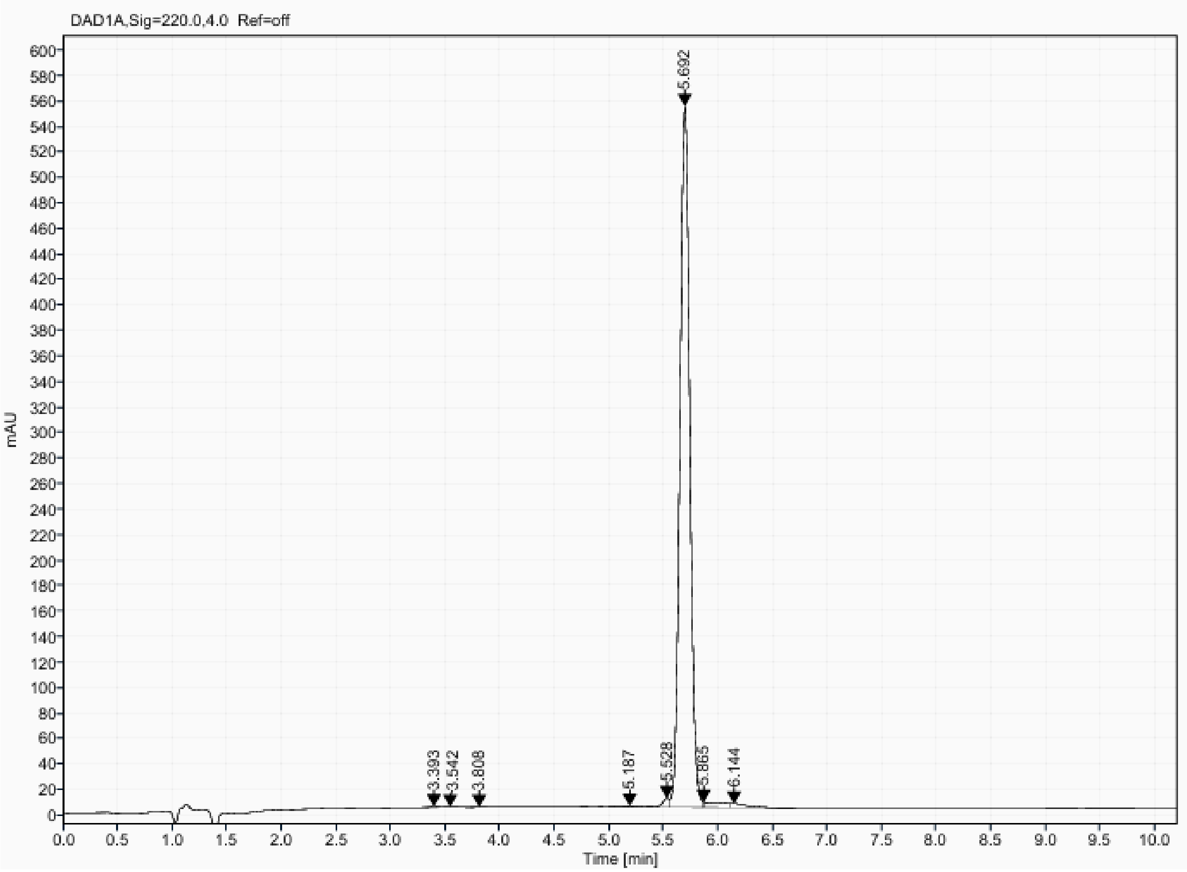

Analytical UPLC and mass spectrum of MCB_D4
Sequence: cyclo[LEDAPPEIKALADAVG]

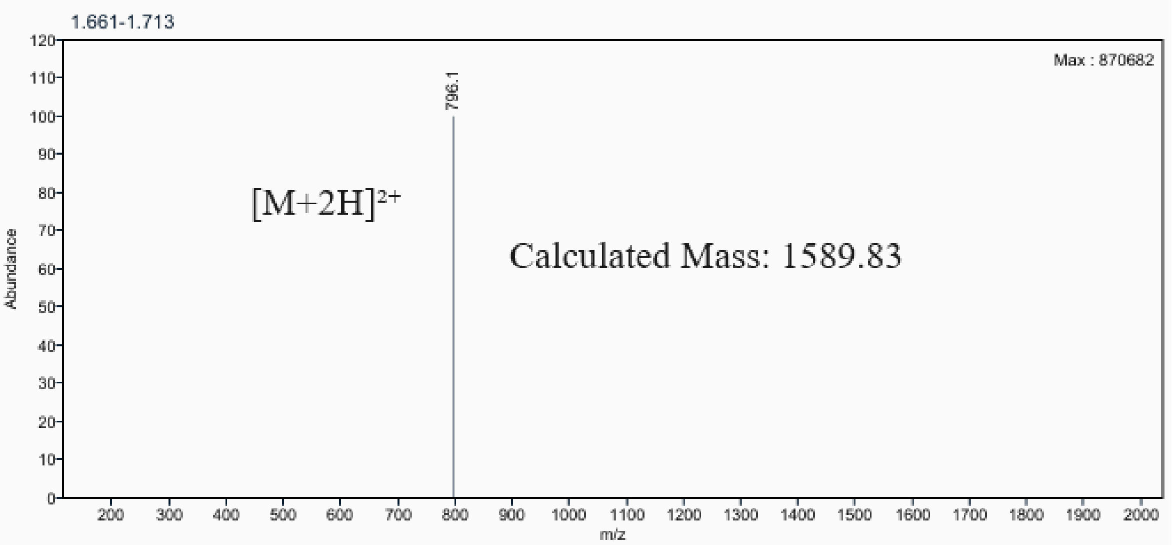

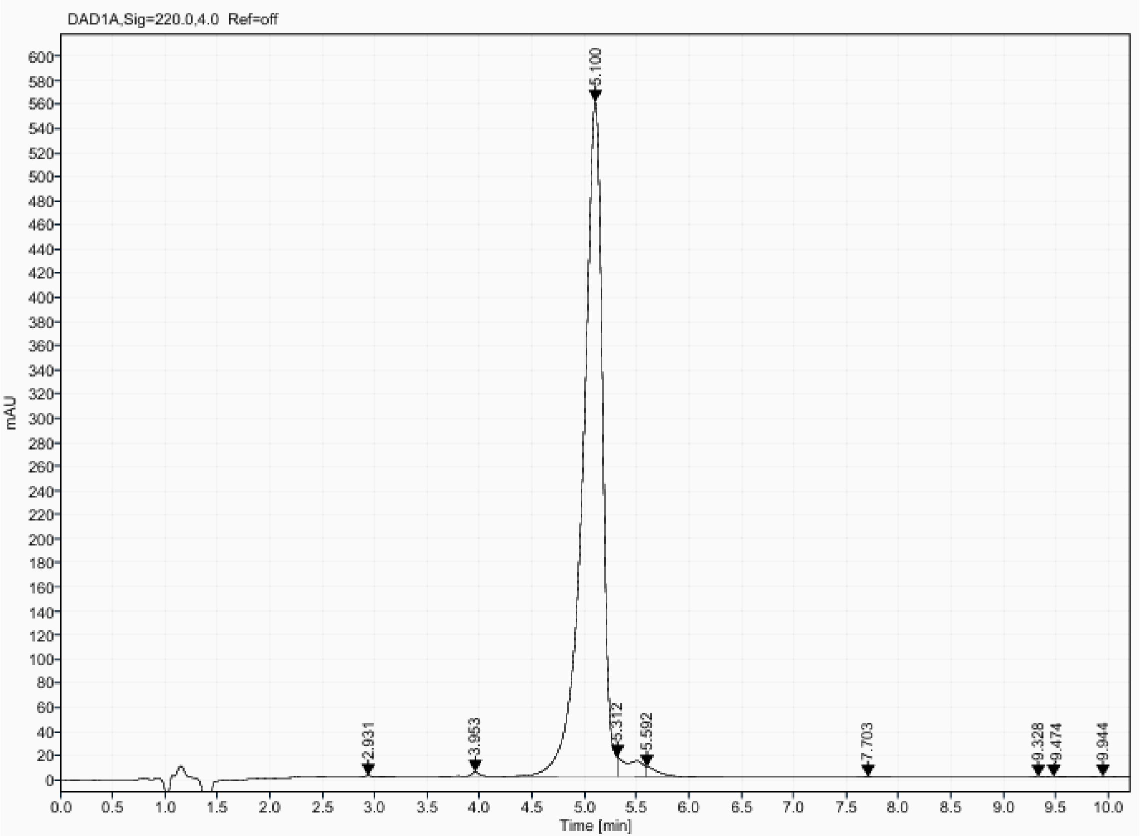

Analytical UPLC and mass spectrum of MCB_D7
Sequence: cyclo[MPVTSSTVAFLQEAVG]

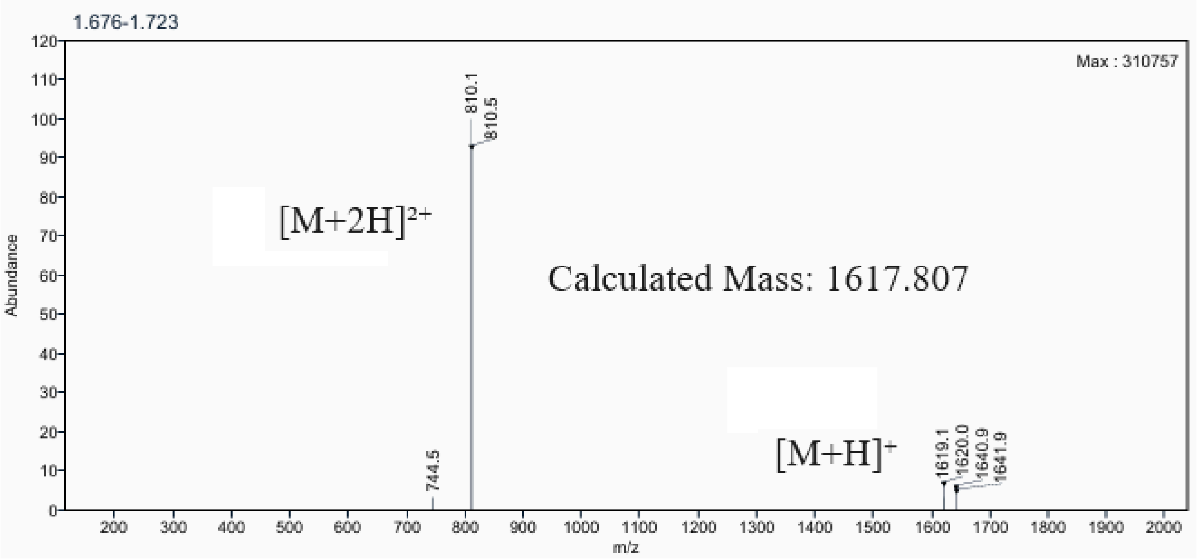

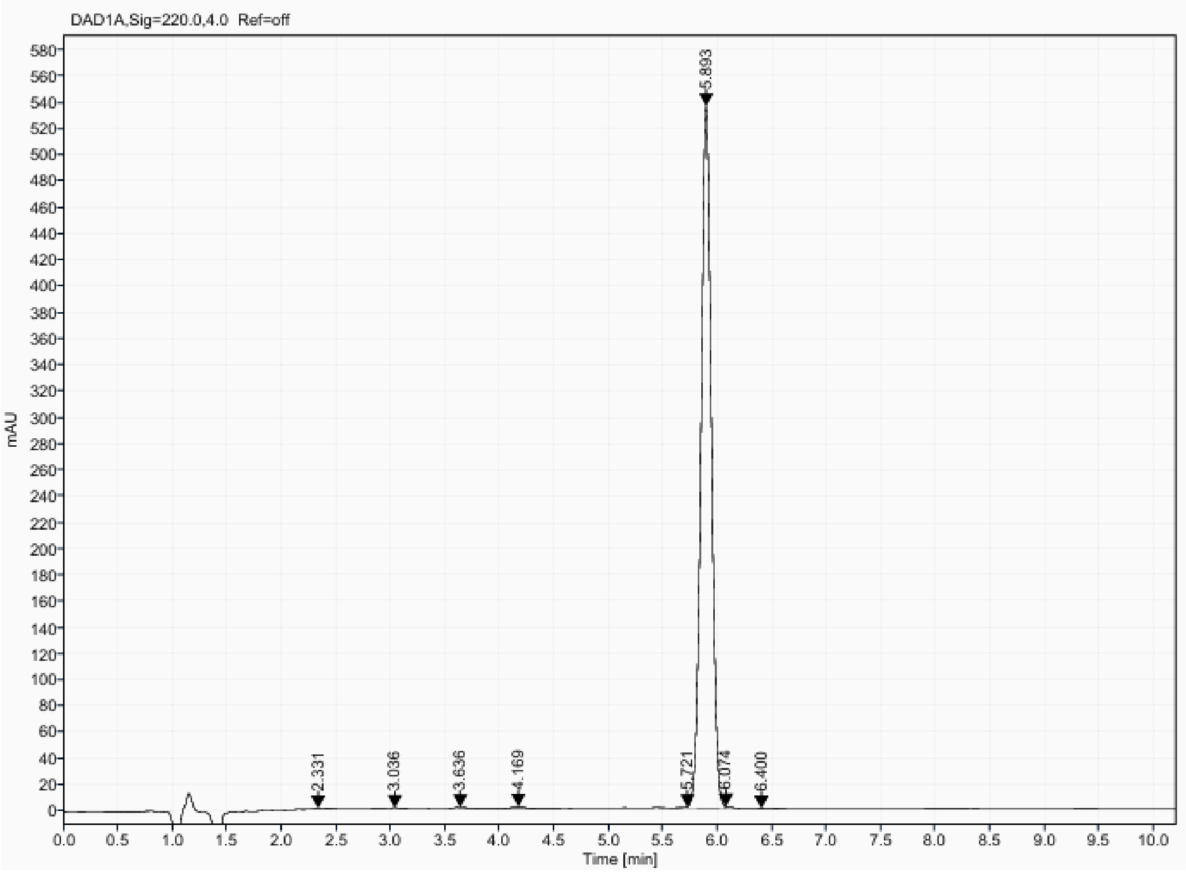

Analytical UPLC and mass spectrum of MCB_D8
Sequence: cyclo[LESDSELIKKLAEAVG]

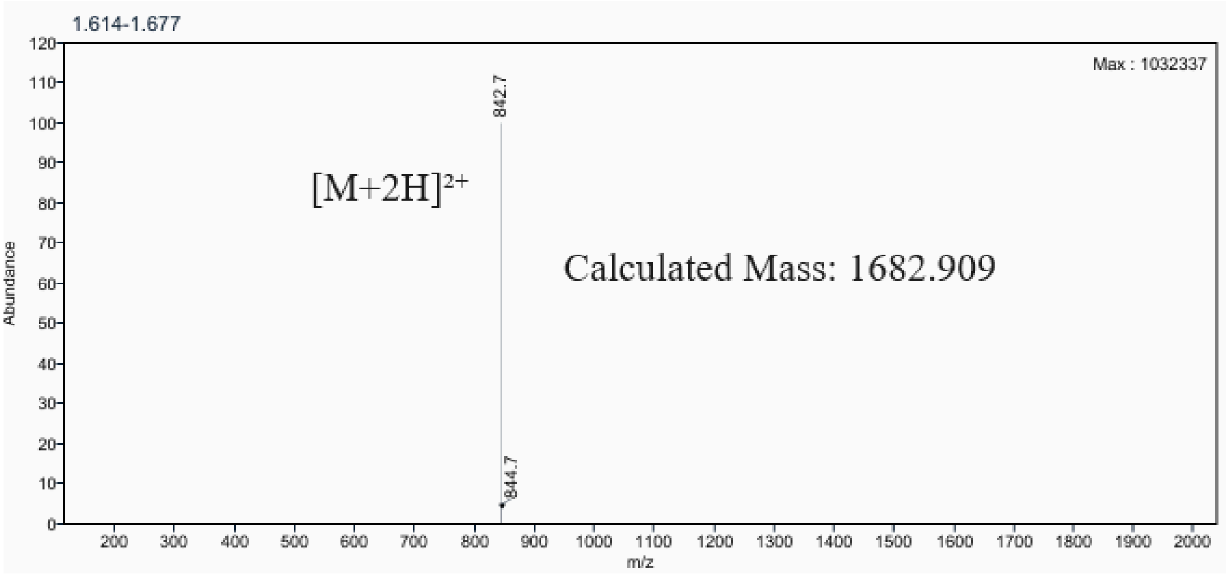

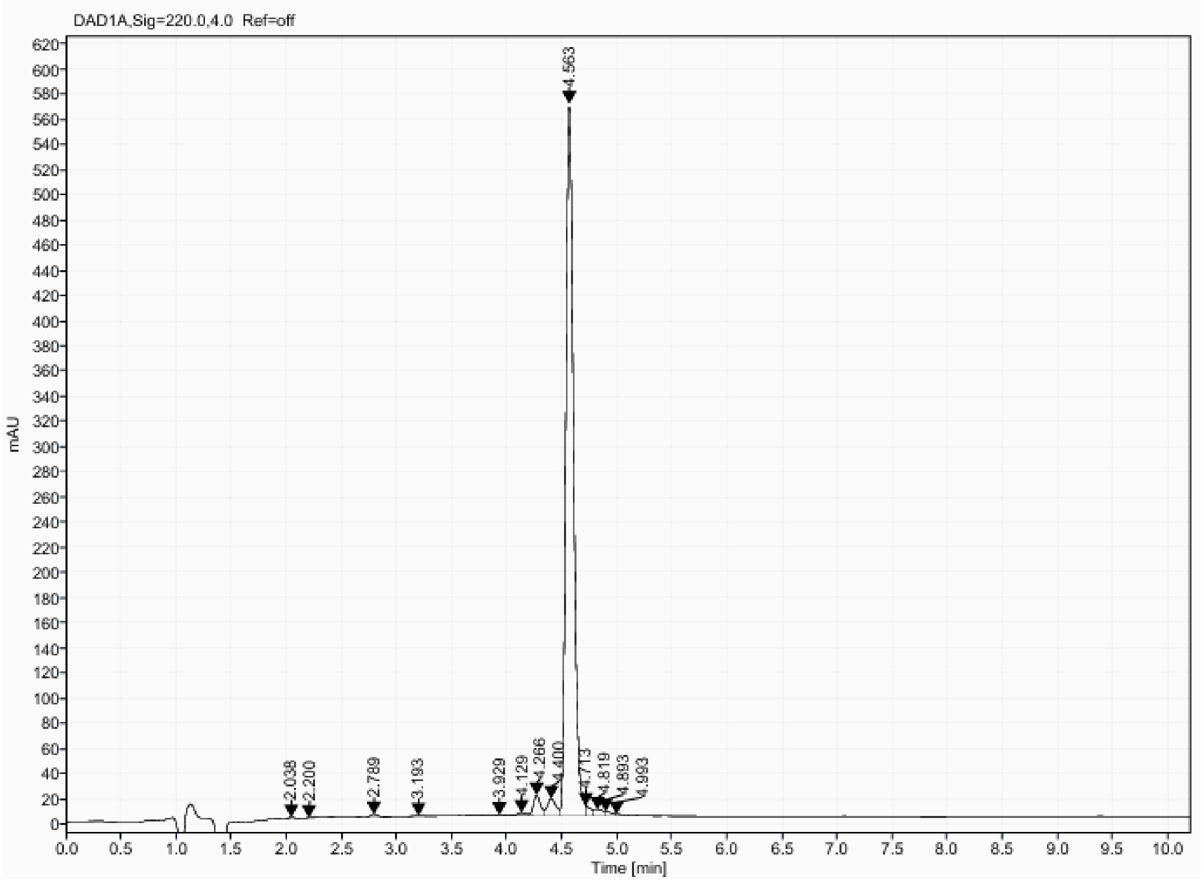

Analytical UPLC and mass spectrum of MCB_D12
Sequence: cyclo[LSVDDPLIRKLAEAVG]

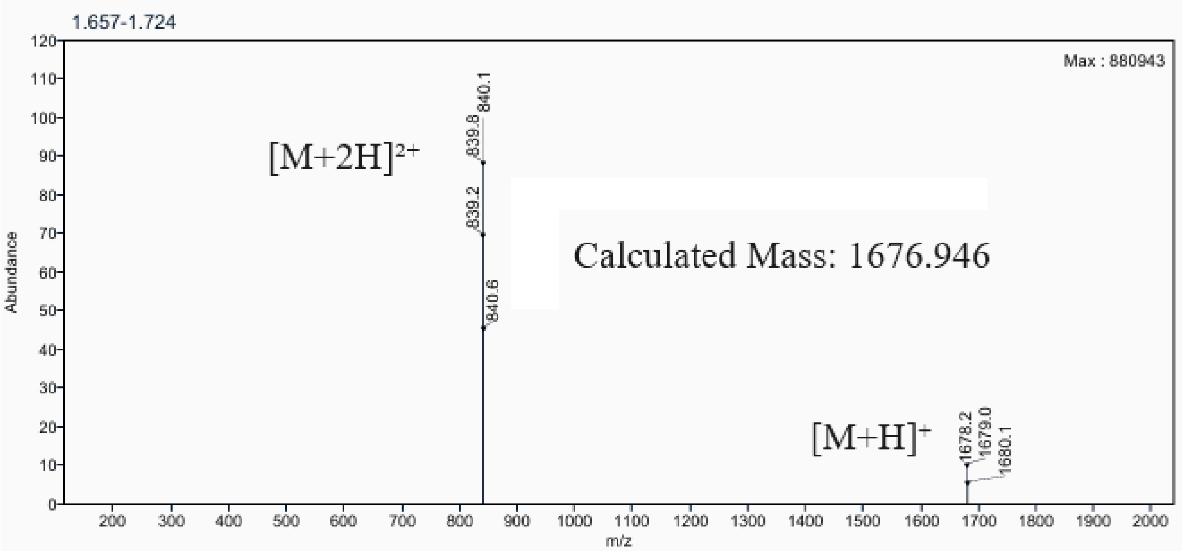

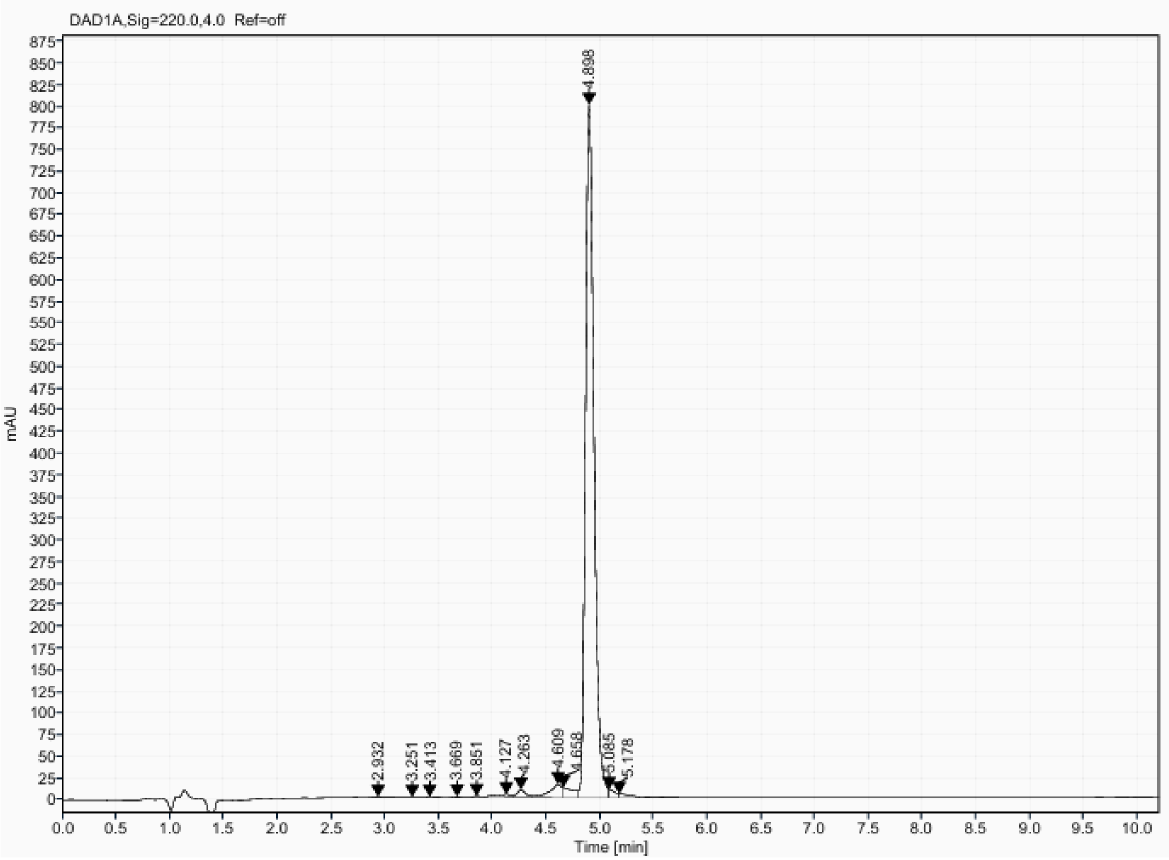

Analytical UPLC and mass spectrum of MCB_D13
Sequence: cyclo[LSIDDPLLKKLAEAVG]

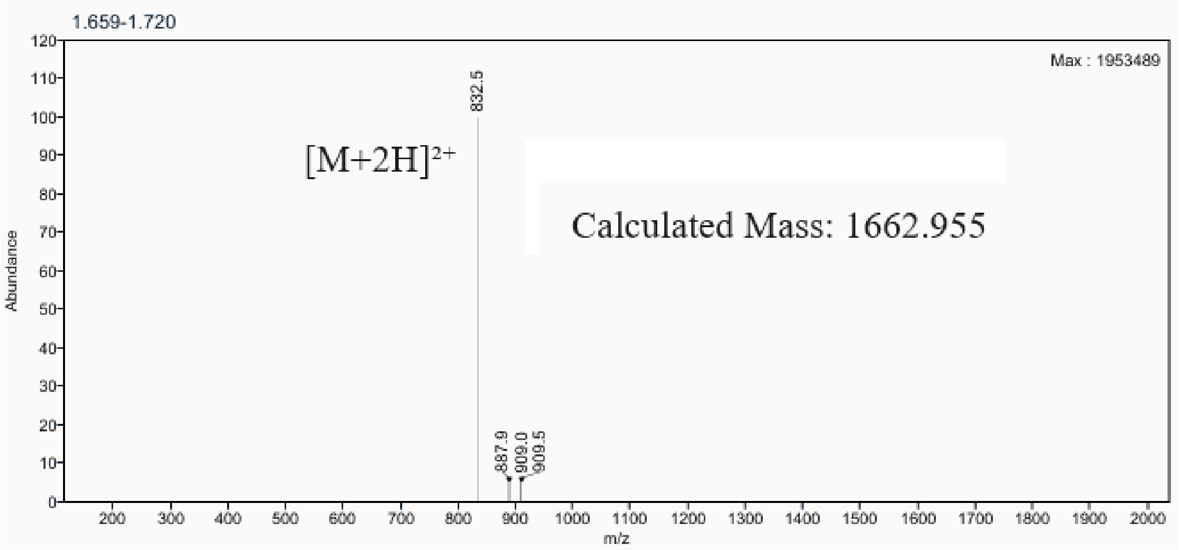

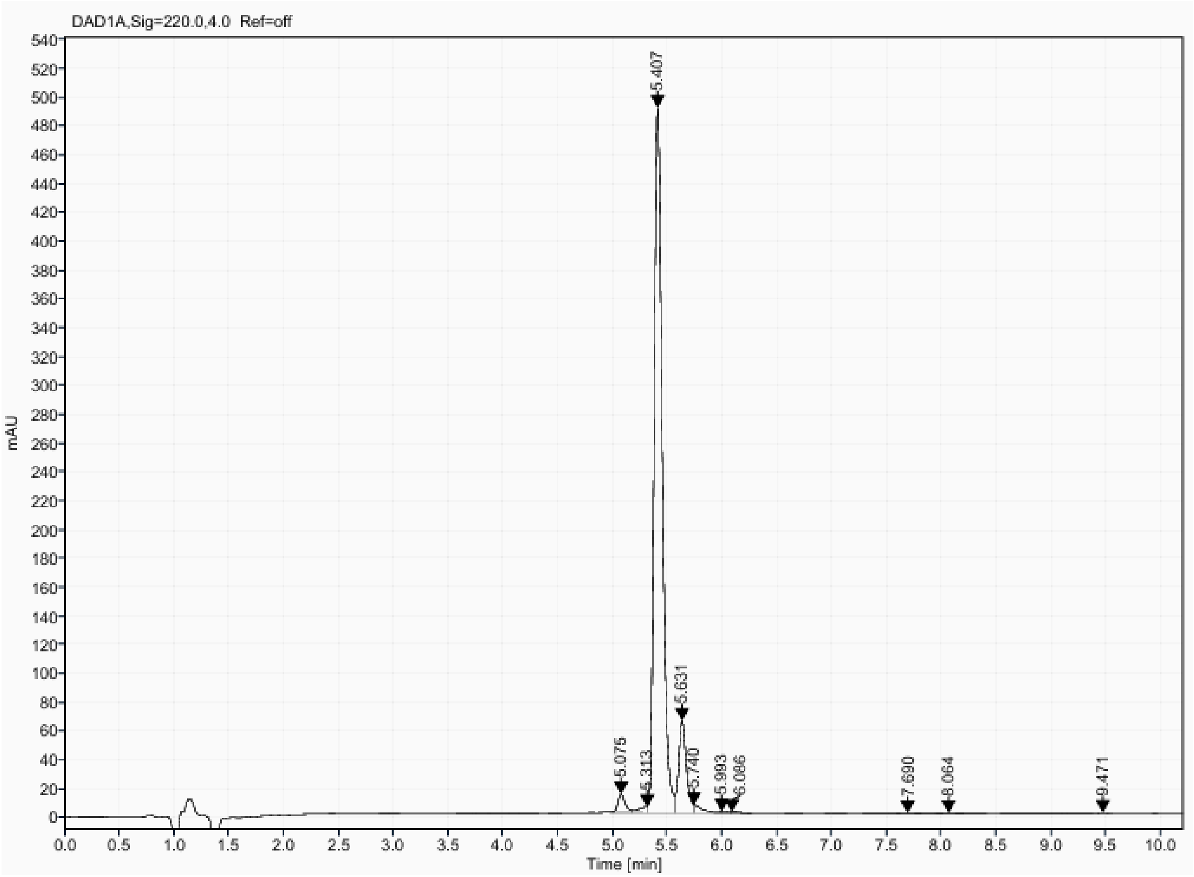

Analytical UPLC and mass spectrum of MCB_D15
Sequence: cyclo[LKRLTVPKNLEEIADA]

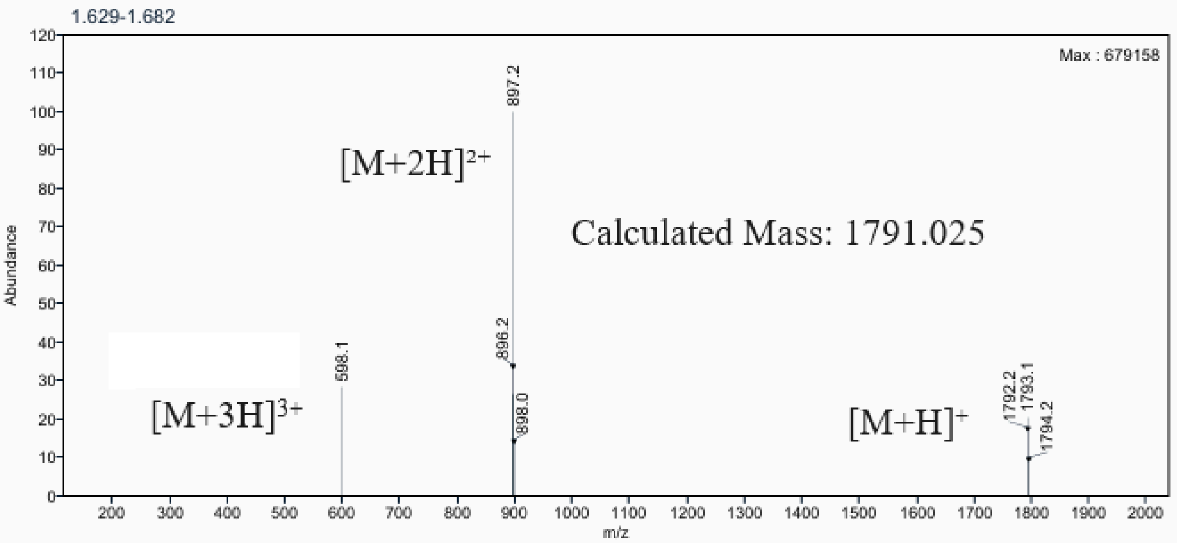

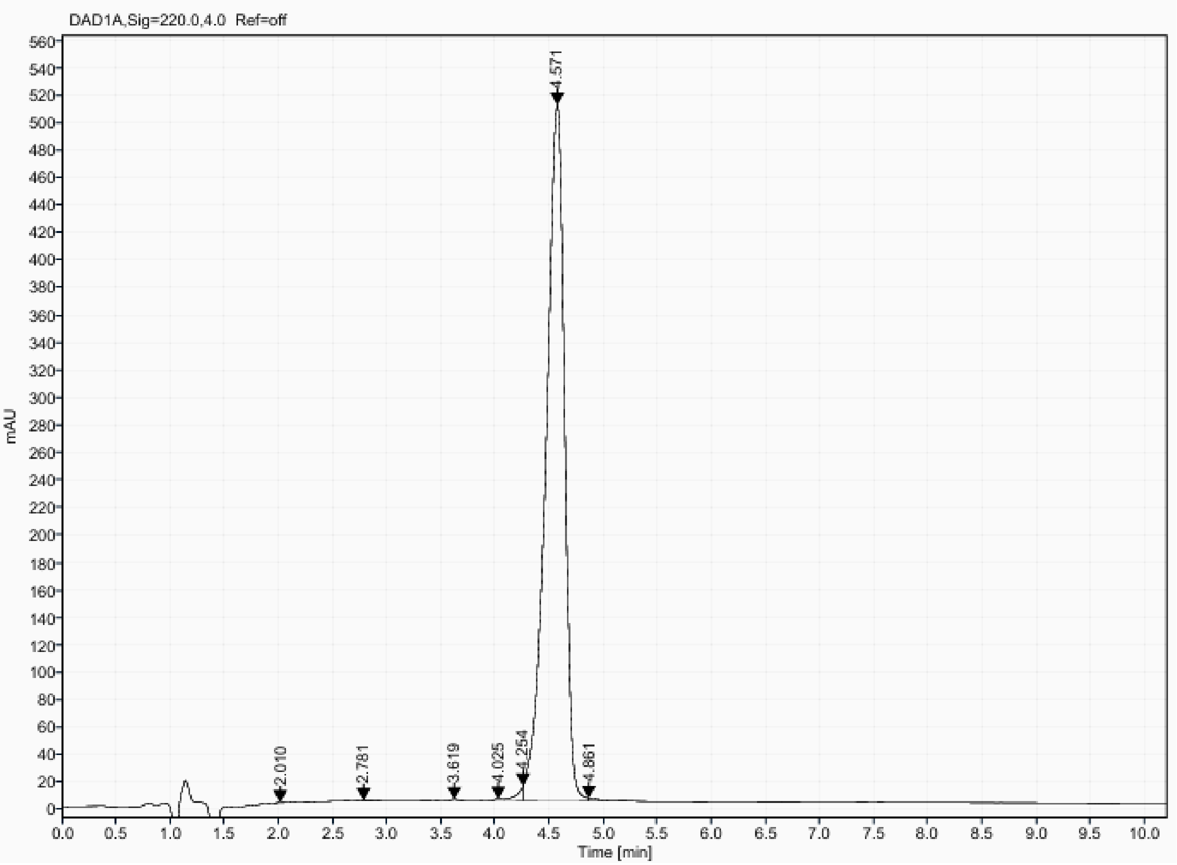

Analytical UPLC and mass spectrum of MCB_D18
Sequence: cyclo[FKPQPDIDPAFAEIVG]

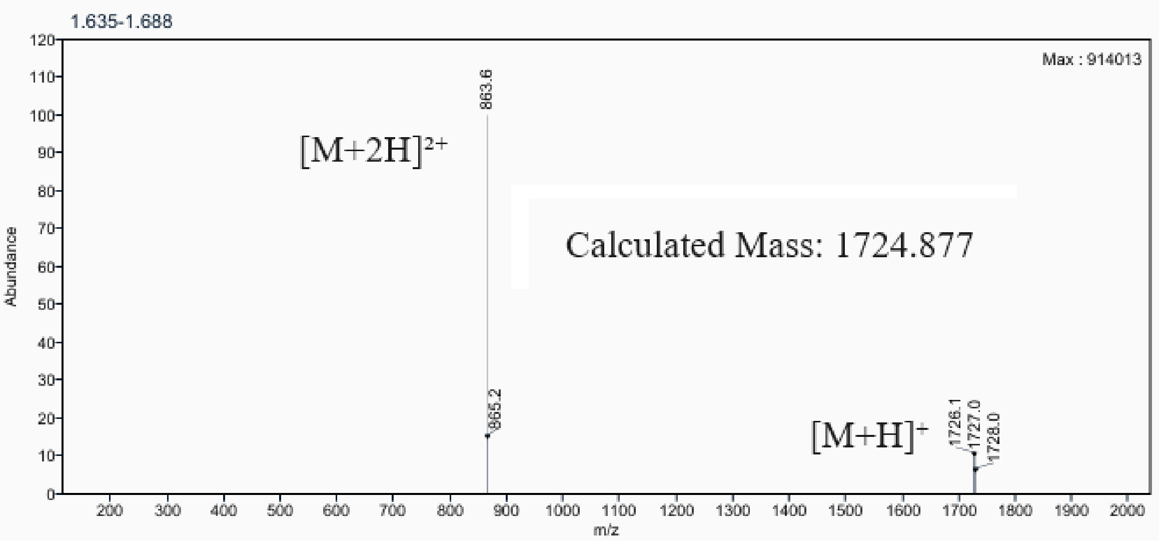

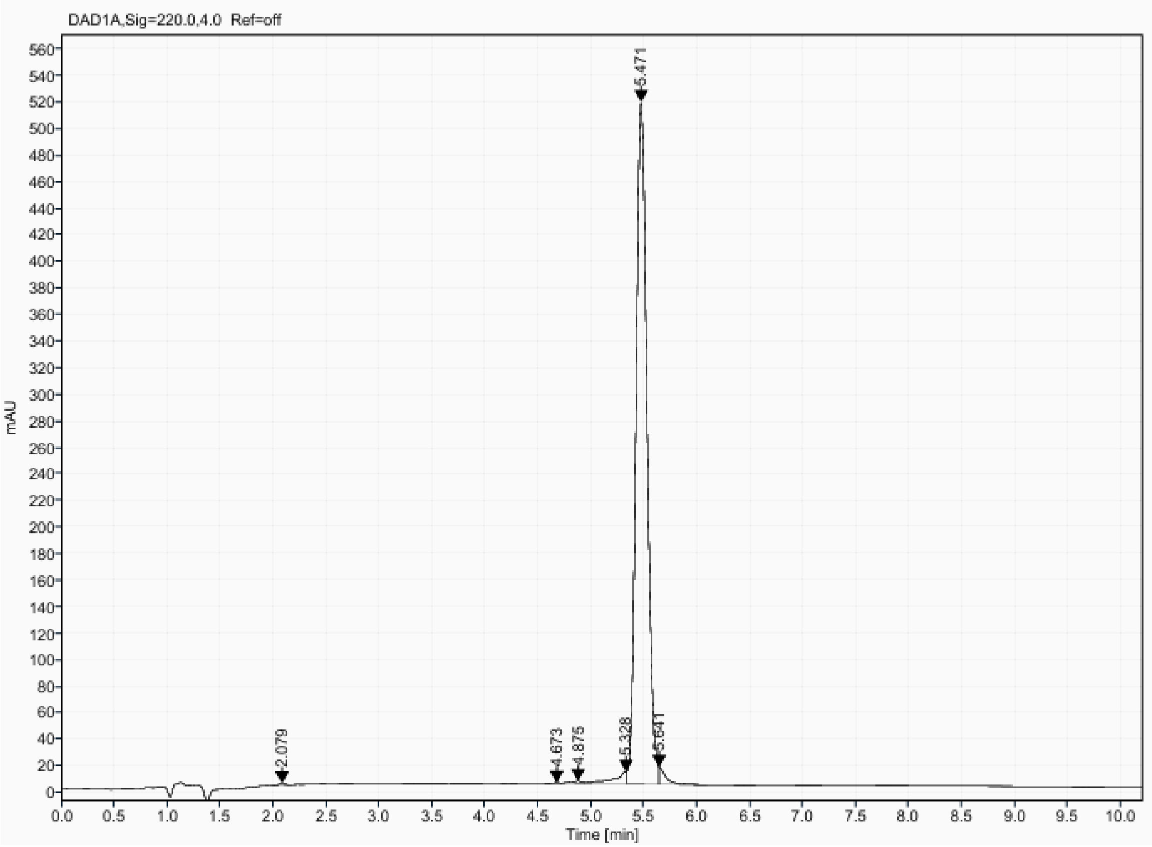

Analytical UPLC and mass spectrum of MCB_D20
Sequence: cyclo[EDTLEGIARGLLTGKV]

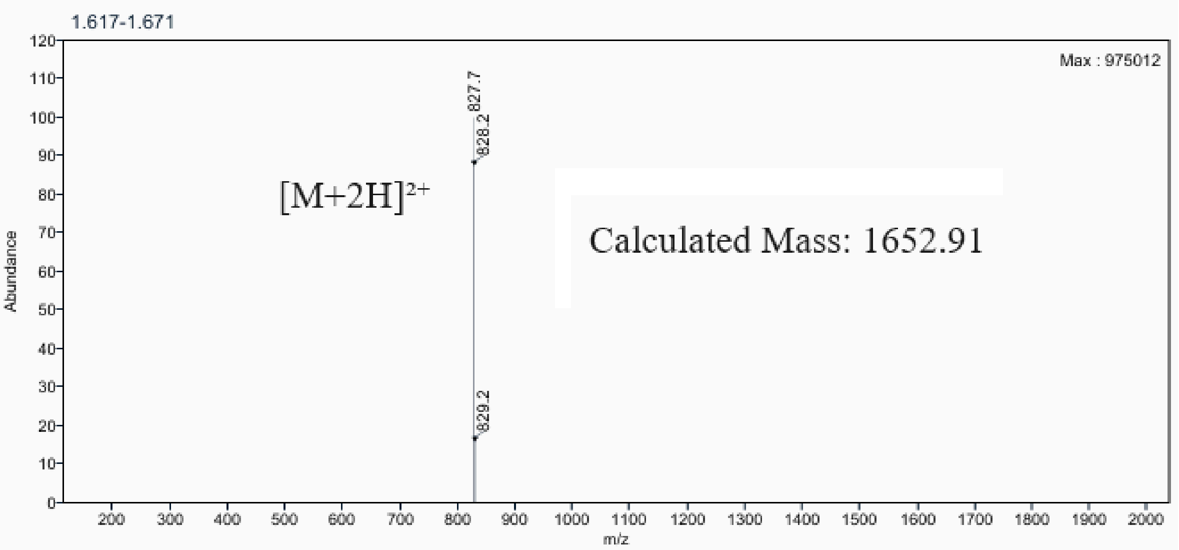

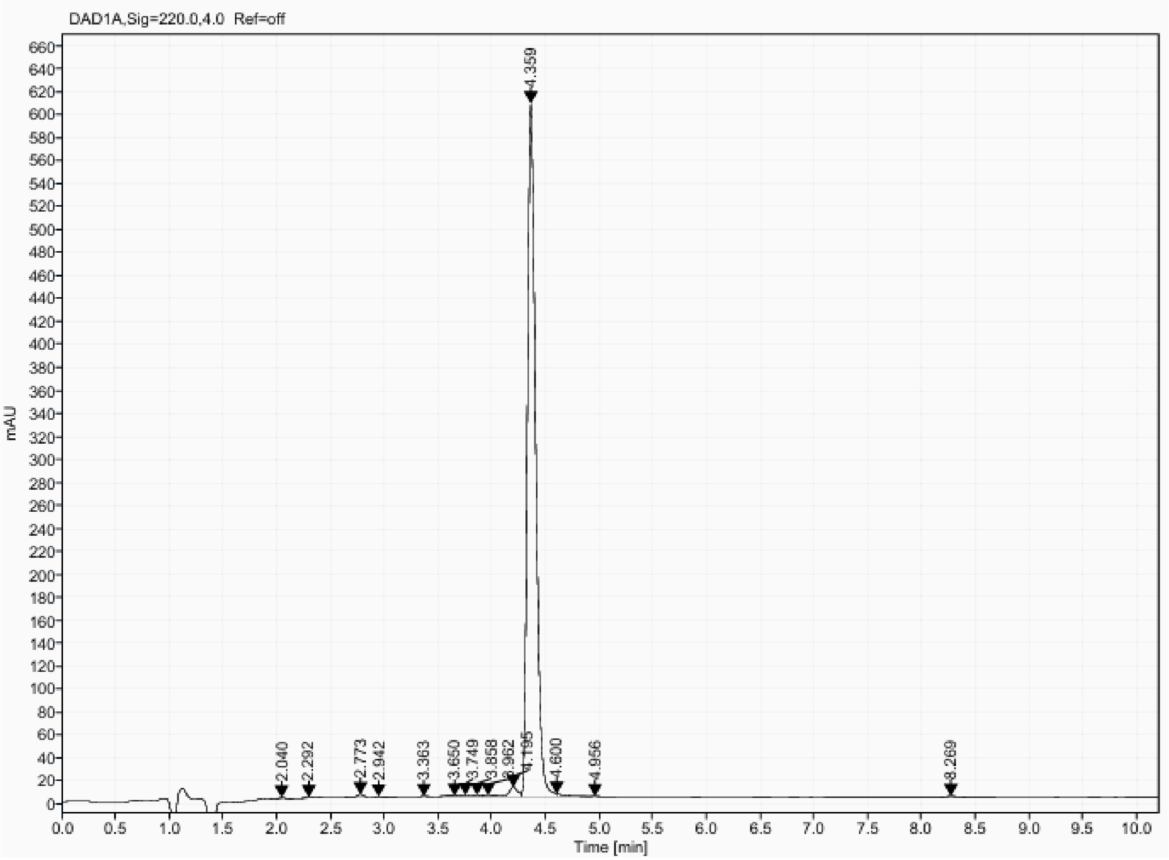

Analytical UPLC and mass spectrum of MCB_D21
Sequence: cyclo[YDTEEGIAEGLLTGKV]

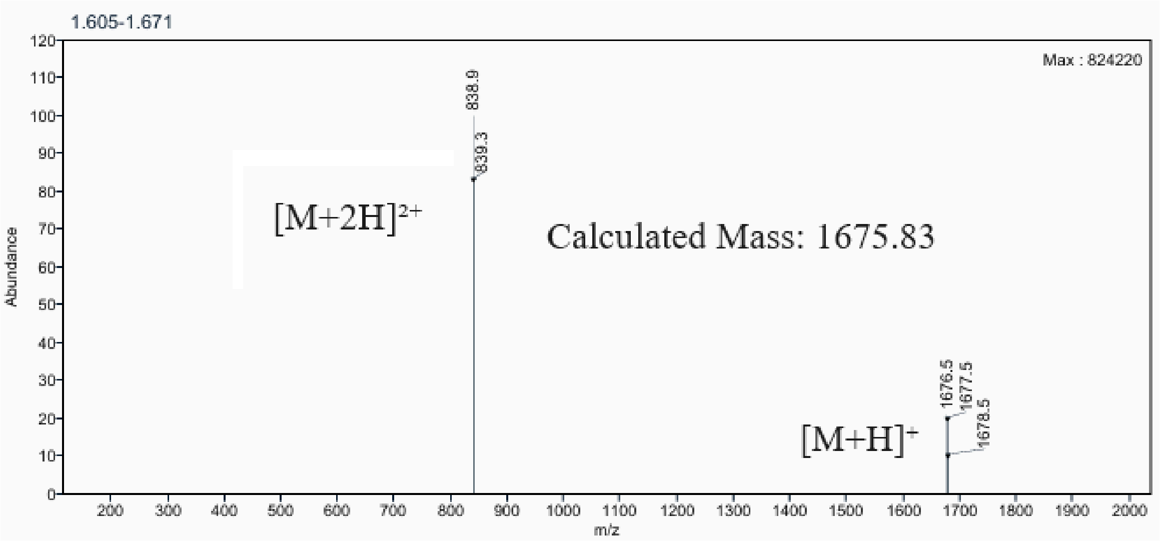

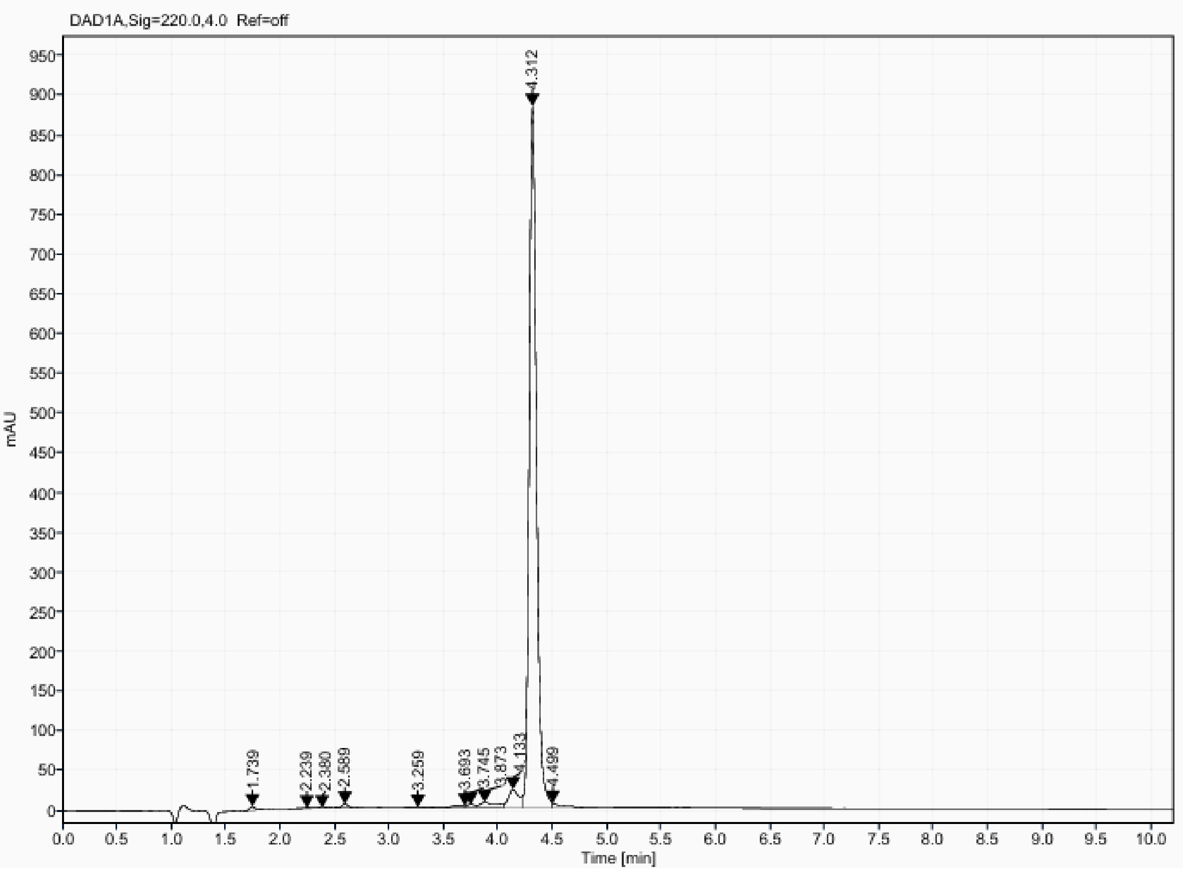

Analytical UPLC and mass spectrum of MCB_D22
Sequence: cyclo[ADTEEGIAKGLLTGKV]

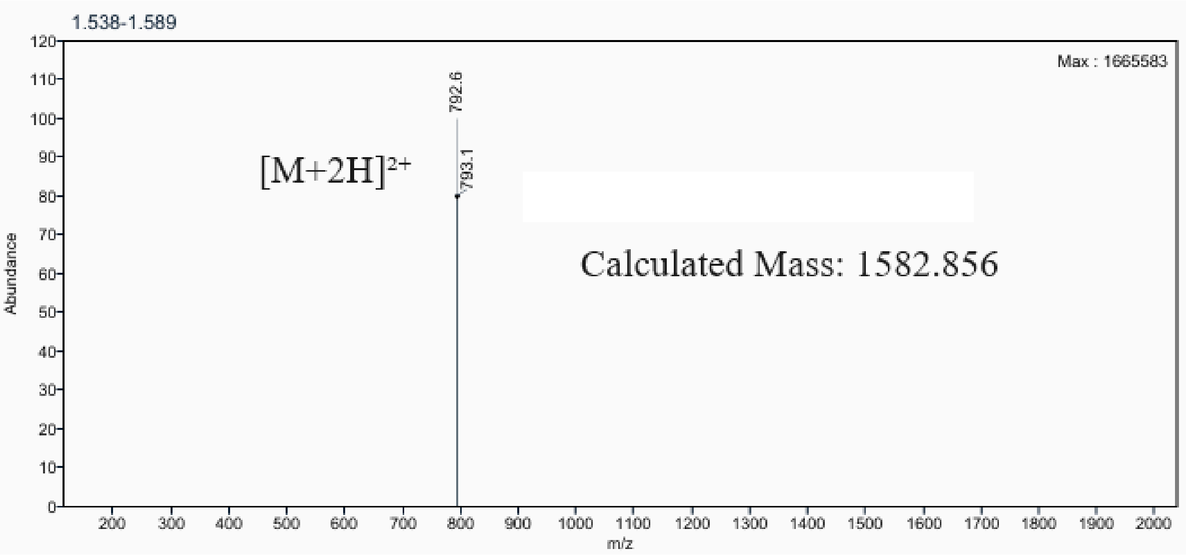

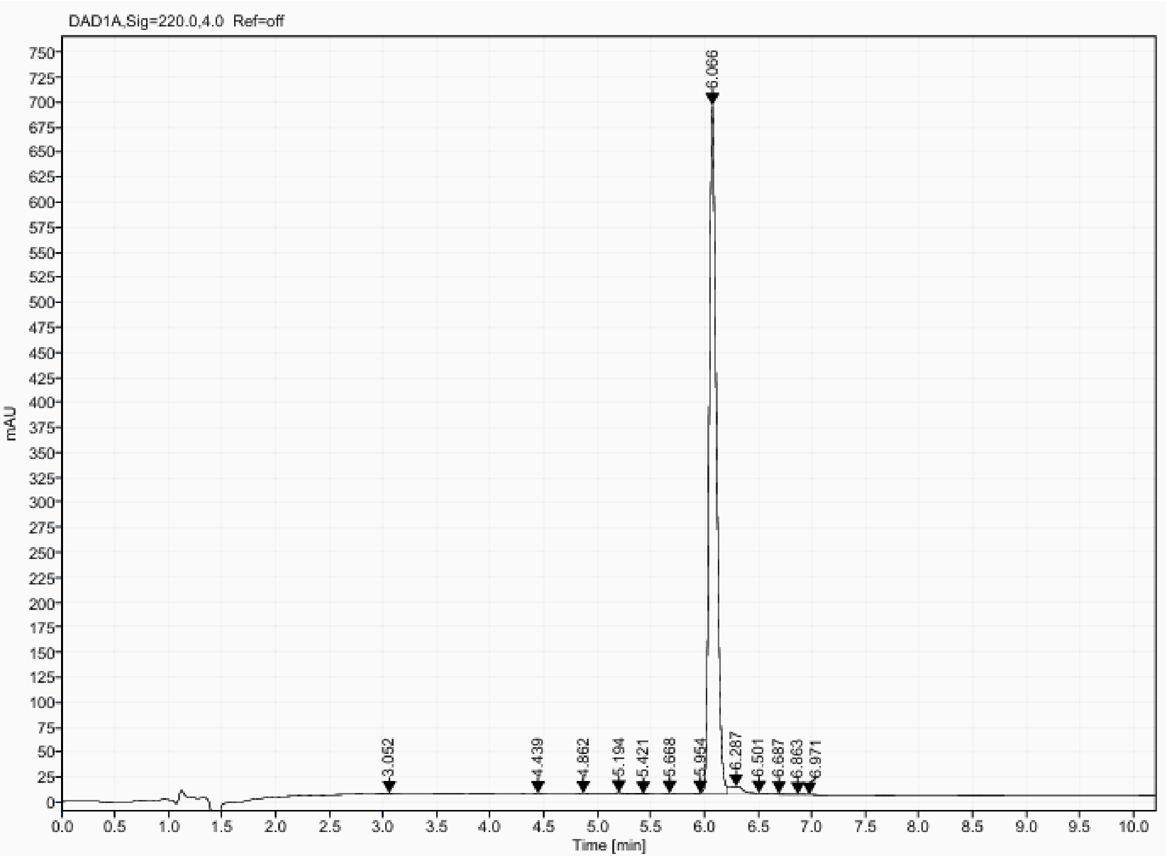

Analytical UPLC and mass spectrum of MCB_D26
Sequence: cyclo[GSPEIRWLMDAFGVDE]

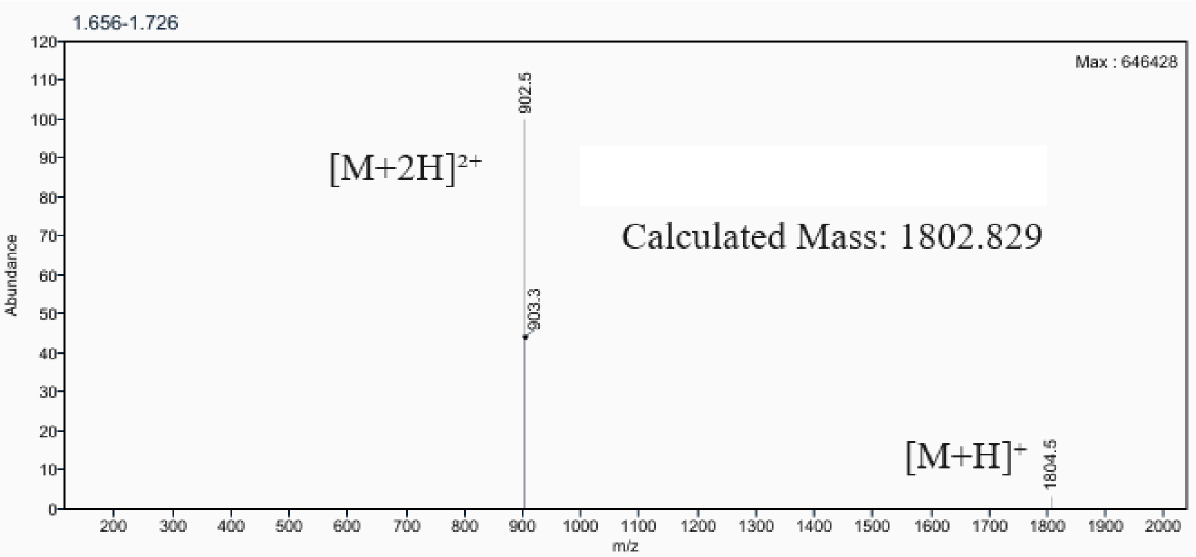

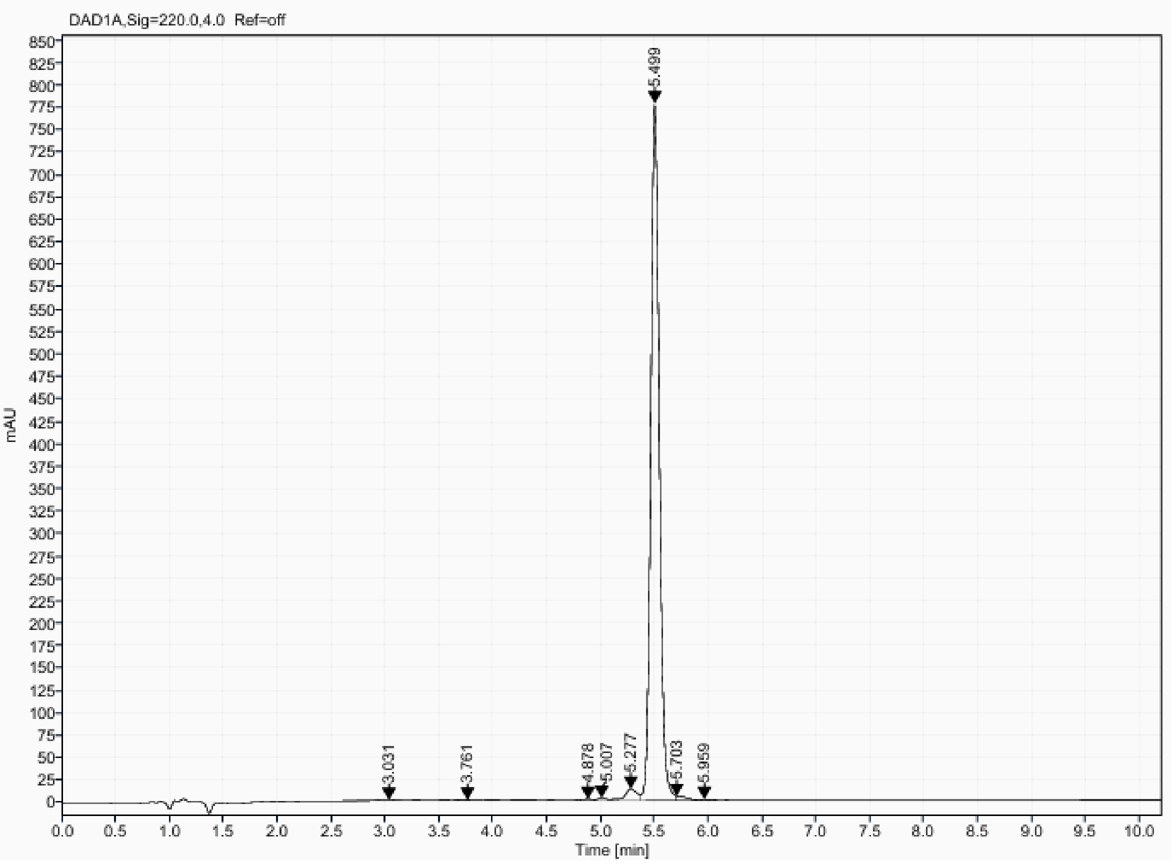

**MDB peptides Mass Spec and HPLC**

Analytical UPLC and mass spectrum of MDB_D1

Sequence: cyclo[LVSMVGKPEKKYNWMVDE]

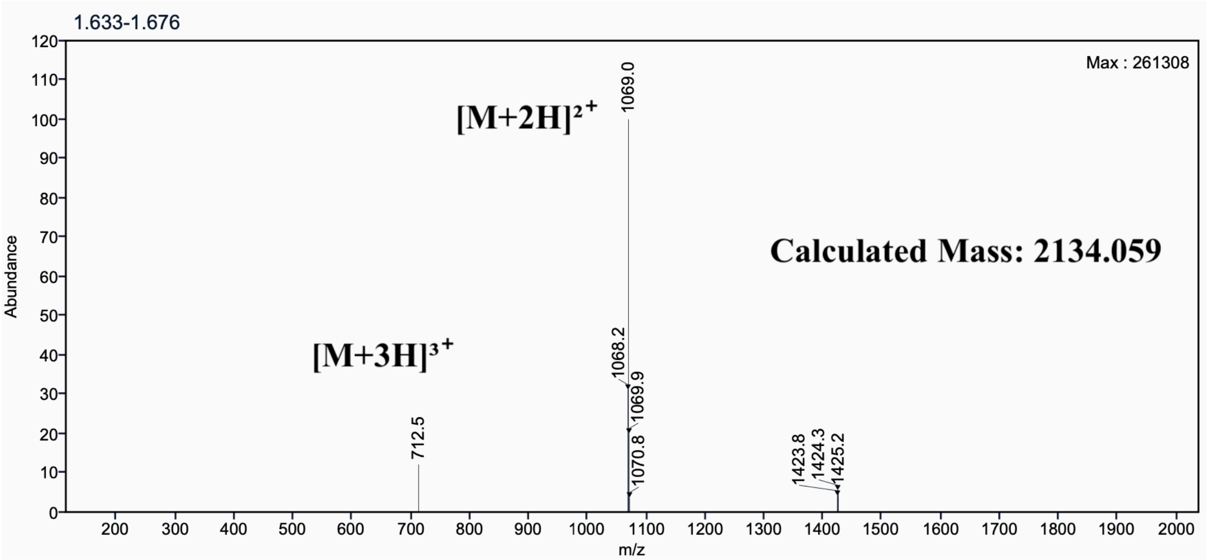

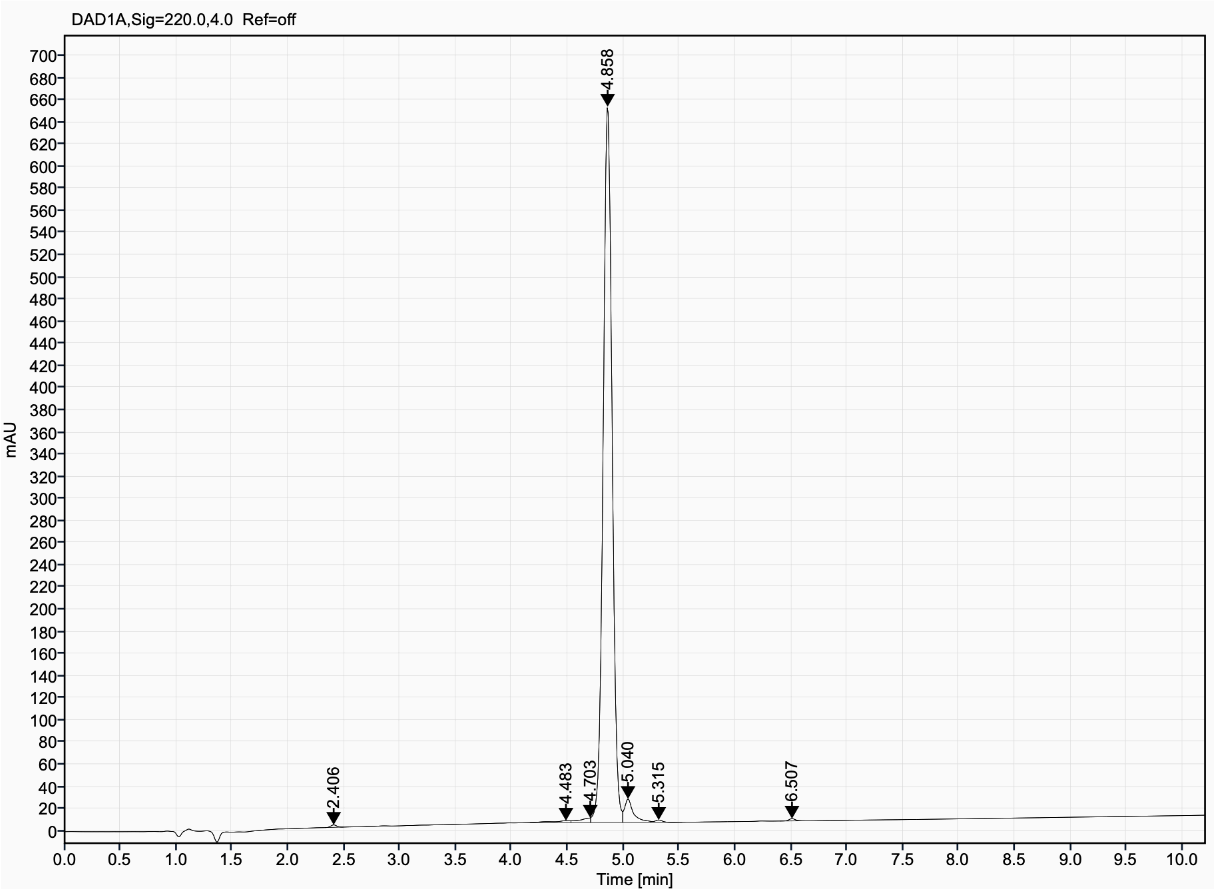

Analytical UPLC and mass spectrum of MDB_D2
Sequence: cyclo[ERVLGIEESEDEDFRVF]

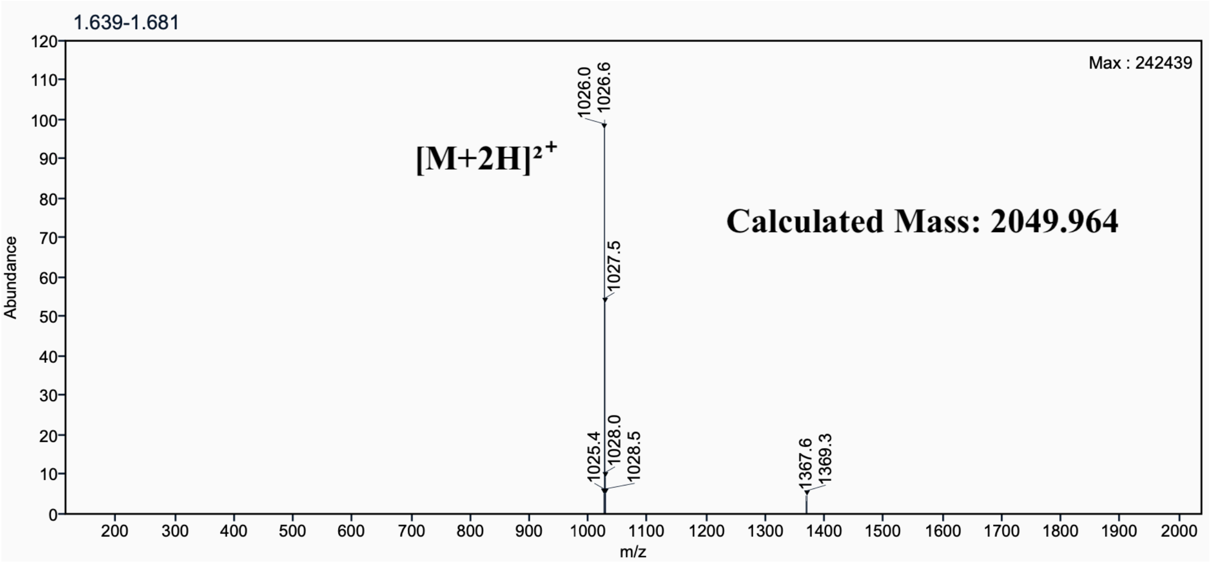

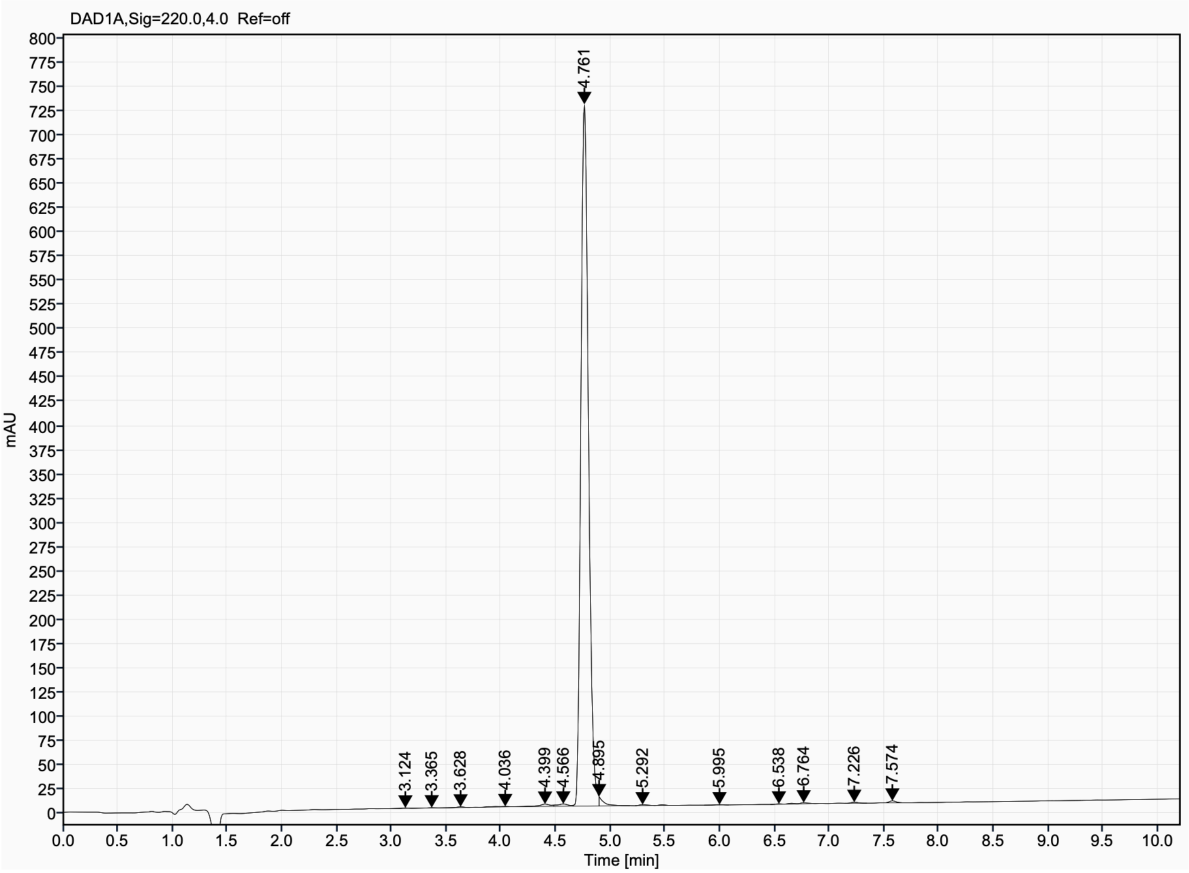

Analytical UPLC and mass spectrum of MDB_D3
Sequence: cyclo[LMEMVGEKYDQNFILRE]

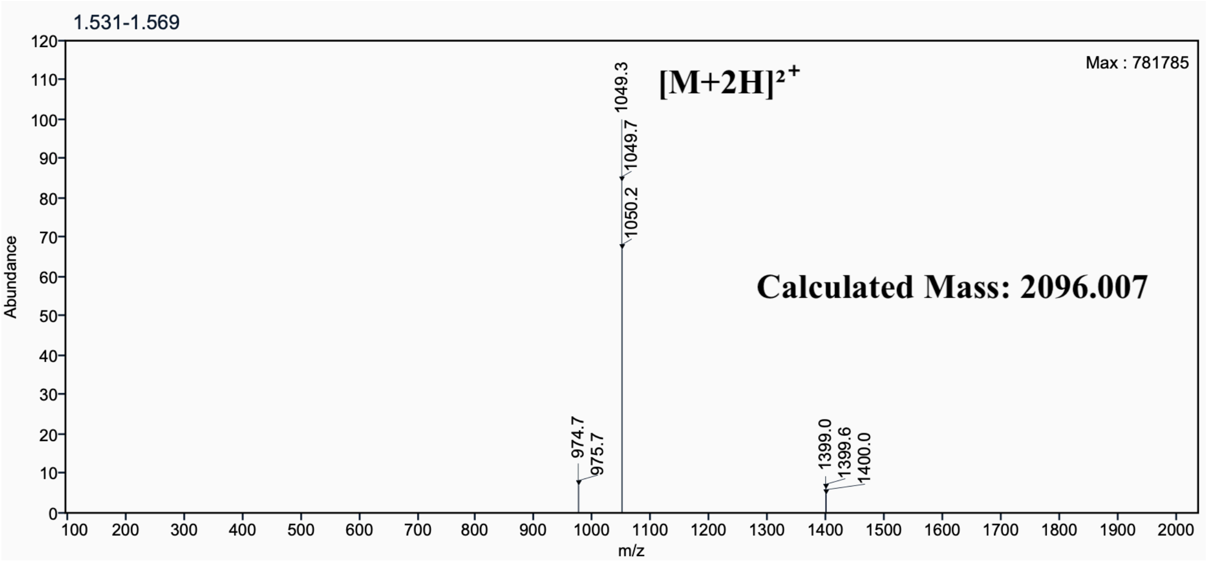

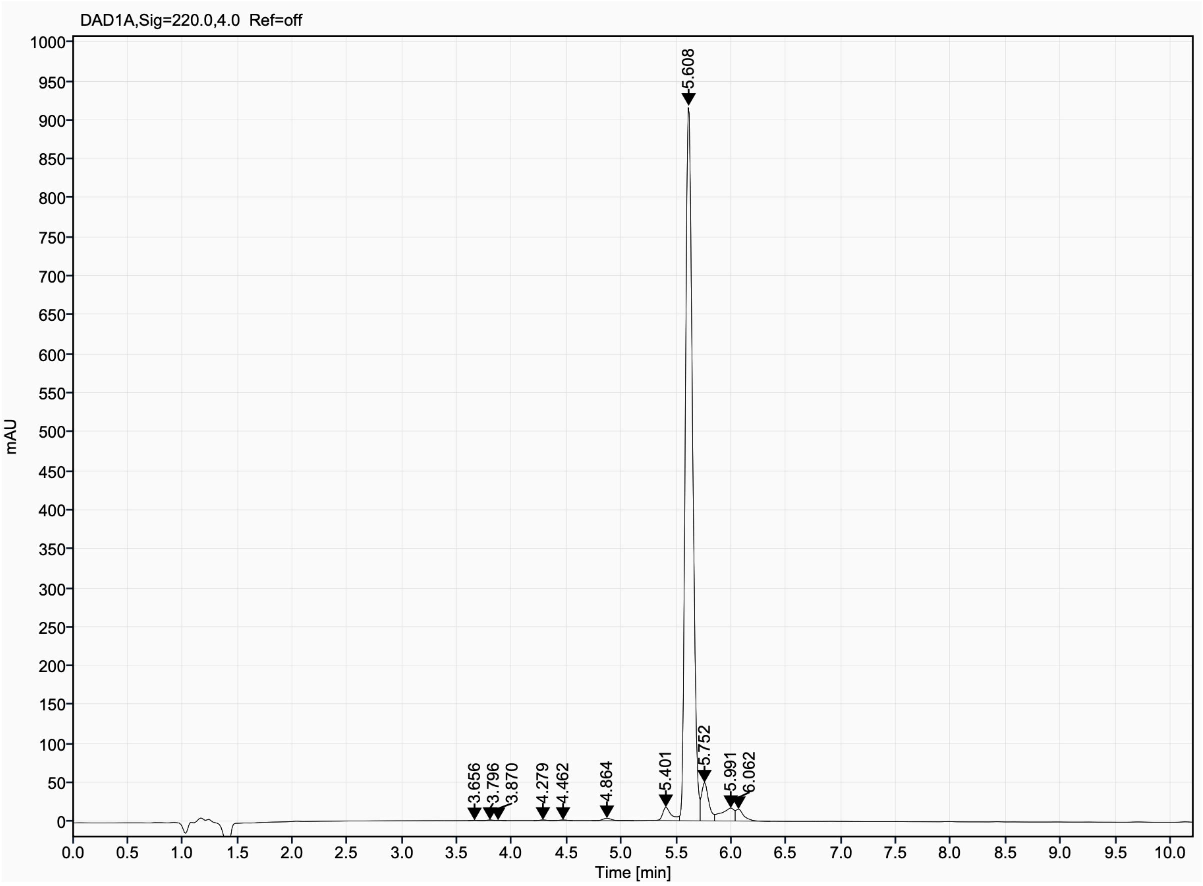

Analytical UPLC and mass spectrum of MDB_D4
Sequence: cyclo[LTDVMGLDIESDDDFKV]

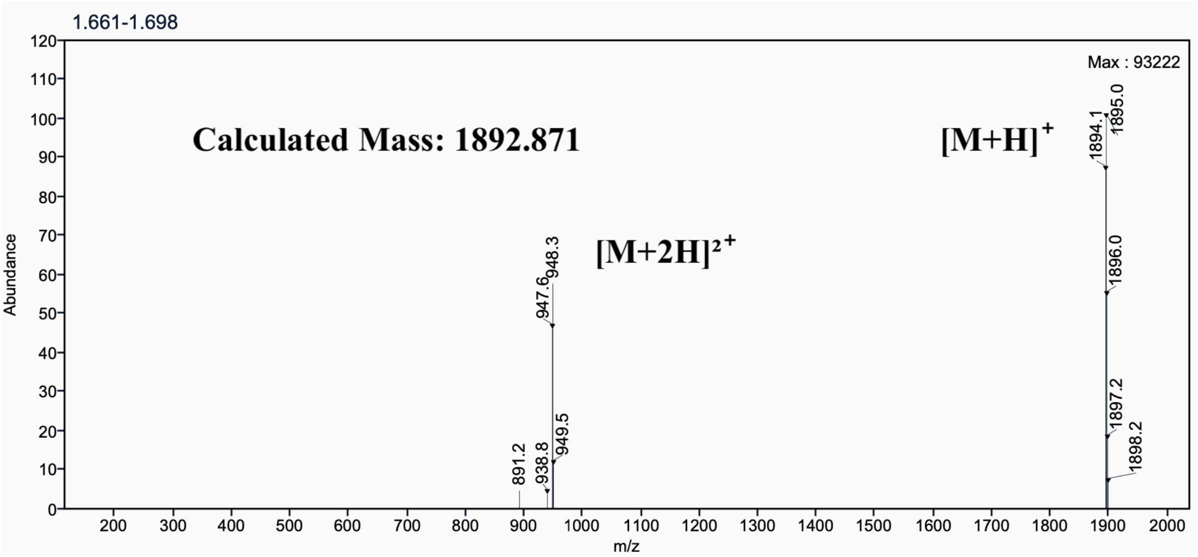

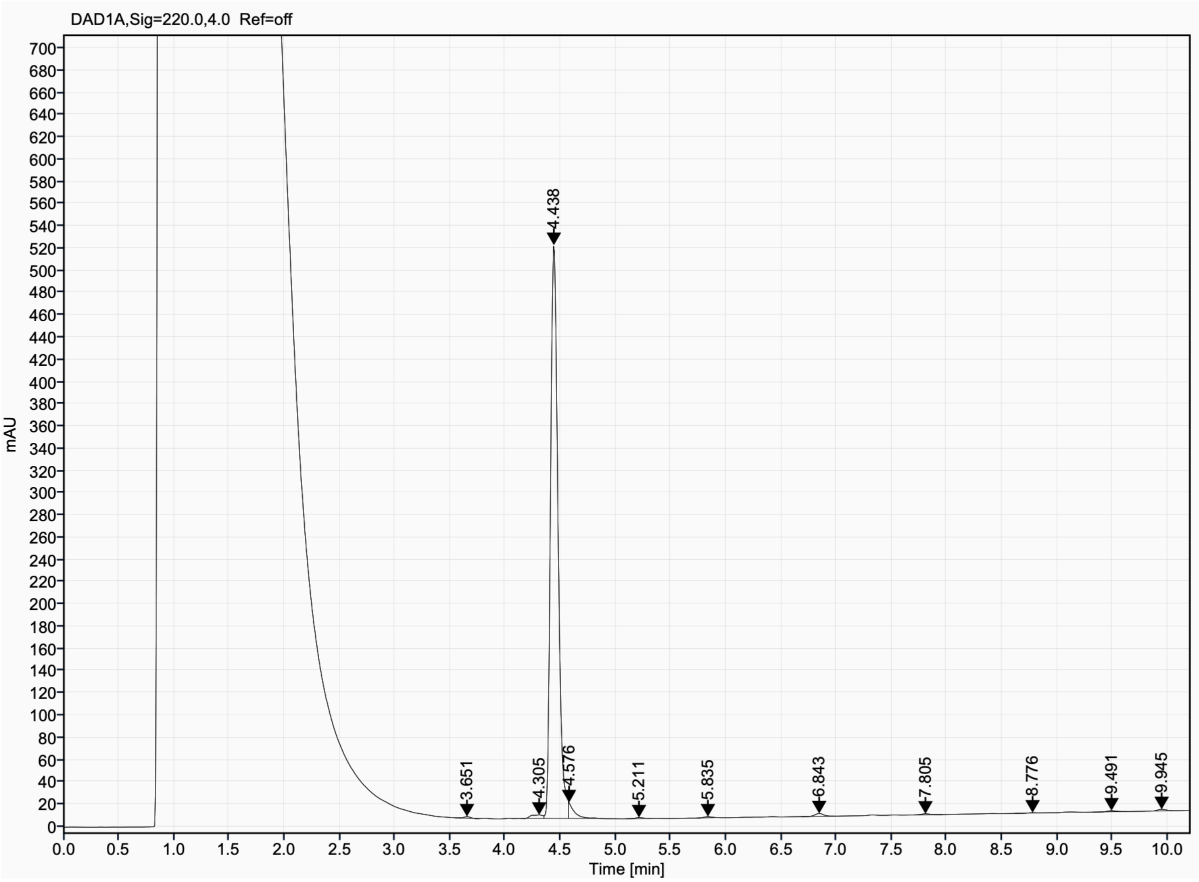

Analytical UPLC and mass spectrum of MDB_D6
Sequence: cyclo[LHDVLGLPLEFDDEDYQV]

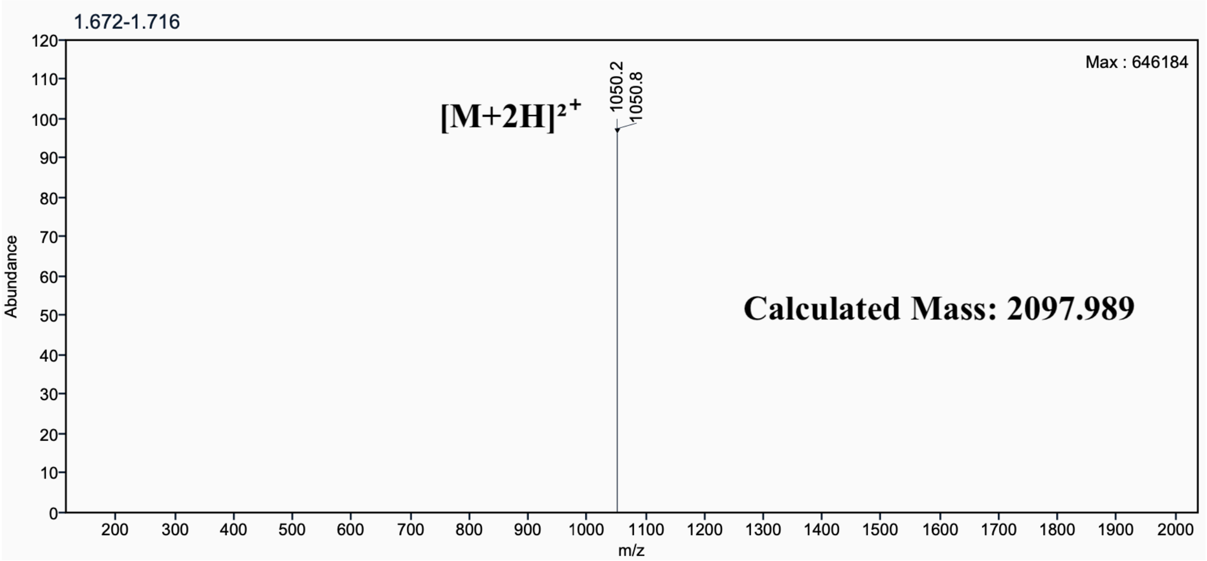

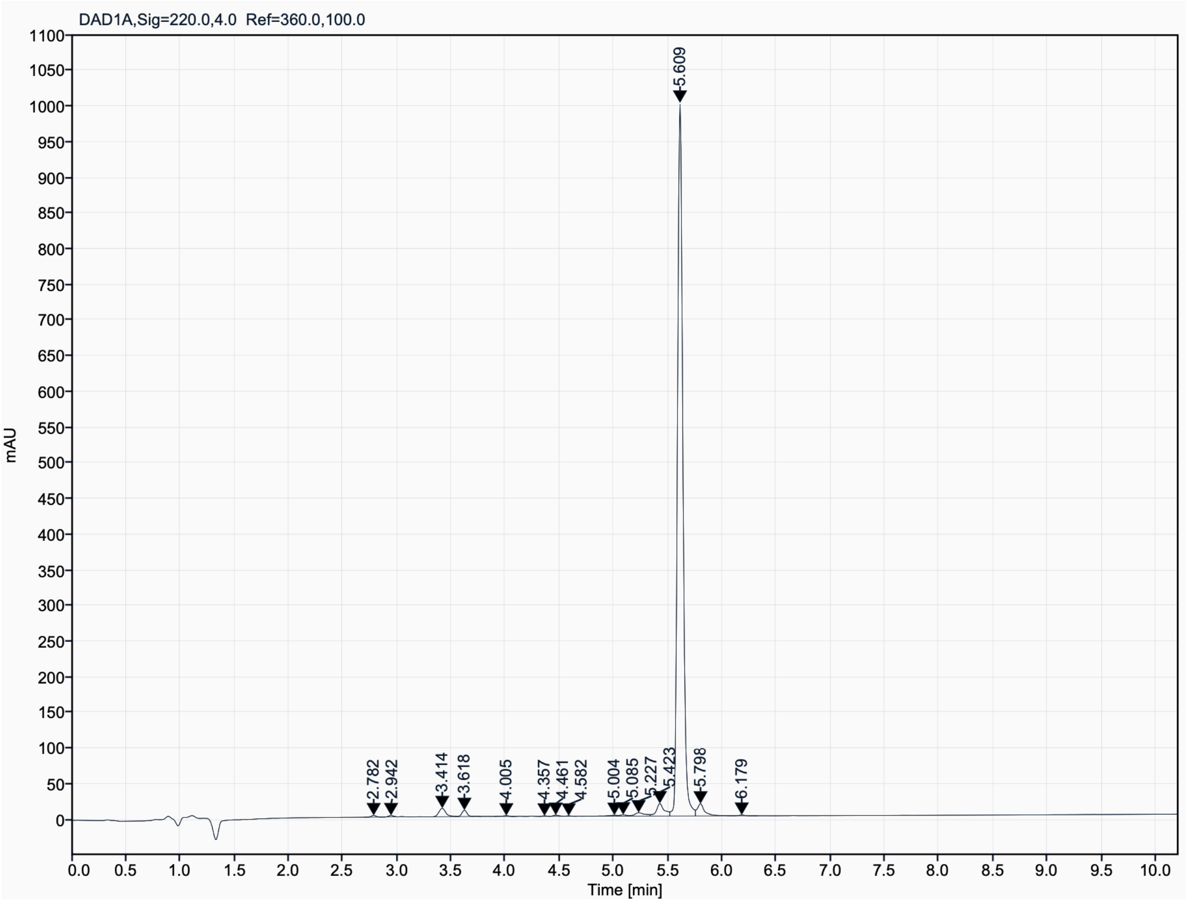

Analytical UPLC and mass spectrum of MDB_D8
Sequence: cyclo[AKNKFEELWNELIDPSRK]

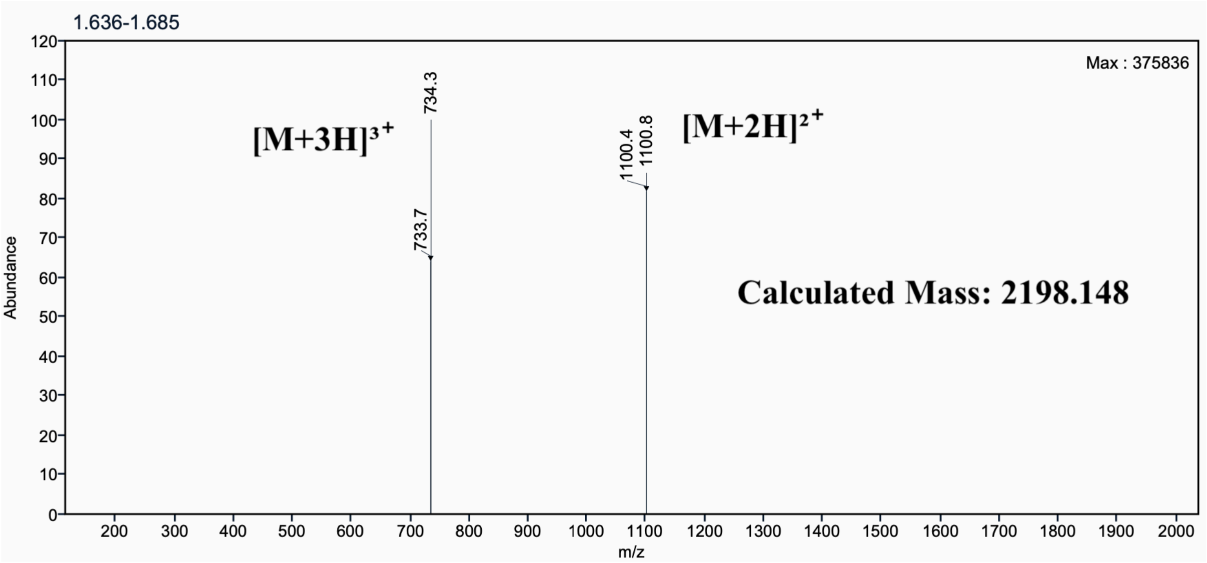

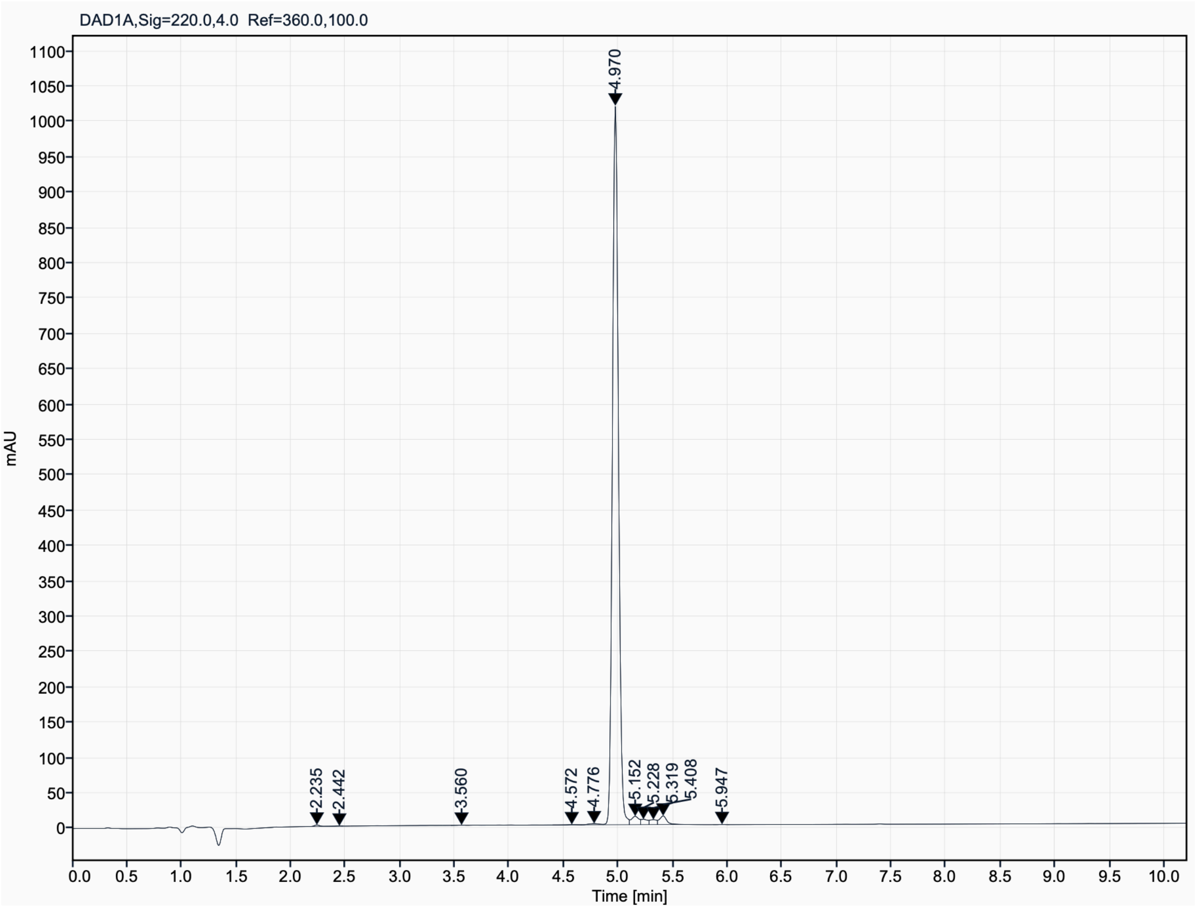

Analytical UPLC and mass spectrum of MDB_D10
Sequence: cyclo[WSKMAGGAPLPGTPFAKE]

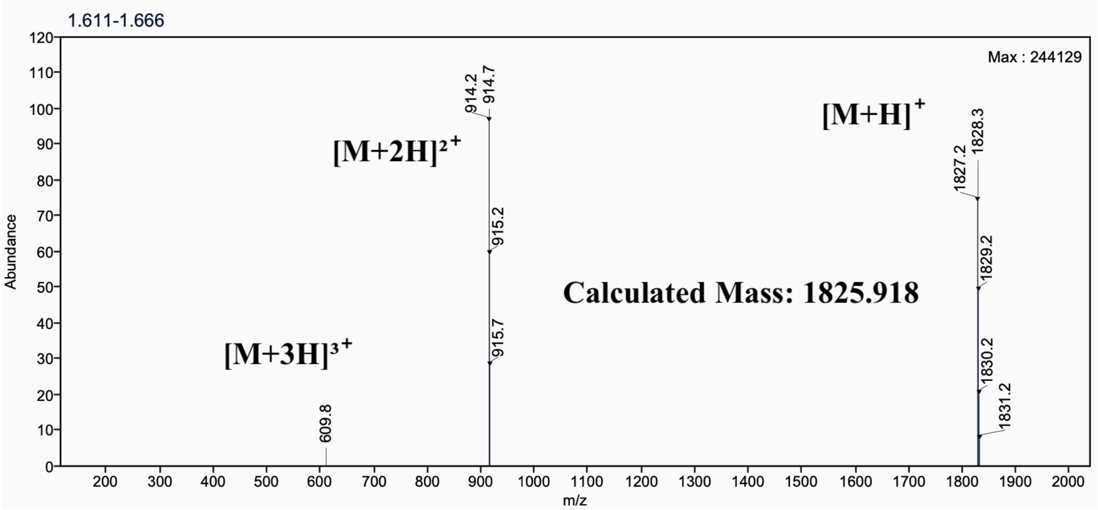

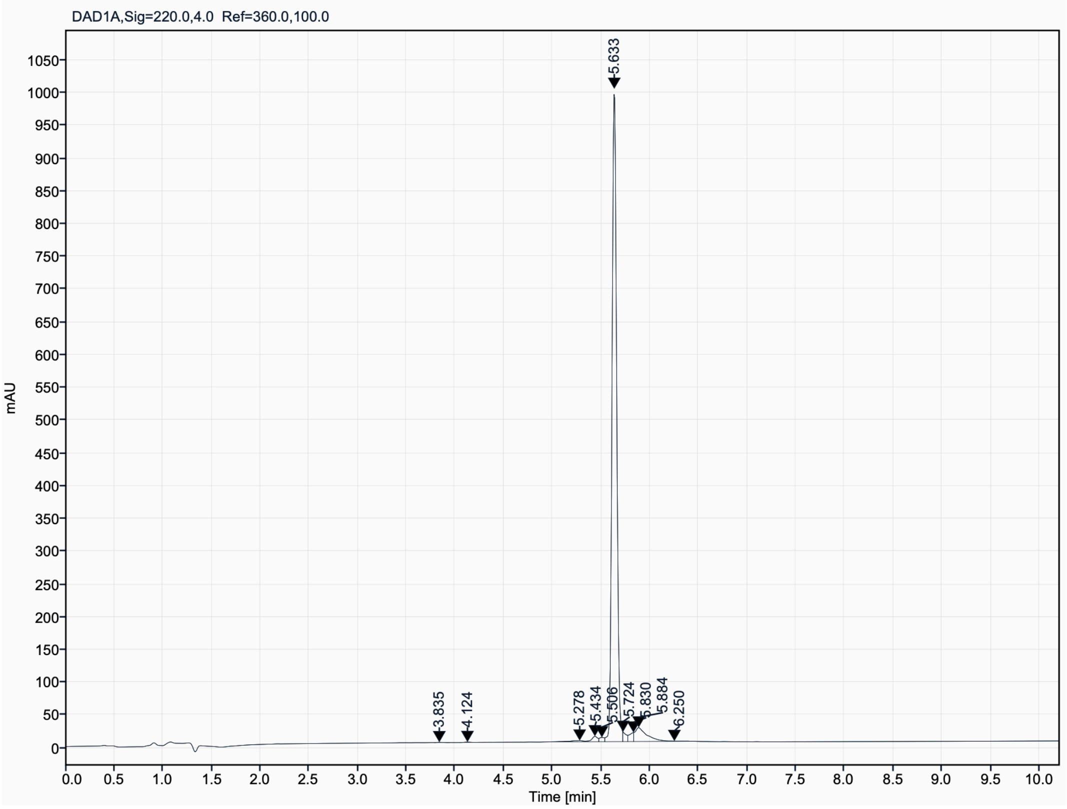

Analytical UPLC and mass spectrum of MDB_D11
Sequence: cyclo[EWQKMVGLDLETDSEFAK]

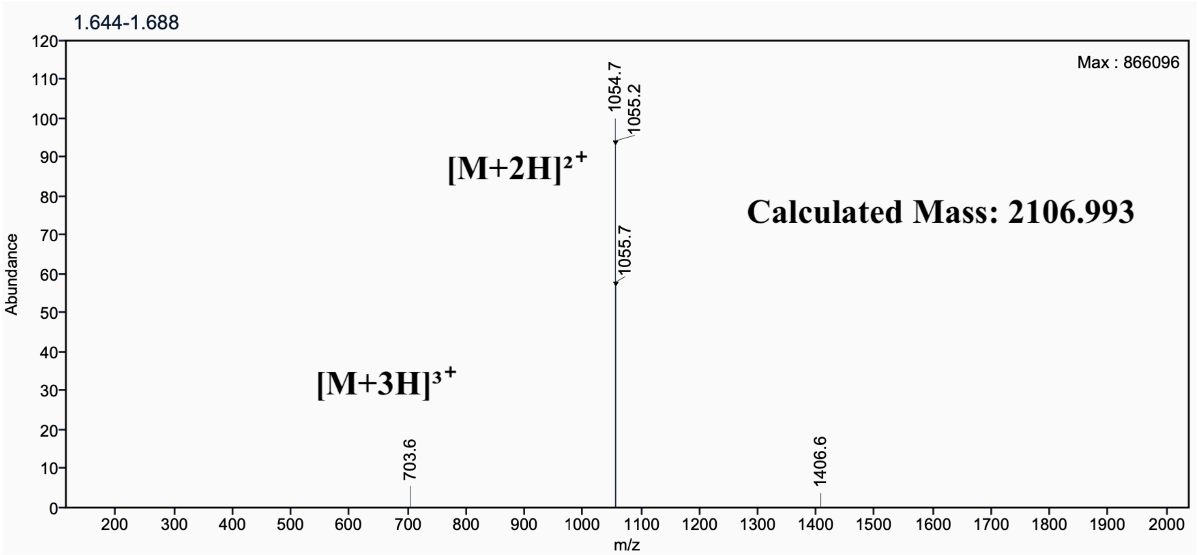

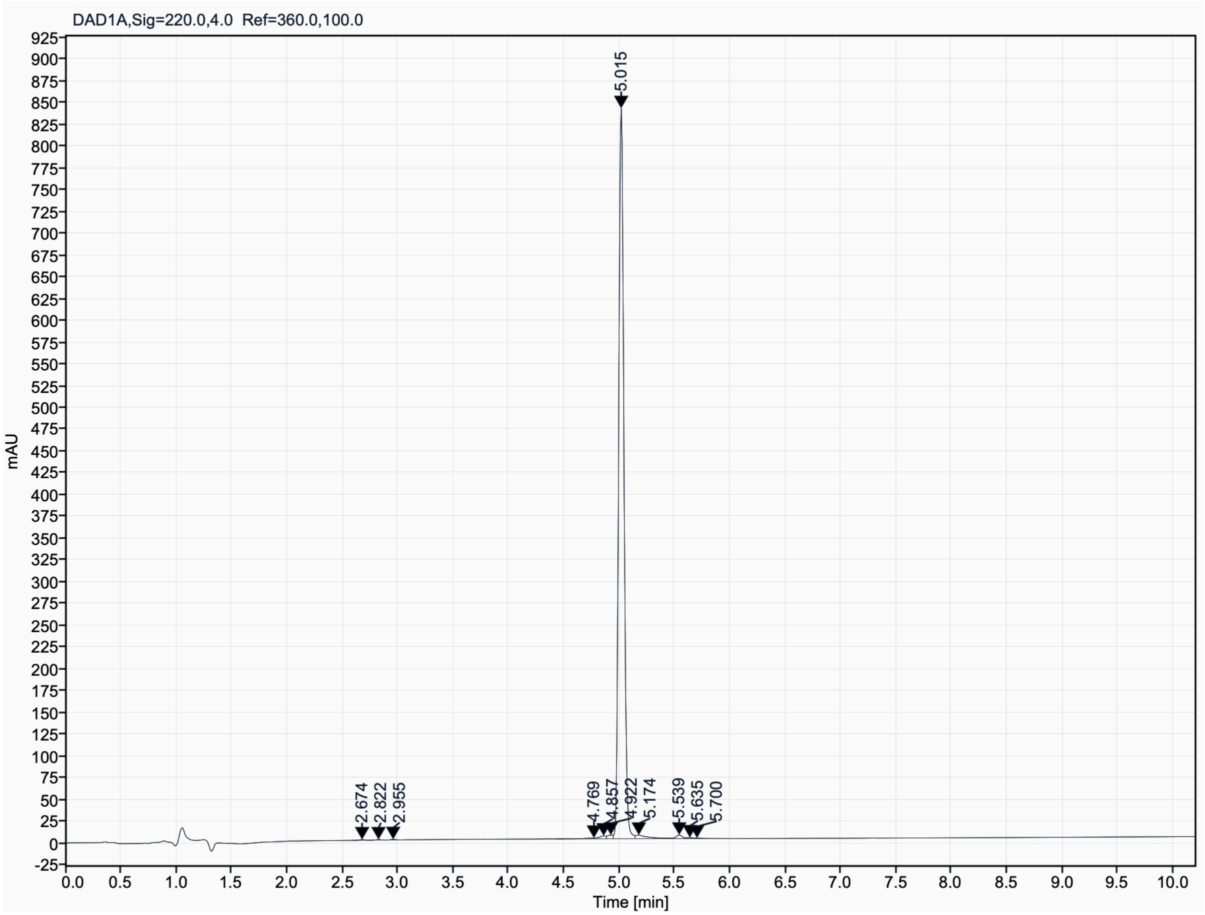

**GAB peptides Mass Spec and HPLC**

Analytical HPLC and mass spectrum of GAB_D1
Sequence: cyclo[ETGEVENIDGVEIIYP]

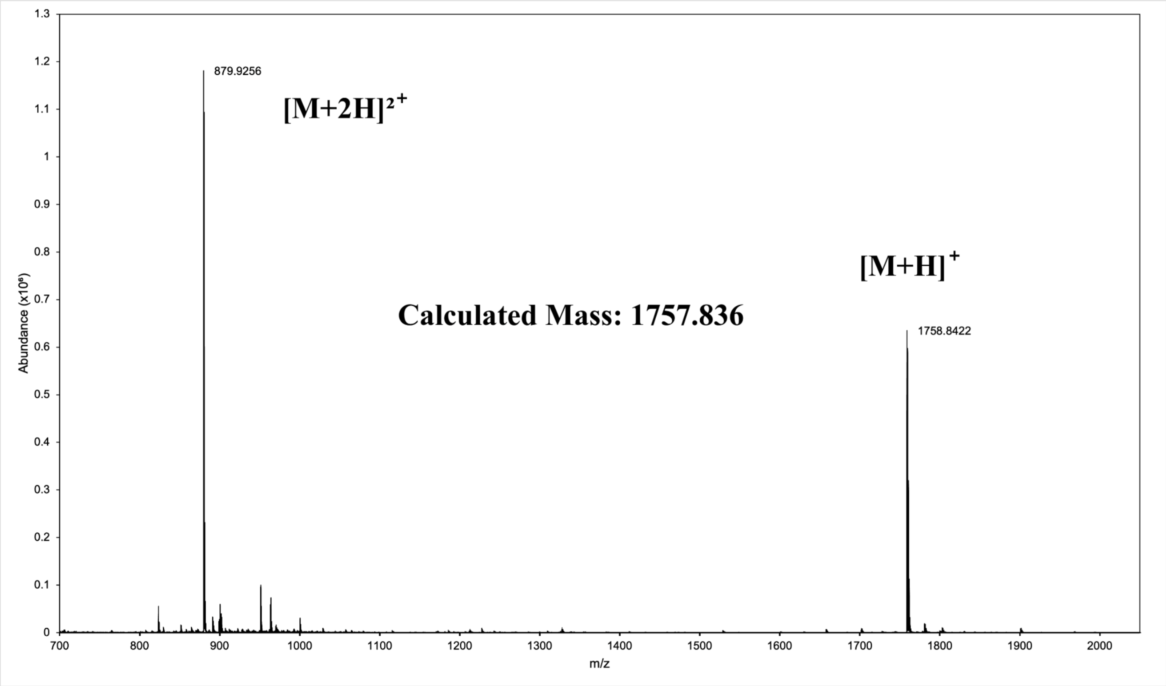

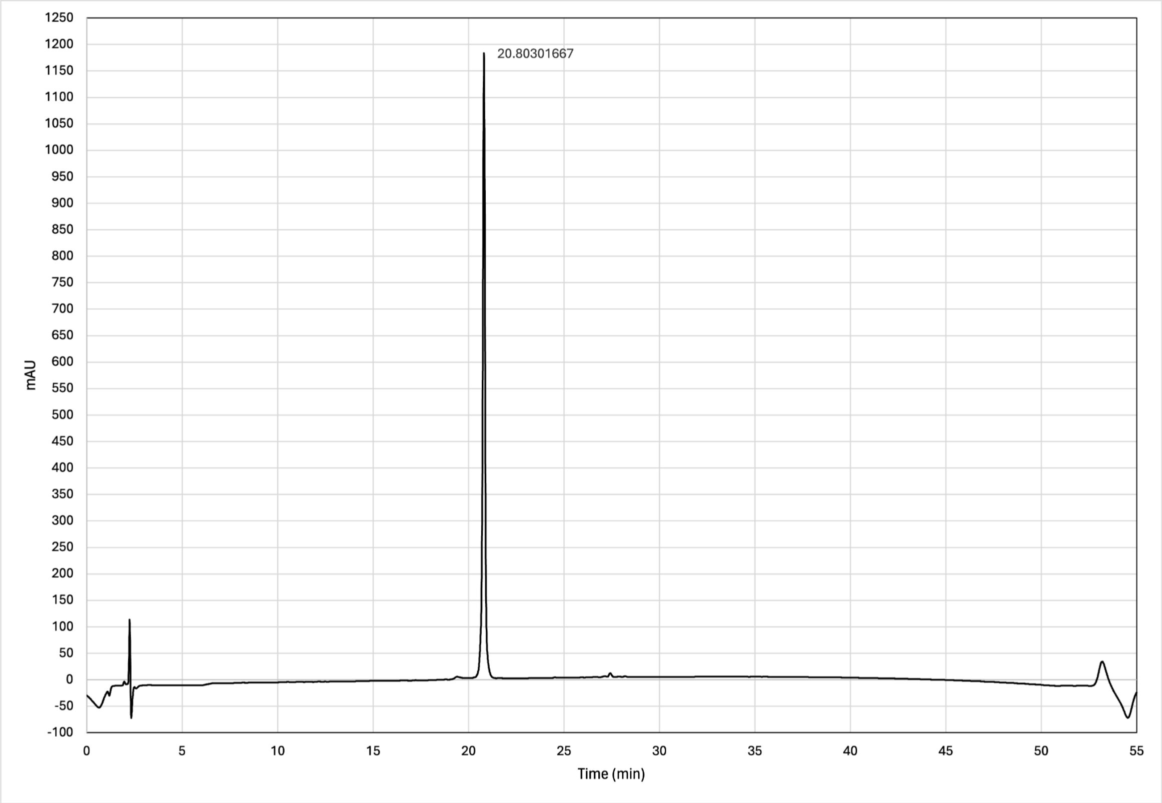

Analytical HPCL and mass spectrum of GAB_D8
Sequence: cyclo[GSEYEEDGWTVLEPD]

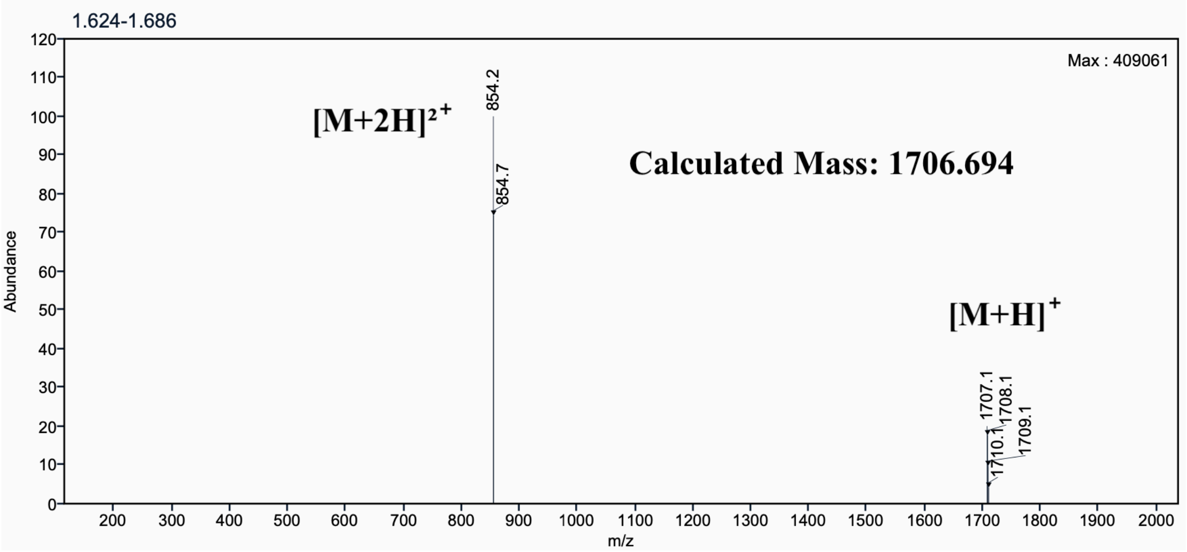

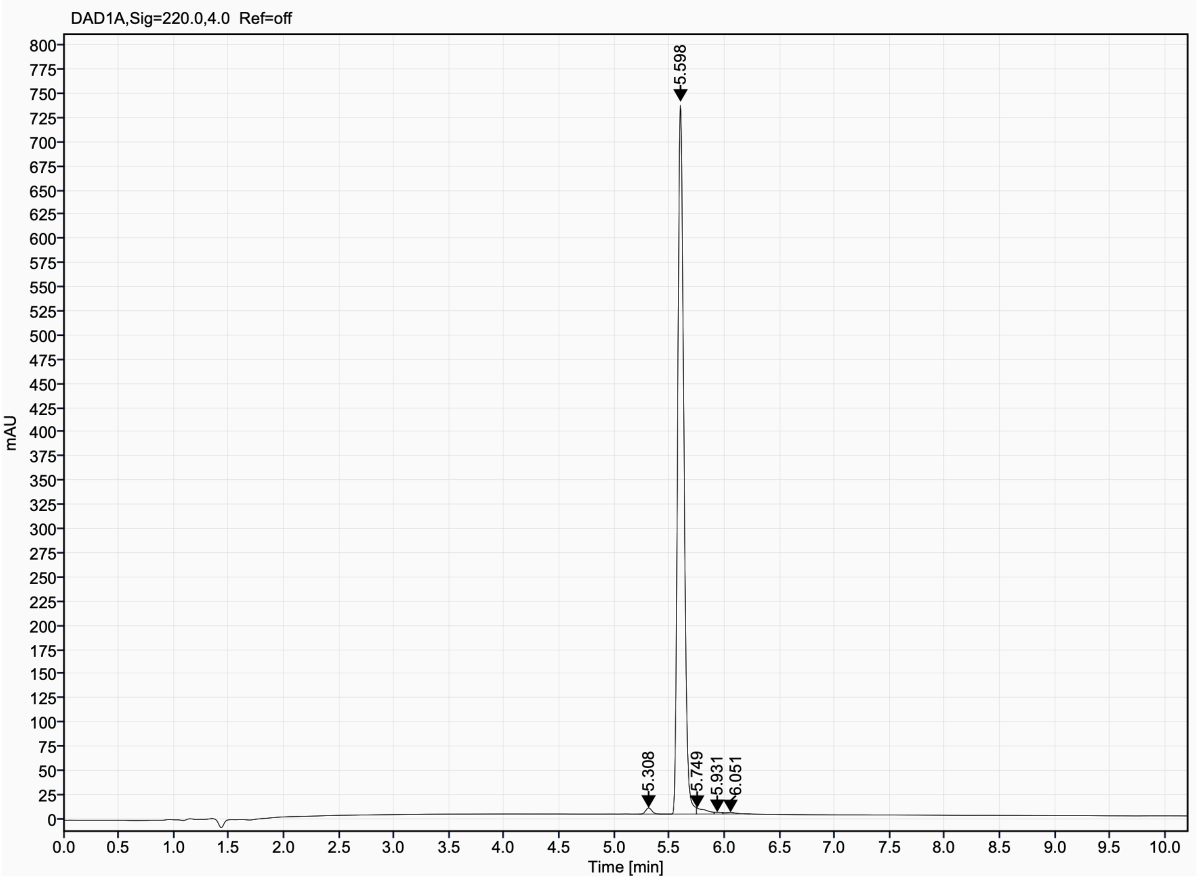

Analytical HPLC and mass spectrum of GAB_D23
Sequence: cyclo[LEDGWVDIETGKE]

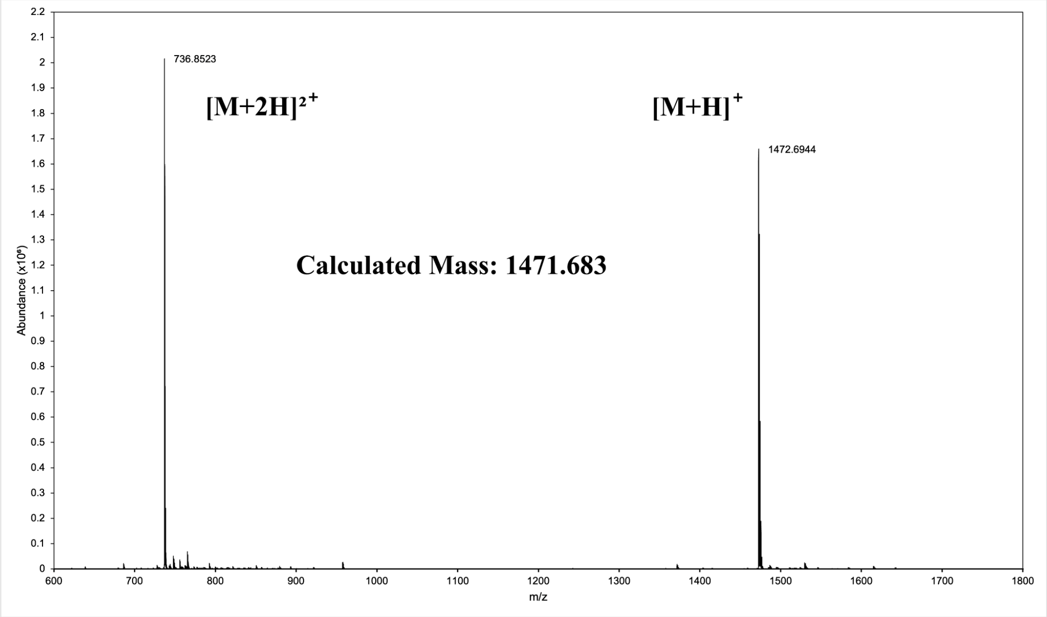

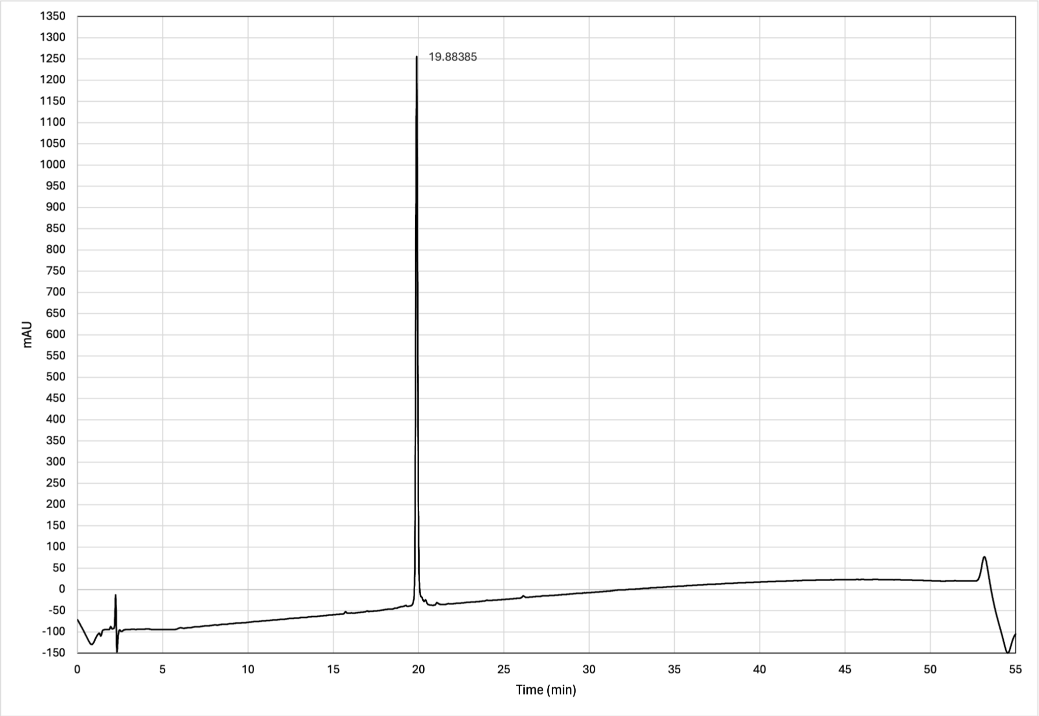

Analytical HPLC and mass spectrum of GAB_D26
Sequence: cyclo[LDTGEVYKAPNGQEVIE]

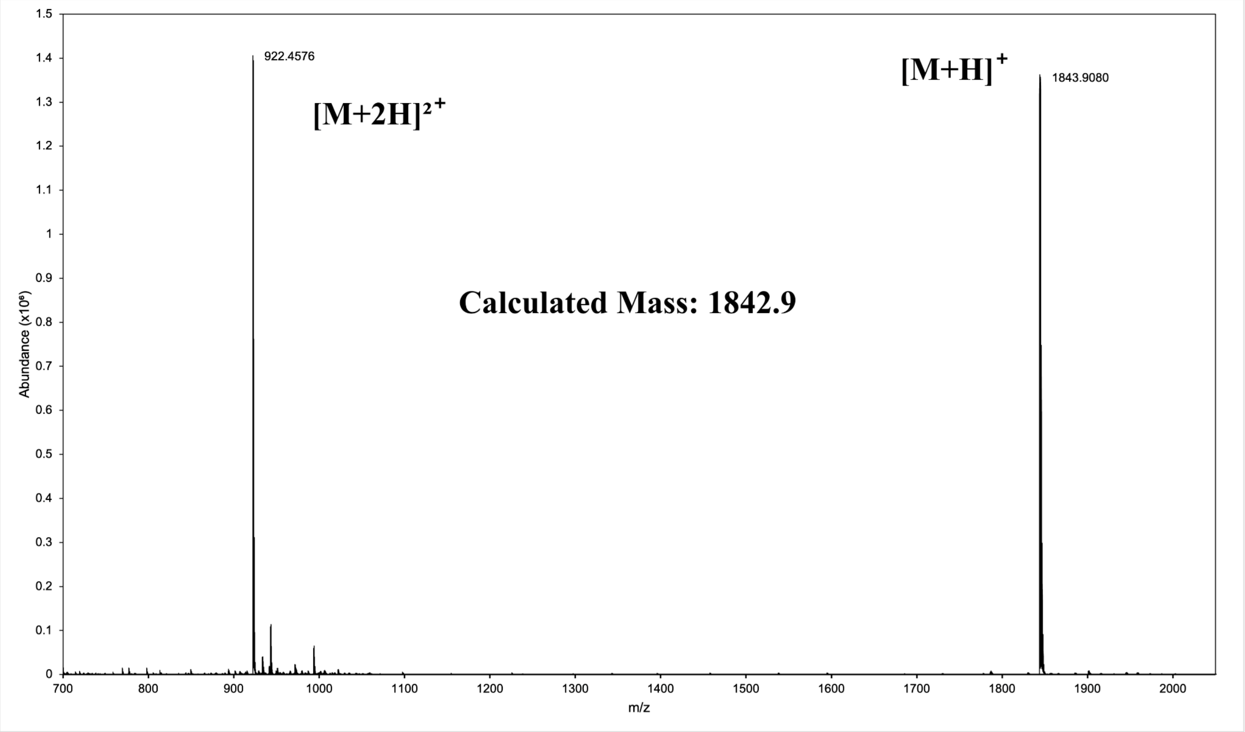

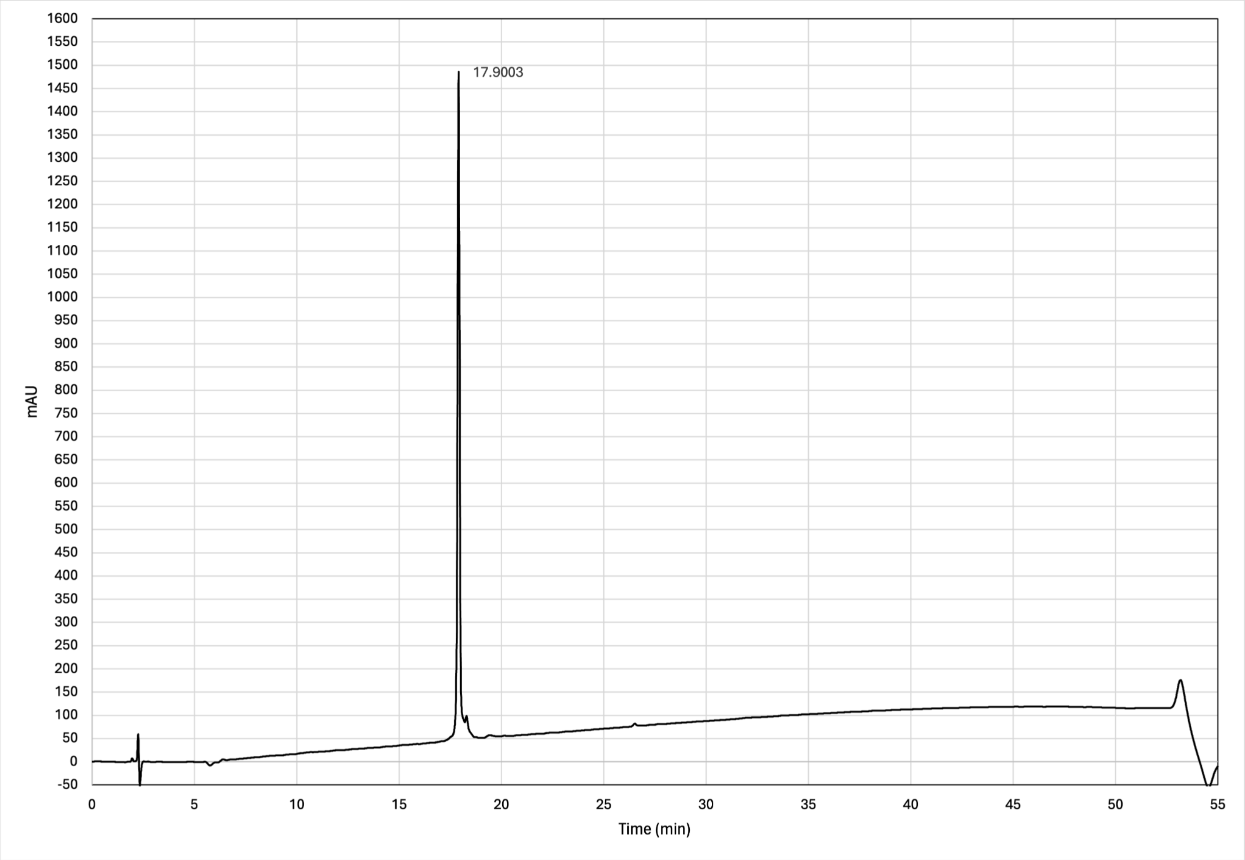

Analytical UPLC and mass spectrum of GAB_D27

Sequence: cyclo[IDIDTEEEVMPGV]

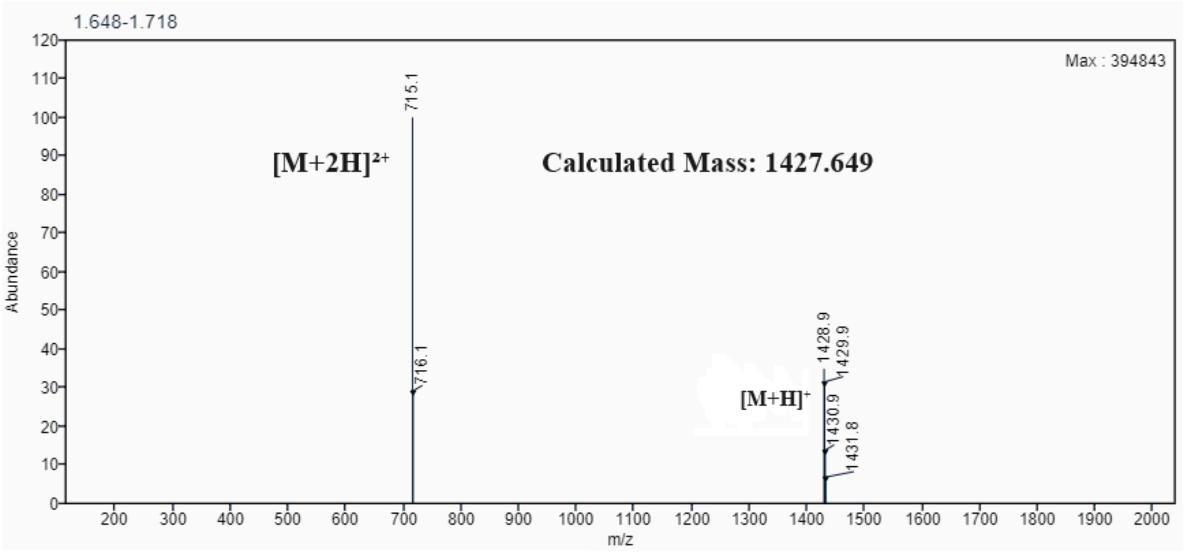

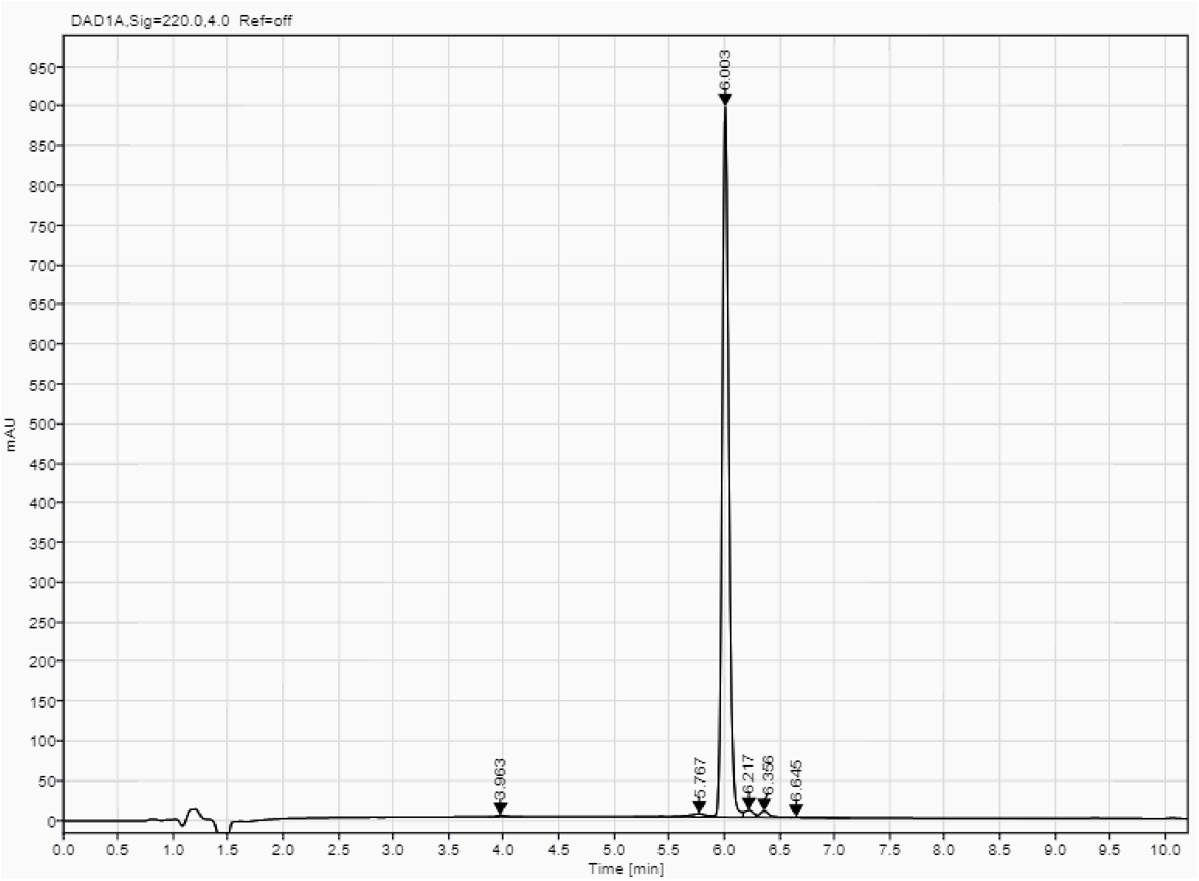

Analytical UPLC and mass spectrum of GAB_D28

Sequence: cyclo[VMPGIIDIDTEEE]

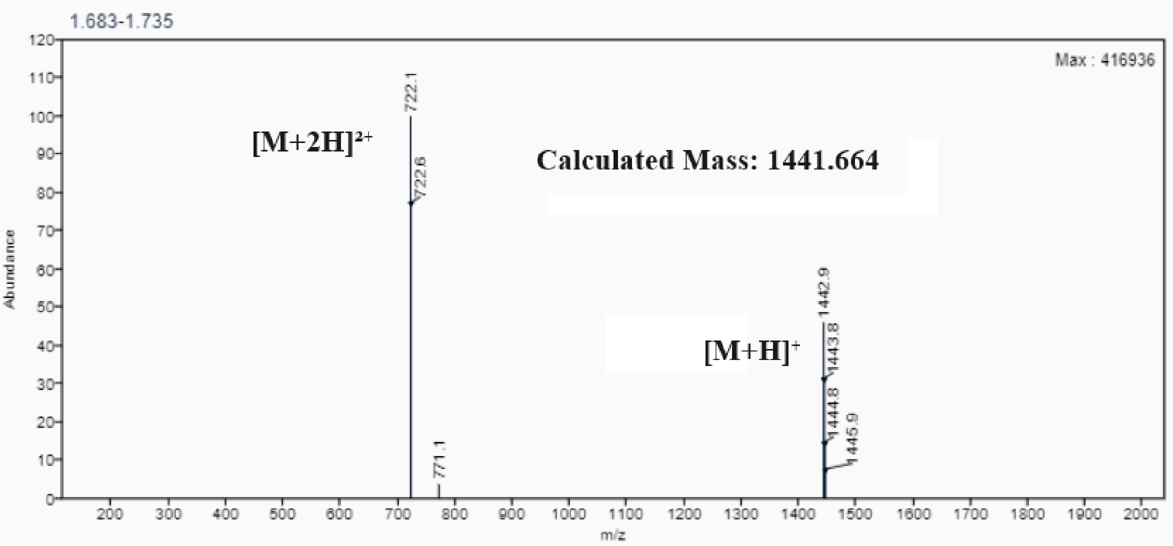

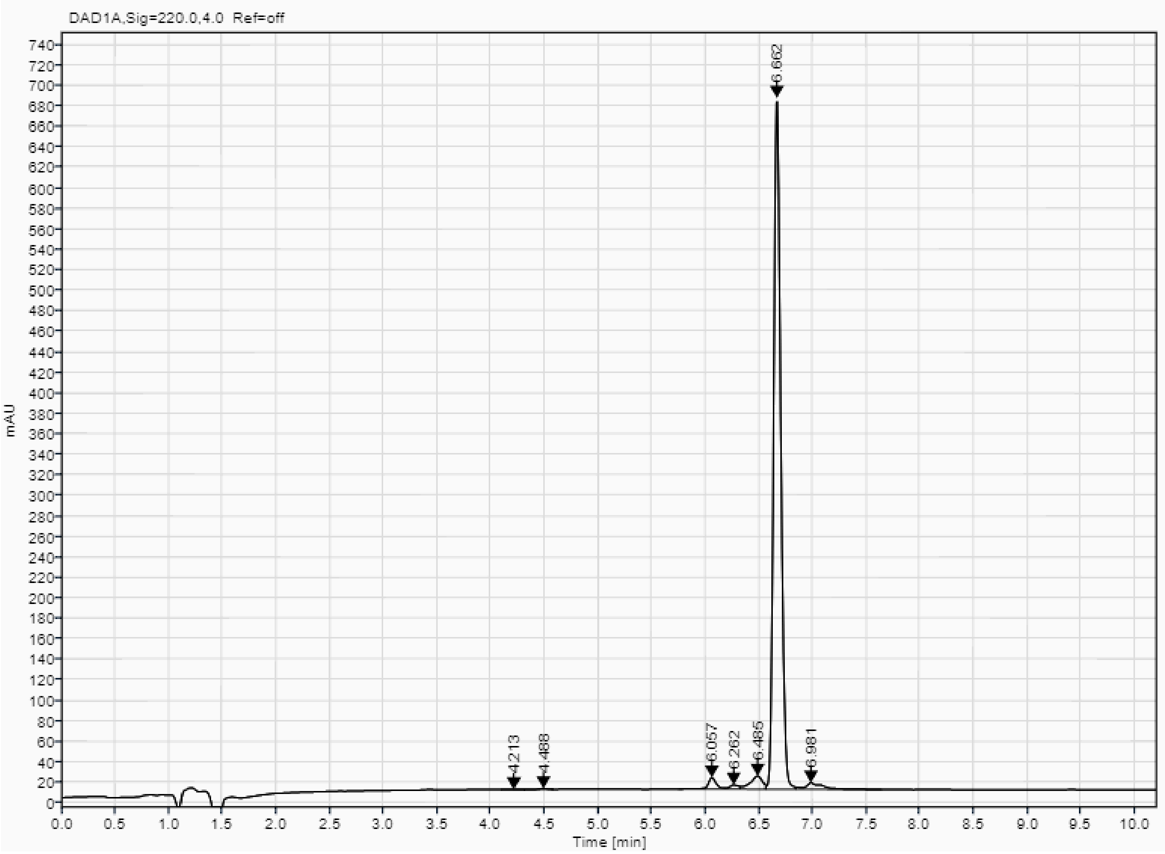

**RBB peptides Mass Spec and HPLC**

Analytical UPLC and mass spectrum of RBB_D1
Sequence: cyclo[SSLPEYVRDVLGV]

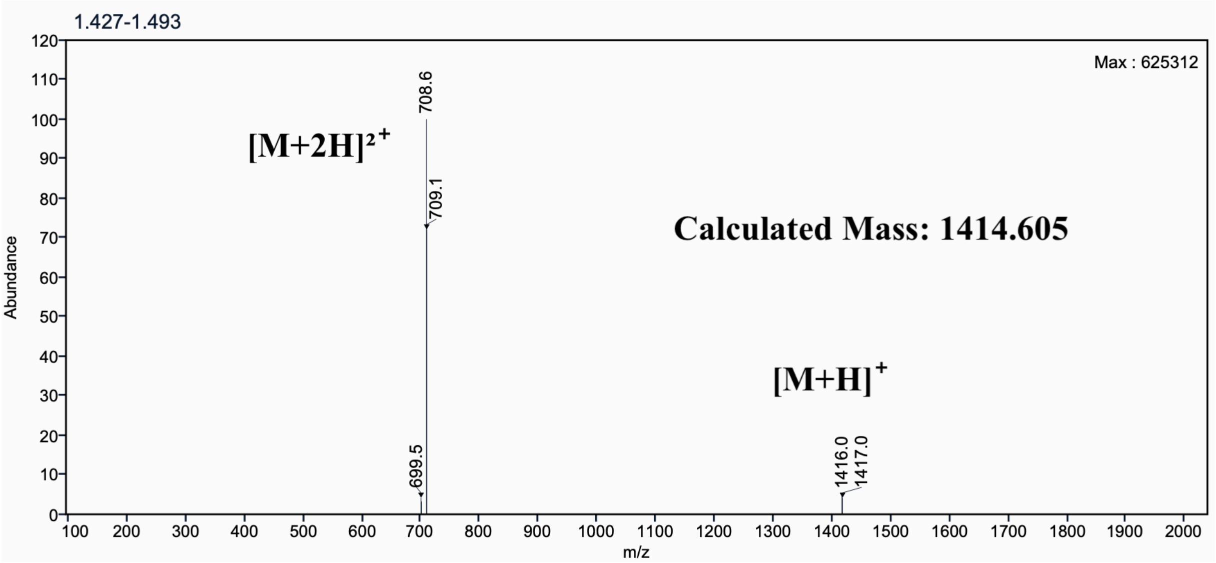

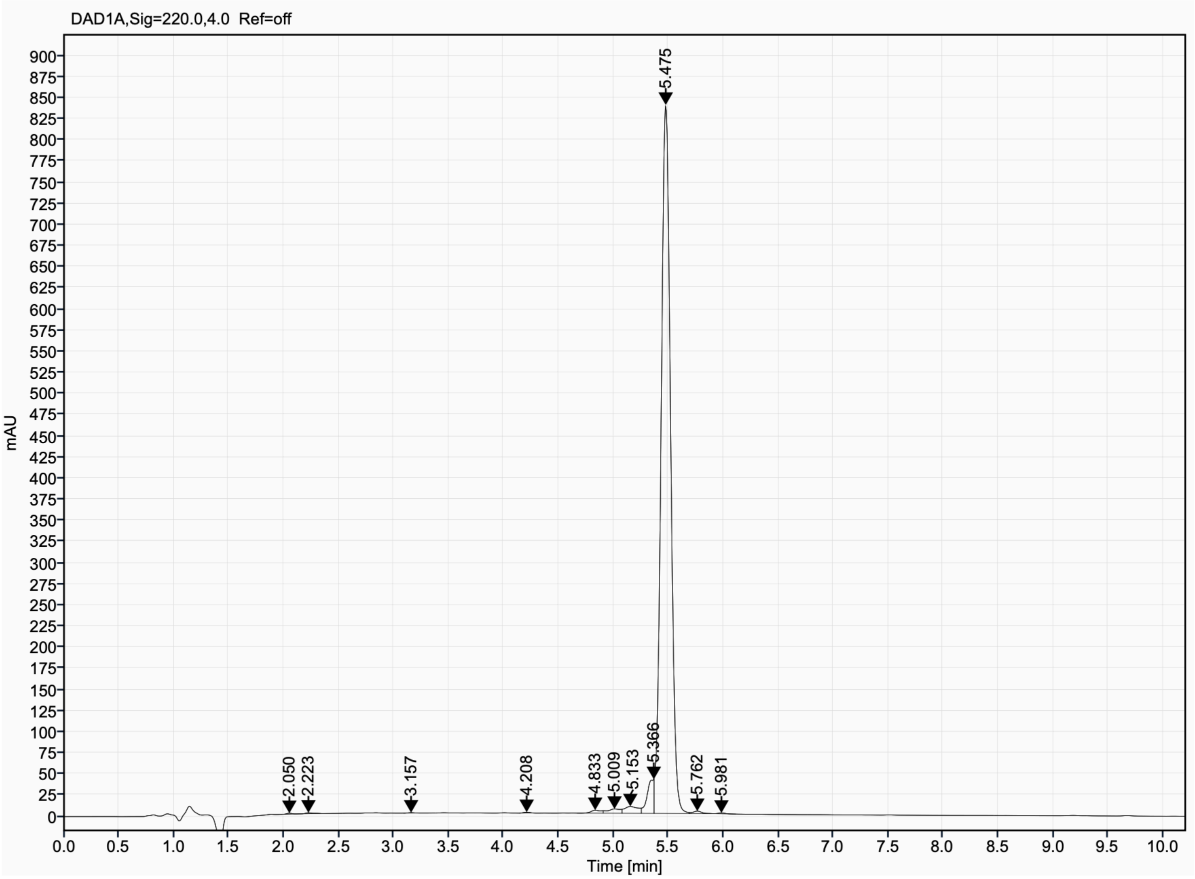

Analytical UPLC and mass spectrum of RBB_D2
Sequence: cyclo[GKPGDSPWEKIESVVFG]

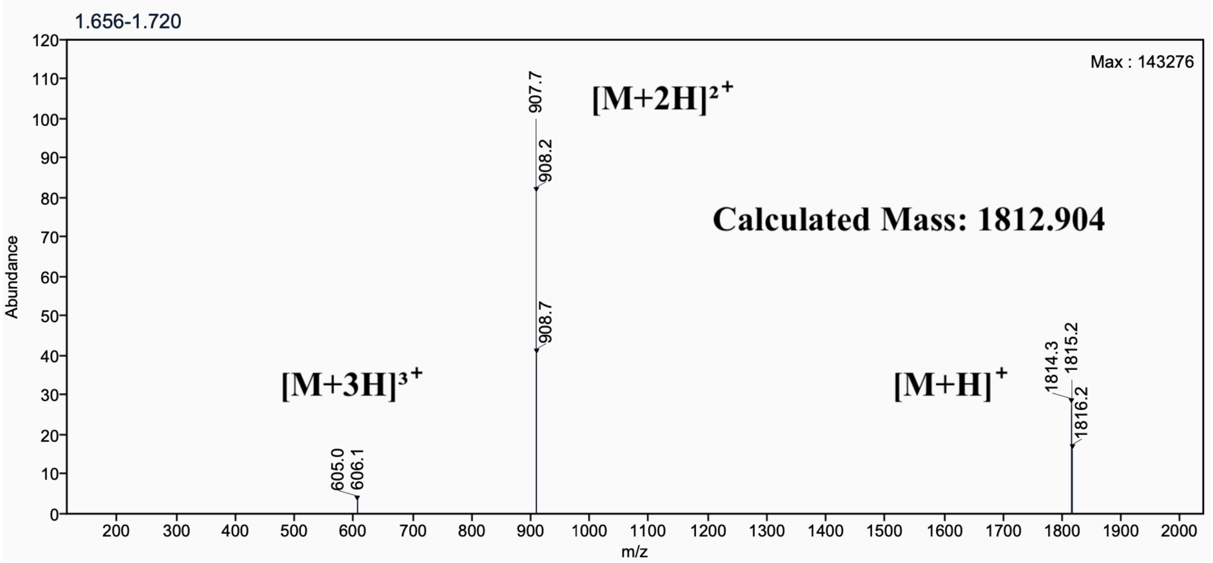

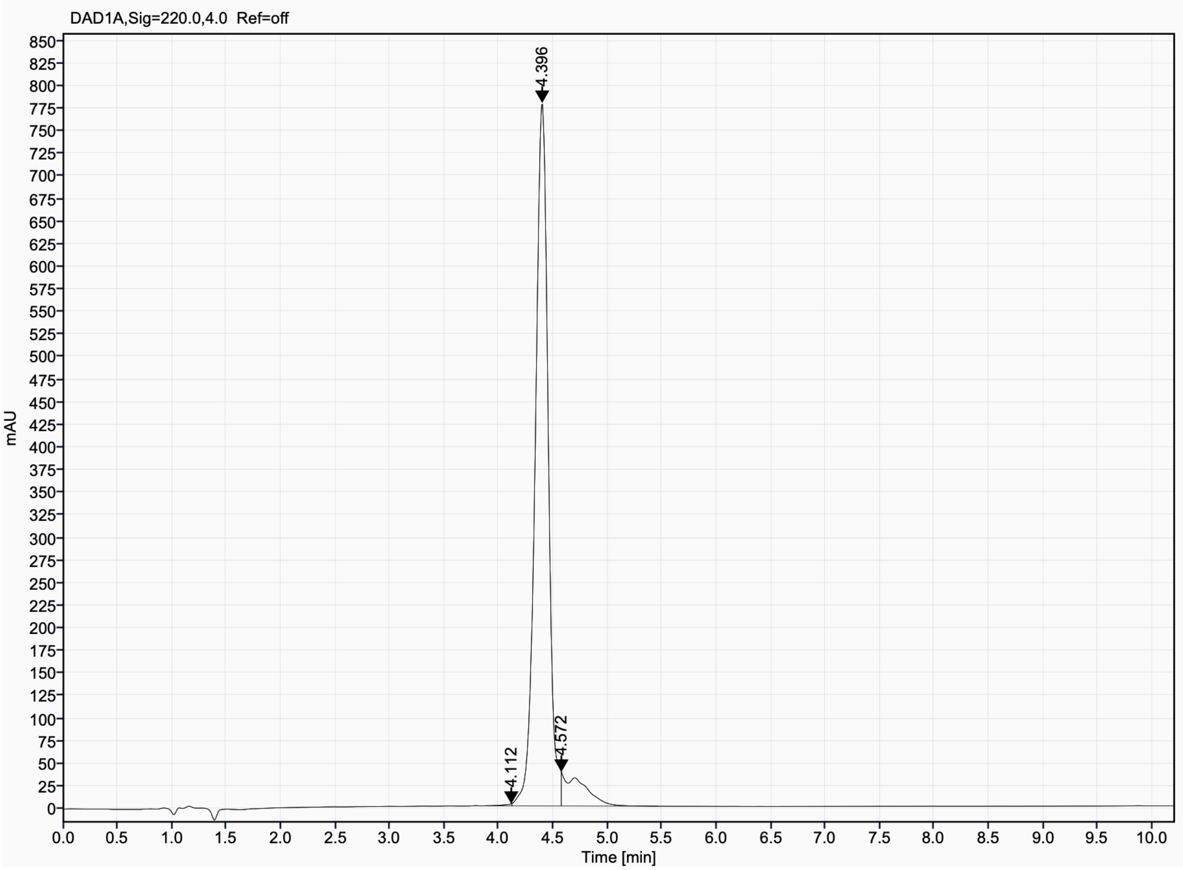

Analytical UPLC and mass spectrum of RBB_D3
Sequence: cyclo[PDPAIPENVQKVFKEHML]

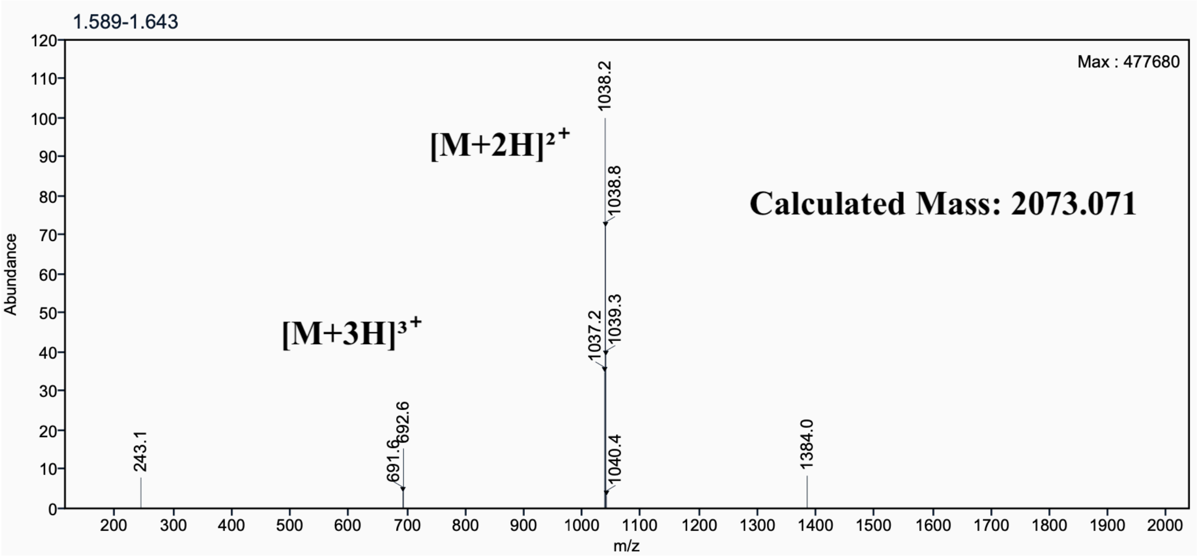

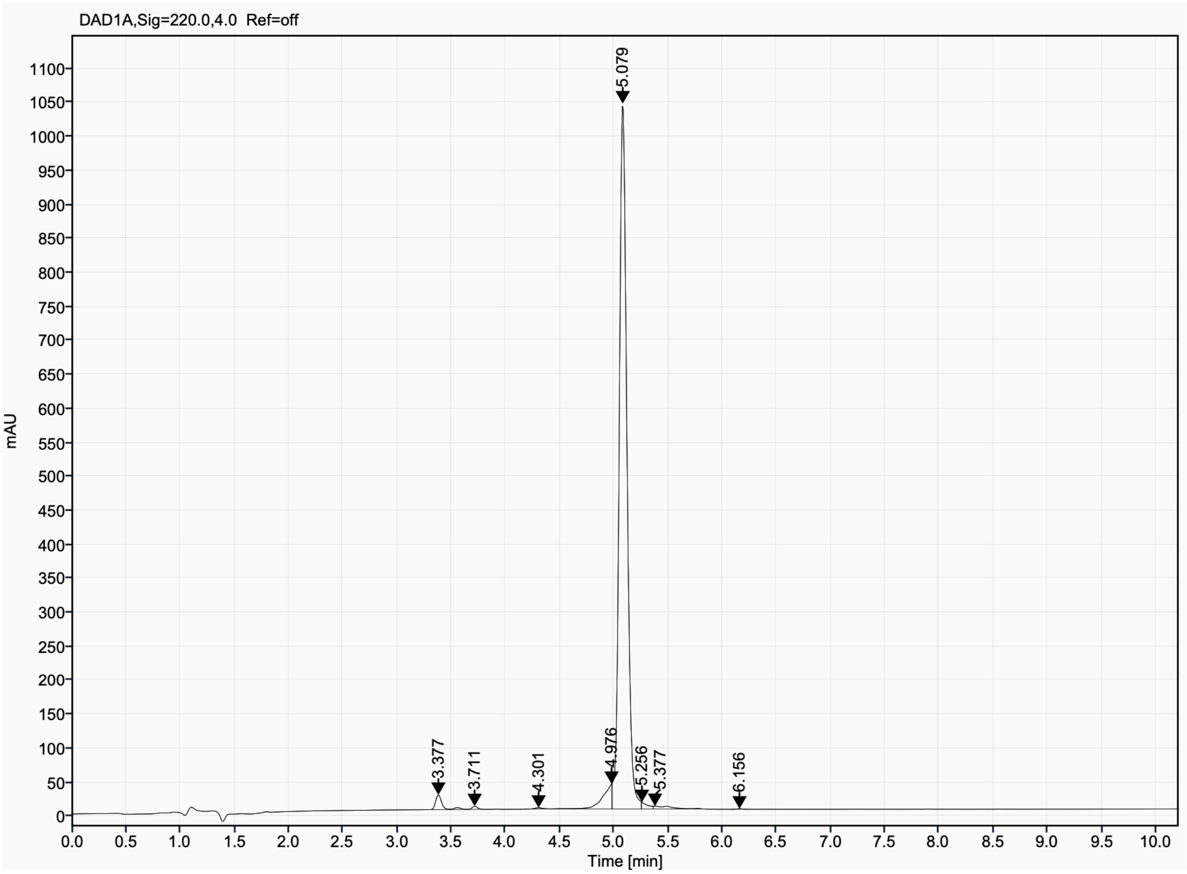

Analytical UPLC and mass spectrum of RBB_D4
Sequence: cyclo[PDPAFPENVQKFKDKIE]

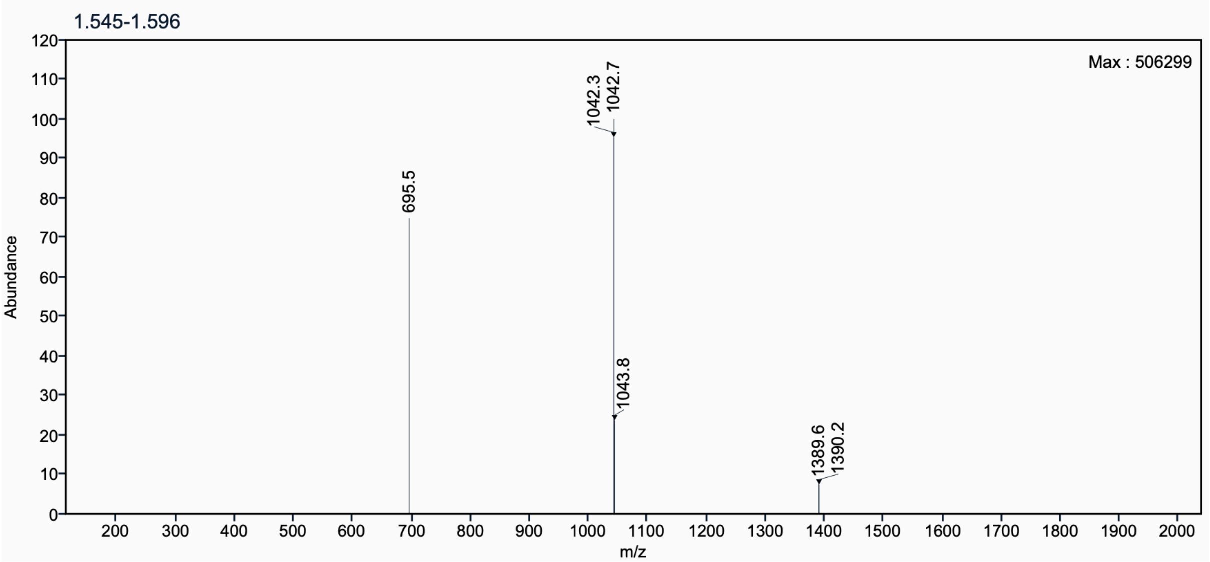

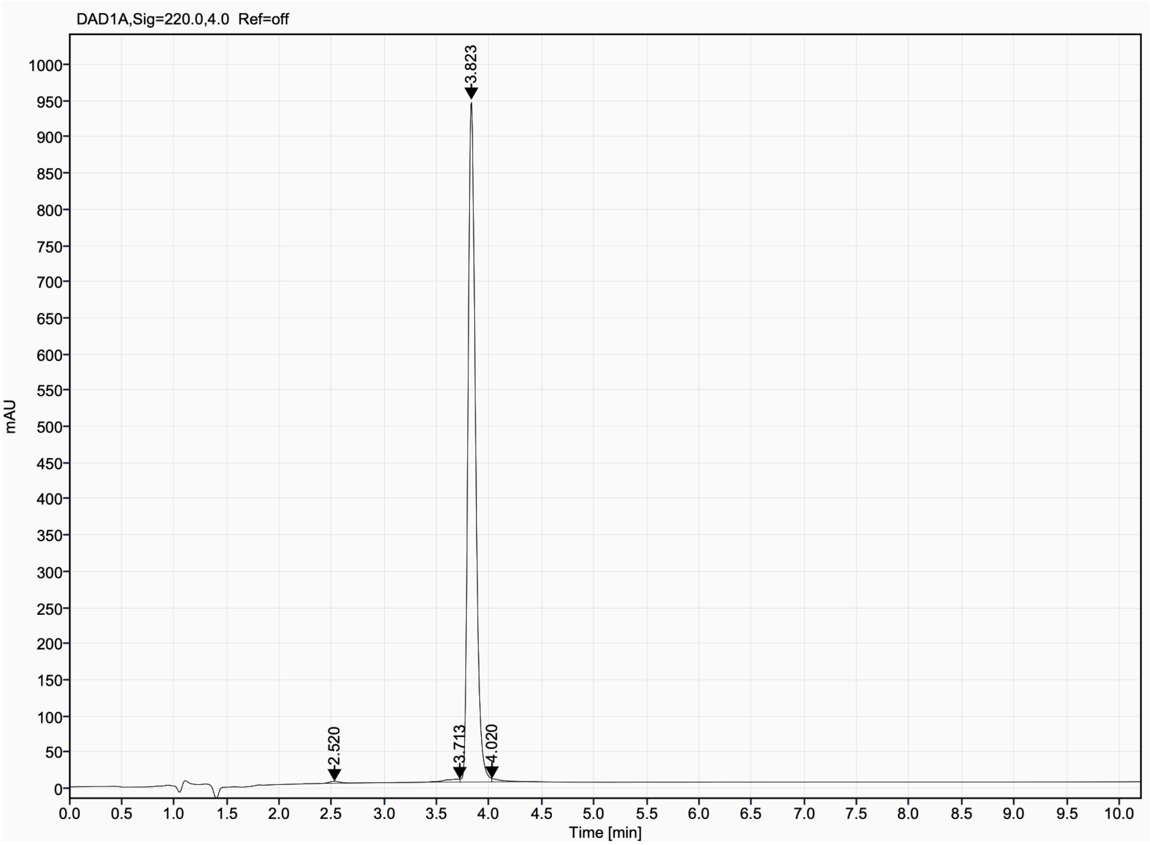

Analytical UPLC and mass spectrum of RBB_D5
Sequence: cyclo[PDPAFPESIHKVFKDKII]

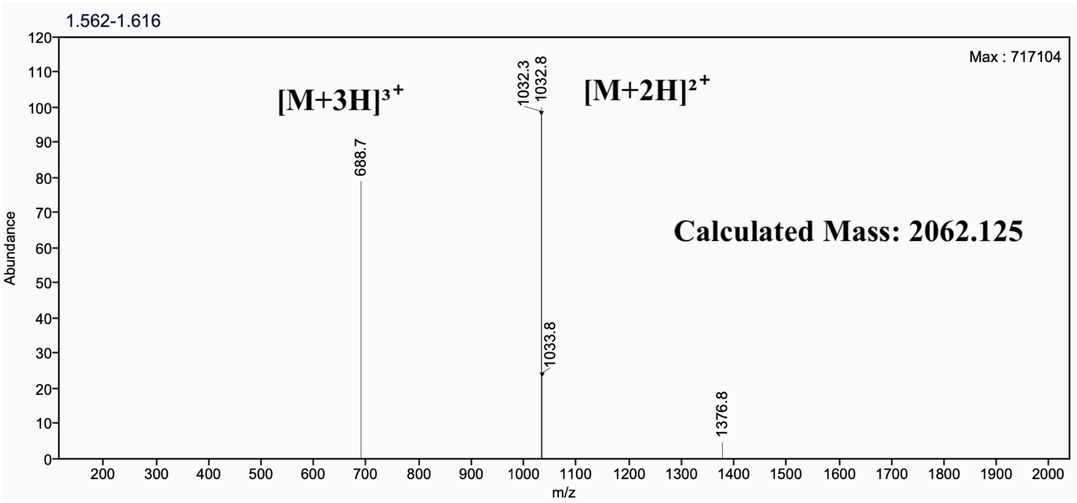

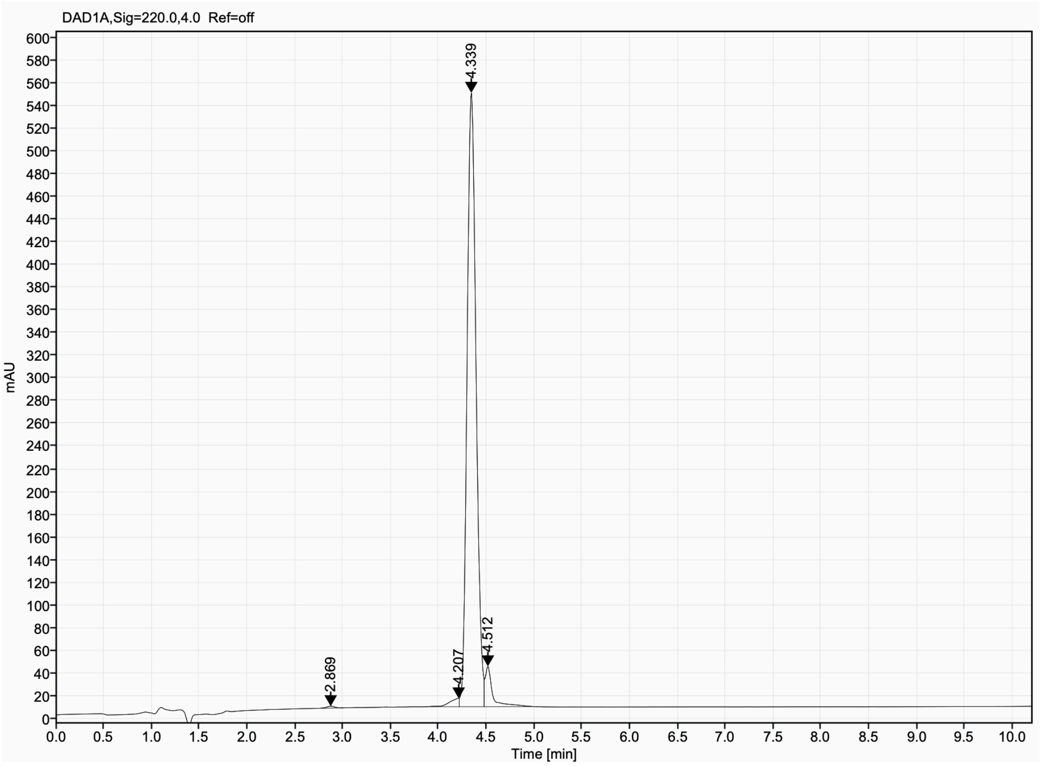

Analytical UPLC and mass spectrum of RBB_D6
Sequence: cyclo[FSKYEPVDDSYPEGIRKV]

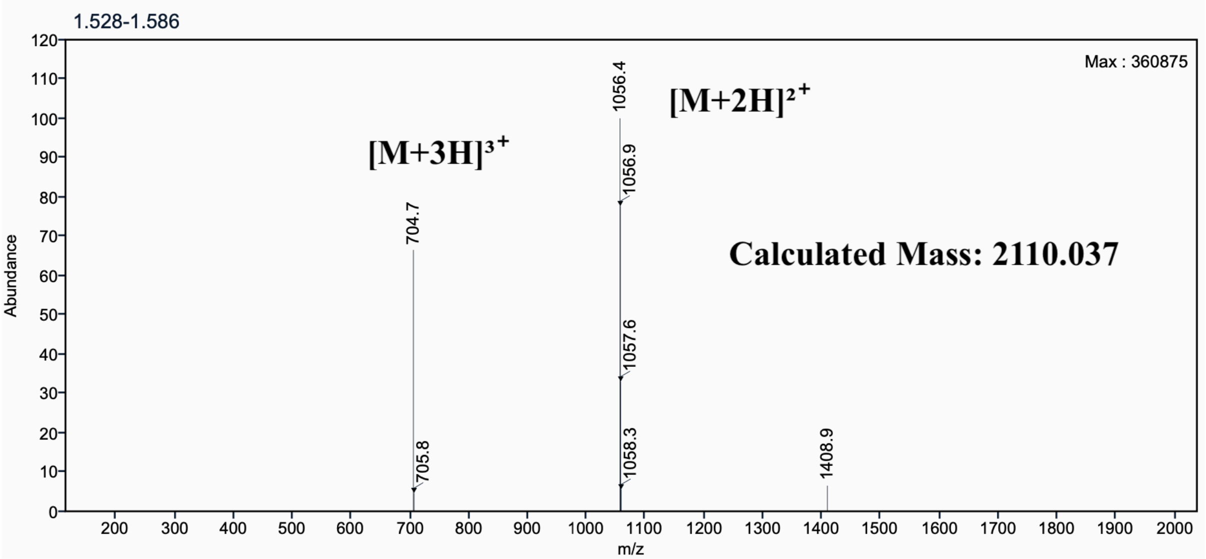

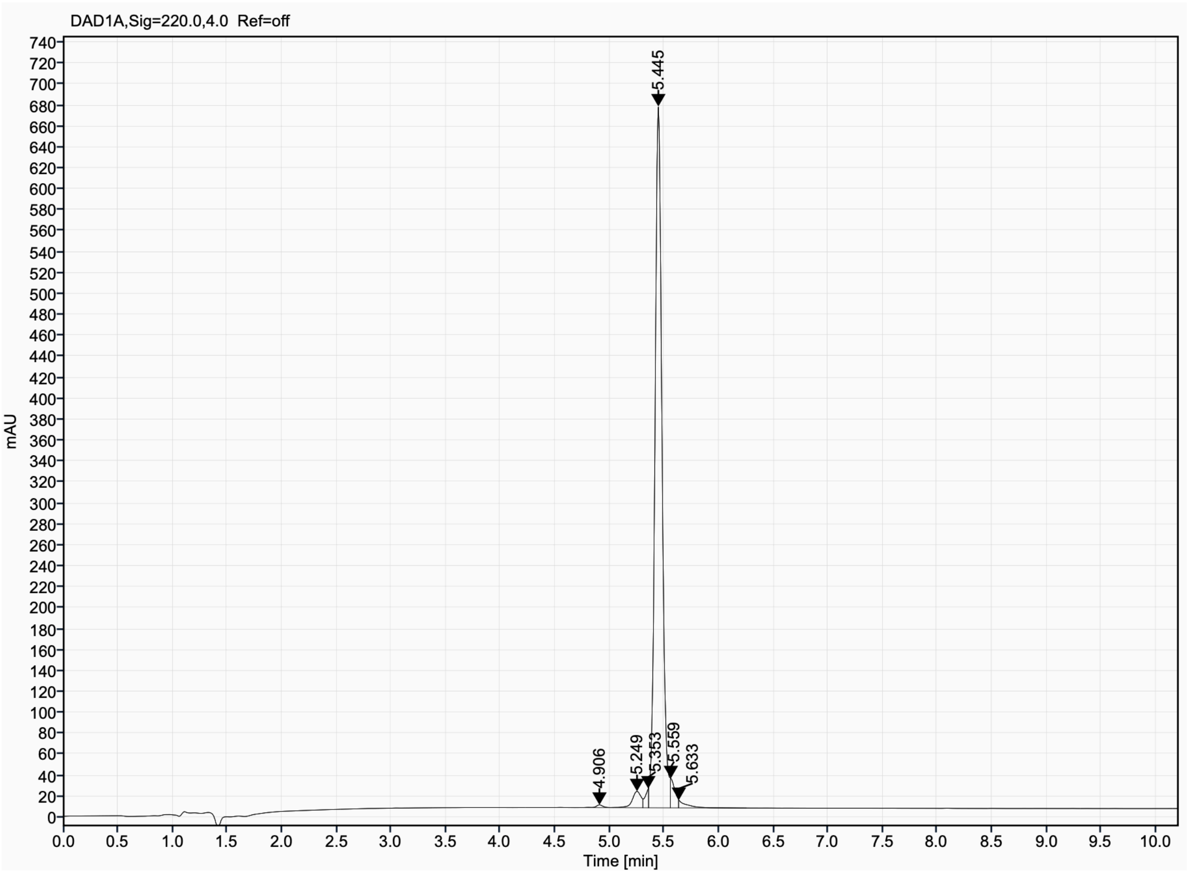

Analytical UPLC and mass spectrum of RBB_D7
Sequence: cyclo[FEKYDPMDESYPENIRKV]

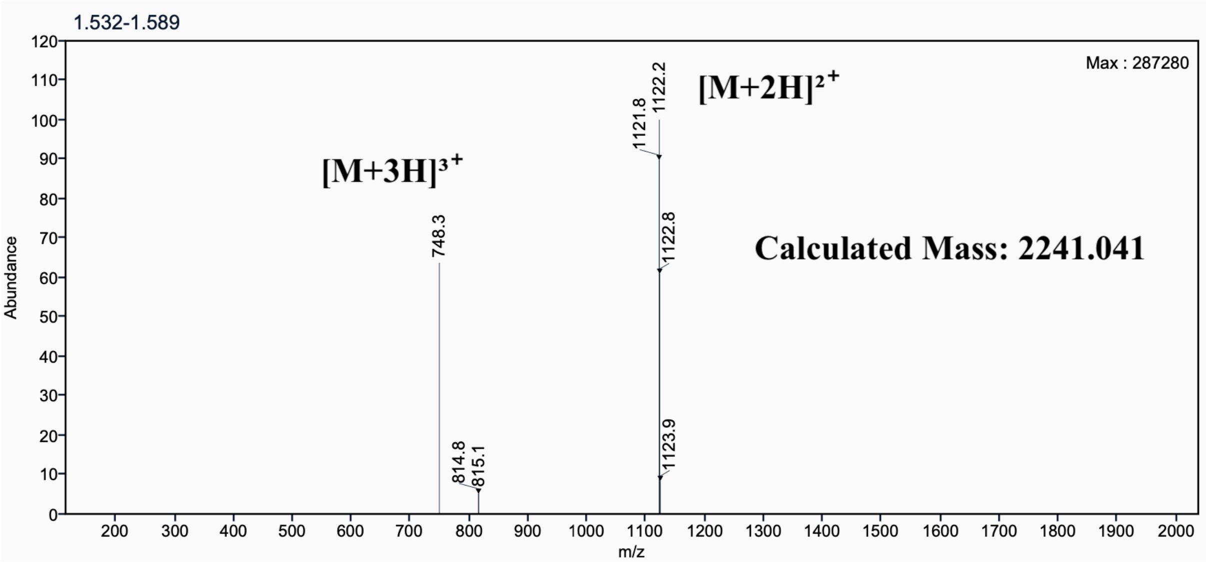

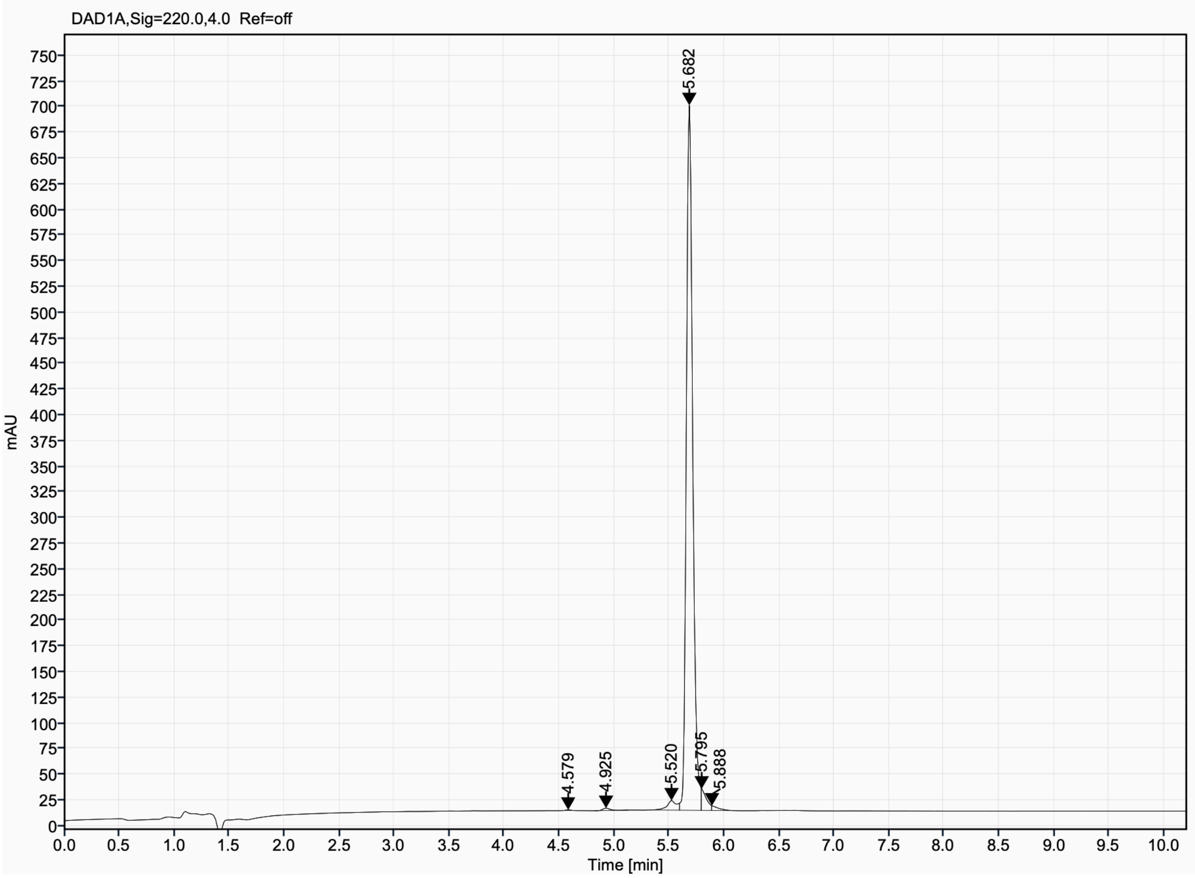

Analytical UPLC and mass spectrum of RBB_D8
Sequence: cyclo[VEGNDKGSDIYEFIRDK]

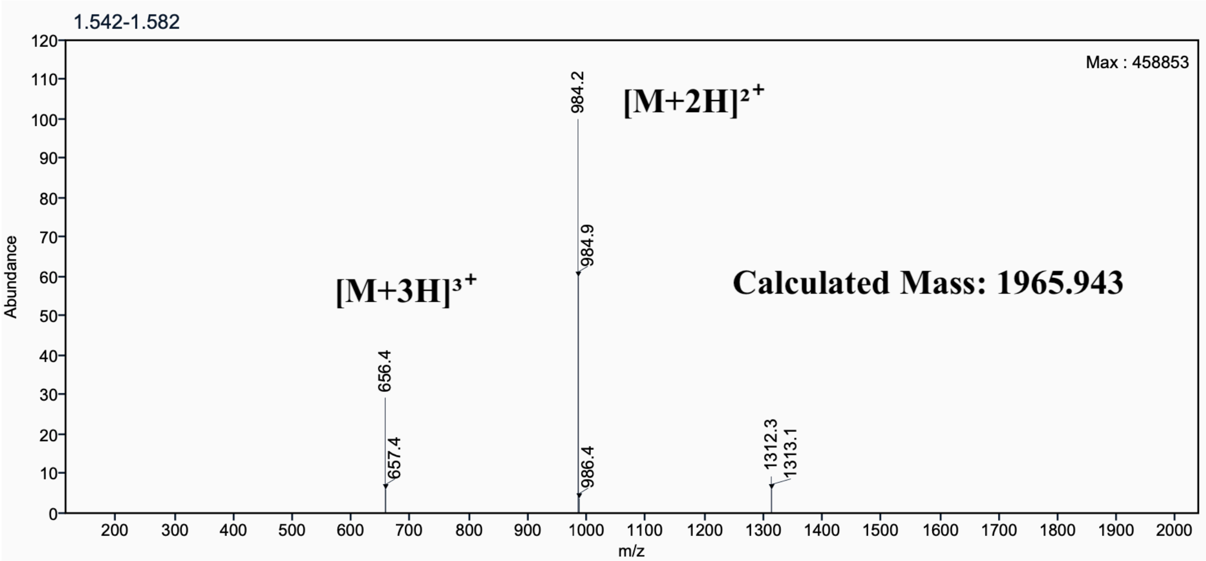

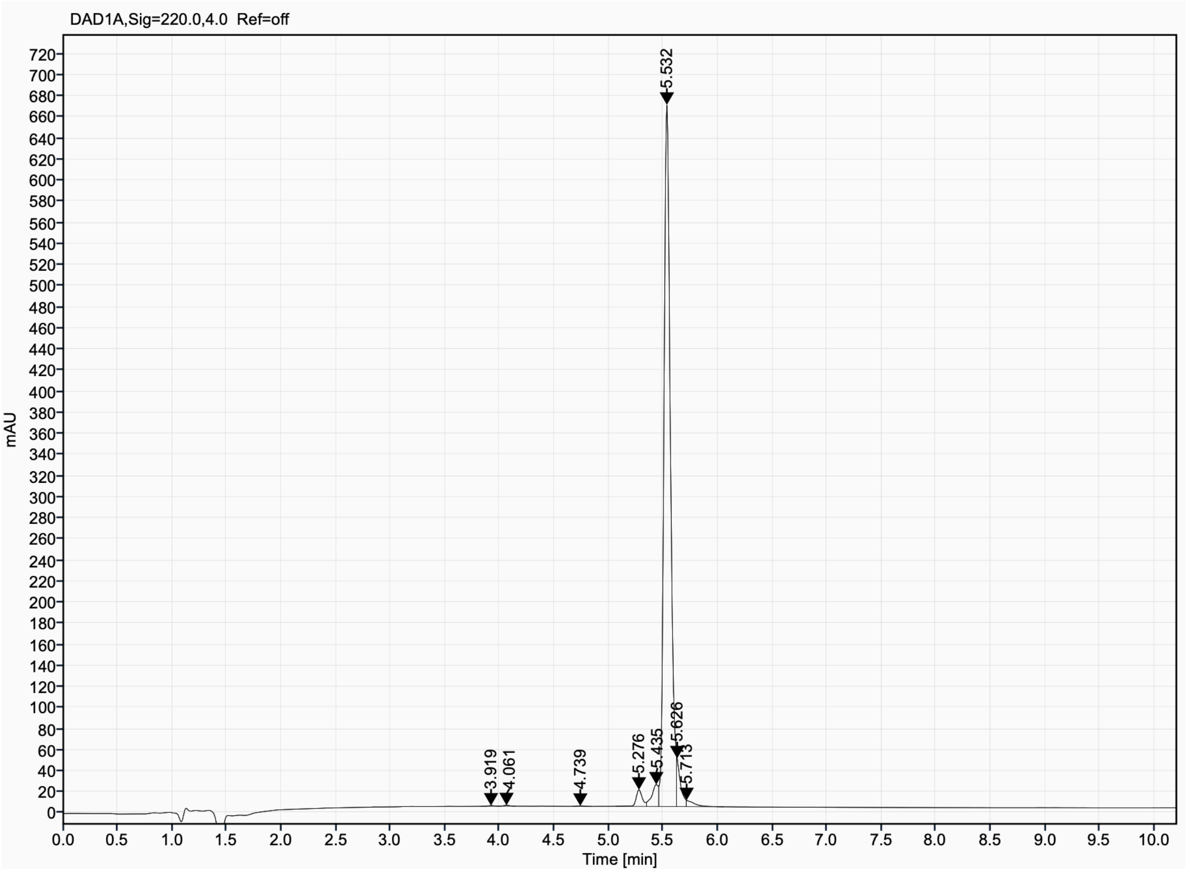

Analytical UPLC and mass spectrum of RBB_D10
Sequence: cyclo[KLFGPDPYLPENVQ]

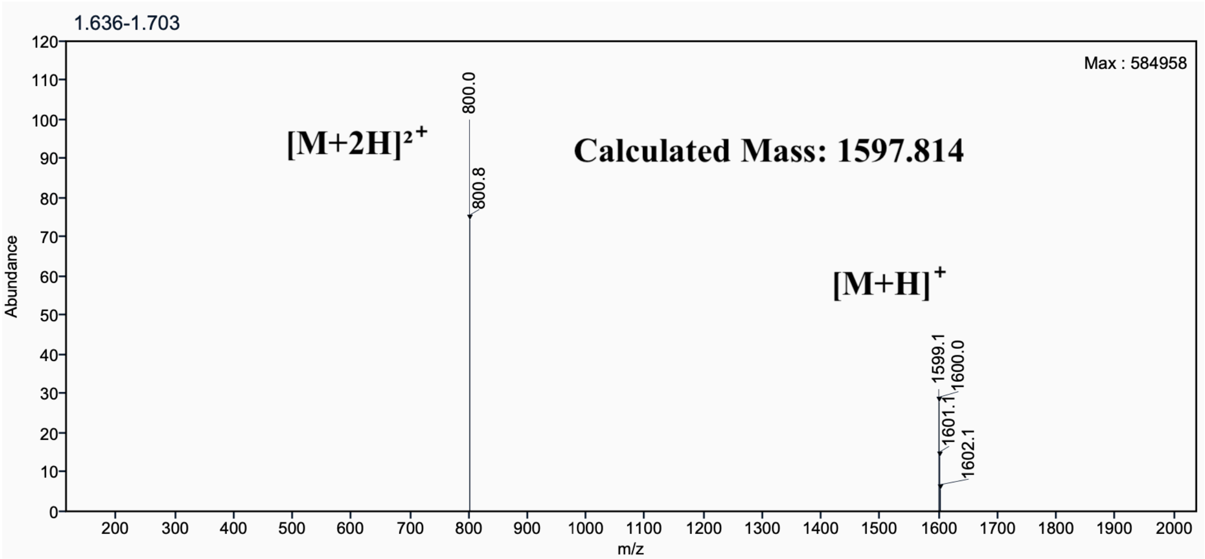

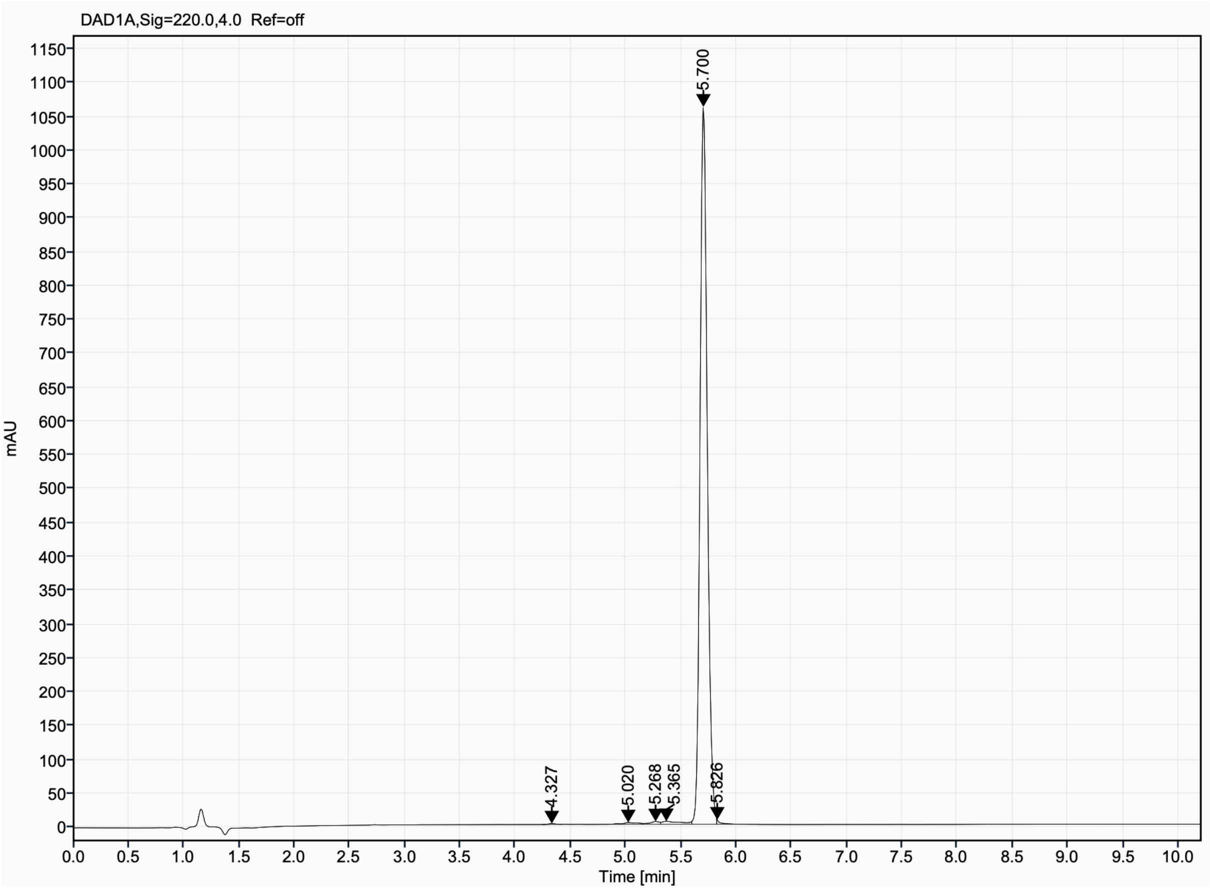

Analytical UPLC and mass spectrum of RBB_D14
Sequence: cyclo[SVAKEIAEWIGIPSKVPP]

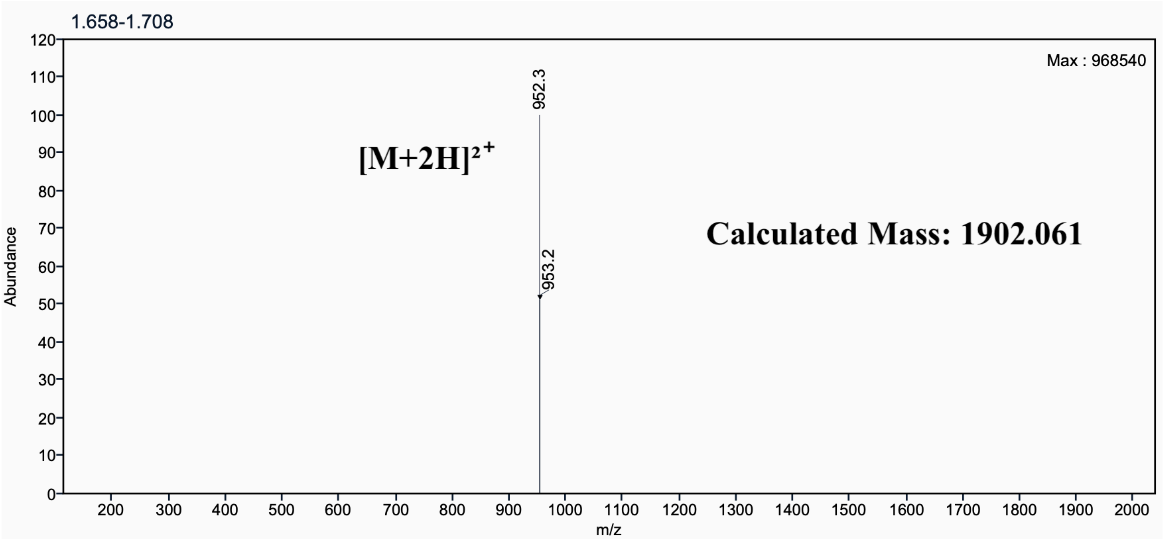

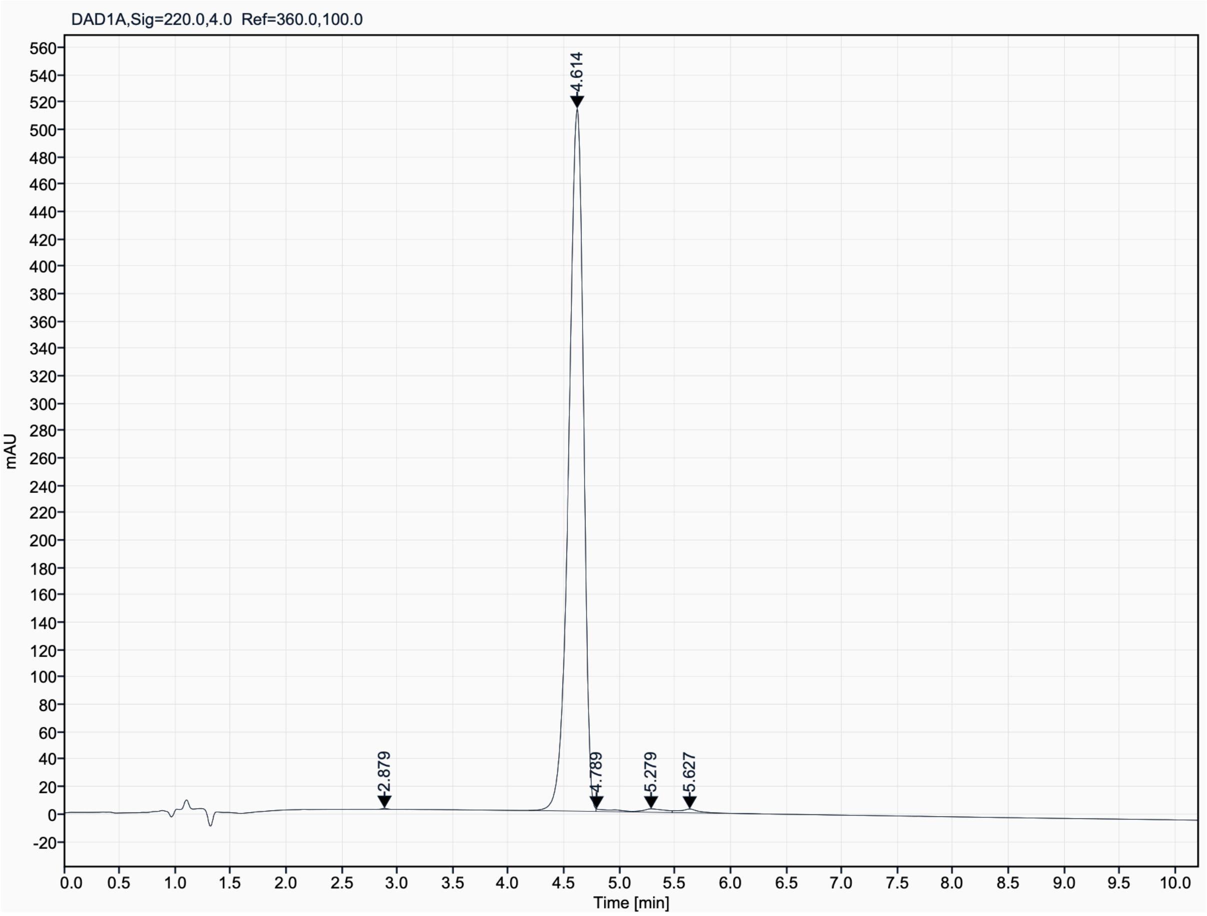

Analytical UPLC and mass spectrum of RBB_D15
Sequence: cyclo[SLAKEIADWIGIPSSVPP]

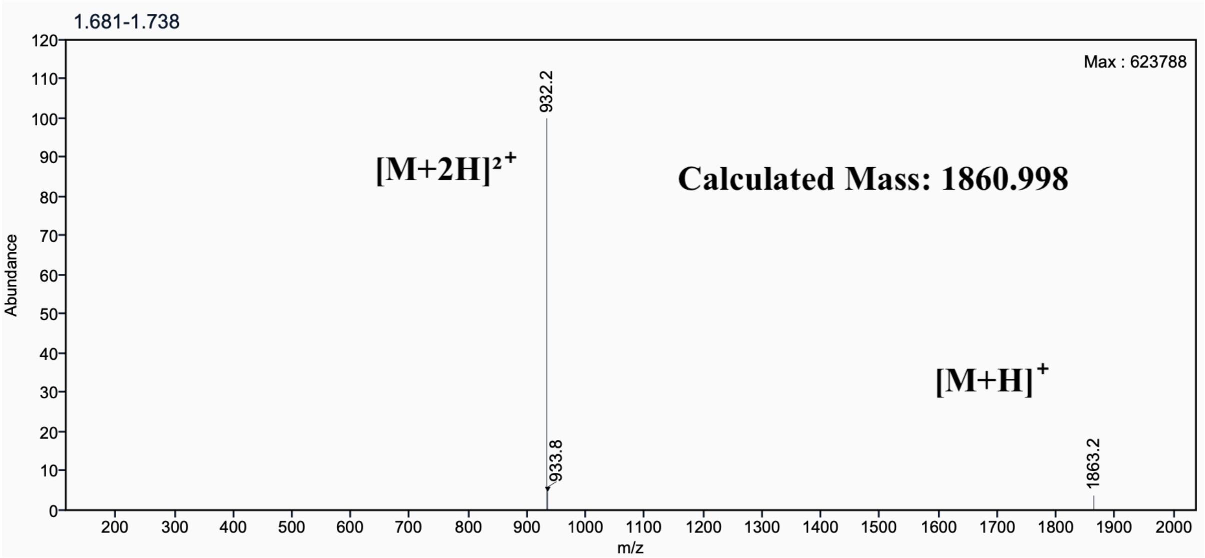

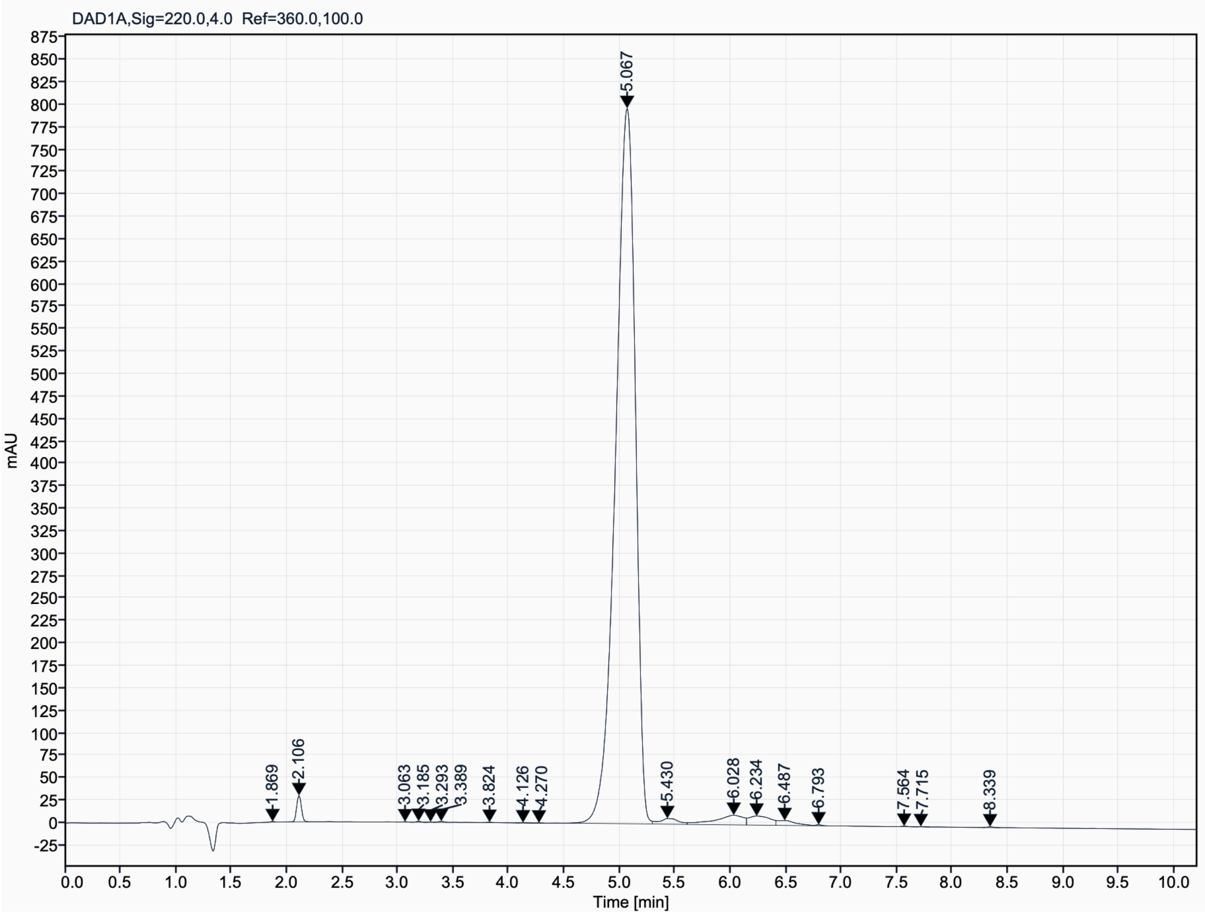
